# Brominated Lipid Probes Expose Structural Asymmetries in Constricted Membranes

**DOI:** 10.1101/2021.08.24.457428

**Authors:** Frank R. Moss, James Lincoff, Maxwell Tucker, Arshad Mohammed, Michael Grabe, Adam Frost

**Author notes:** Corresponding author. (A.F.) and (M.G.). These authors contributed equally to this work (Frank R. Moss, James Lincoff).

## Abstract

Lipids in biological membranes are thought to be functionally organized, but few experimental tools can probe nanoscale membrane structure. Using brominated lipids as contrast probes for cryo-EM and a model ESCRT-III membrane remodeling system, we observed leaflet-level and protein-localized lipid structural patterns within highly constricted and thinned membrane nanotubes. These nanotubes differed markedly from protein-free, flat bilayers in leaflet thickness, lipid diffusion rates, and lipid compositional and conformational asymmetries. Simulations and cryo-EM imaging of brominated SDPC showed how a pair of phenylalanine residues scored the outer leaflet with a helical hydrophobic defect where polyunsaturated docosahexaenoyl (DHA) tails accumulated at the bilayer surface. Combining cryo-EM of halogenated lipids with molecular dynamics thus enables new characterizations of the composition and structure of membranes on molecular length scales.

**One-Sentence Summary:** Cryo-EM imaging of halogenated lipids and MD simulations provide molecular-scale insight into constricted bilayer structure.

## Main Text

Cells utilize molecular machines to form and remodel their membrane-defined compartments’ compositions, shapes, and connections. The regulated activity of these membrane remodeling machines drives processes like vesicular traffic and organelle homeostasis. However, the precise mechanisms by which proteins generate mechanical force to catalyze membrane fission, fusion, and shape changes remain elusive. Further, the structural evolution of lipids and lipid bilayers during these processes are challenging to study, as well as the energetic contributions of individual lipid species to membrane remodeling. A better understanding of membrane and lipid structure during remodeling is crucial to understanding both the mechanisms of membrane remodeling and, more broadly, the interactions between proteins and membranes.

Molecular-scale insights into the lipid-leaflet, lipid-lipid, and lipid-protein dynamics that generate extreme membrane curvature could clarify the mechanisms of membrane remodeling. A thorough understanding of membrane mechanics will account for lipid flip/flop, spontaneous lipid curvature, bending rigidity, line tensions, and protein-generated forces including amino acid insertions, lateral pressure from protein crowding, shearing forces, and lipid-protein “friction” (*1–10*). The relative contributions of each of these processes to membrane remodeling generally, and to membrane constriction in the present case, are challenging to measure experimentally or explain in structural terms. We need new measures of how membrane structure, composition, and protein-generated forces influence membrane properties *in vitro* and *in vivo*.

To date, cryo-EM reconstructions of membrane-bound proteins have generated insights into the machines that shape membranes (*11–19*). However, these studies have generally failed to reveal molecular-level information about the membrane itself because most lipids display nearly indistinguishable electron scattering (*20, 21*). Fluid bilayers, moreover, are generally thought to lack a structured pattern at the nanoscale that is recoverable by image averaging. Therefore, it is typically impossible to distinguish individual lipid species, even approximately, except in high-resolution structures where isolated, static lipids are resolved bound to transmembrane proteins (*22–25*). To overcome this shortcoming, here we describe halogen-labeled lipid probes as contrast-enhancing agents for cryo-EM. We use a model system consisting of human ESCRT-III proteins CHMP1B and IST1 to remodel vesicles containing lipid probes into lipid nanotubes with extremely high curvature (*26, 27*). Using labeled lipids, ESCRT-III proteins, cryo-EM, and molecular dynamics simulations, we characterize how ensembles of lipid-protein interactions, lipid conformational changes, and the resulting stabilization of strong membrane asymmetries drive membrane constriction and thinning. Leaflet and lipid shape change together with specific interactions between lipids and the ESCRT-III proteins, stabilize membrane curvature in this snapshot of a membrane on the verge of scission.

## Results

### The human ESCRT-III protein CHMP1B α1 induces a furrow in the bilayer outer leaflet

When exposed to model membranes in vitro, ESCRT-III proteins can shape membranes into structures with zero, positive, or negative curvature (*27–34*). We recently demonstrated how a pair of ESCRT-III proteins, CHMP1B and IST1, assemble helical filaments that act in sequence to remodel membranes into high curvature nanotubes with an inner diameter of only ∼40 Å— nearly to the point of fission (*27, 35*). This system provides an opportunity for probing how peripheral proteins engage distinct lipid species to pattern membrane properties more generally.

Previously, we observed that helix α1 of CHMP1B appeared to “dimple” the outer surface of highly constricted membrane nanotubes, but the properties of this deformation were poorly defined (Fig. 1A-B, E) (*27*). To explore the membrane’s structure in more detail, we performed coarse-grained molecular dynamics (CG-MD) simulations of the membrane tubule in the presence of the CHMP1B-IST1 copolymer using the Martini 2.2 CG force field (Fig. 1C-D) (*36*). We used the previously optimized mixture of SDPC, POPS, PIP2, and CHOL for the simulation and subsequent cryo-EM experiments (see Fig. S1-10 for chemical structures) (*27*). The simulations revealed CHMP1B side chains sterically displacing phospholipid headgroups and exposing the hydrophobic core of the membrane in an apparent “scoring” of the surface—rather than an elastic dimpling of a continuous headgroup surface as we previously thought (Fig. 1F).

**Fig. 1.**
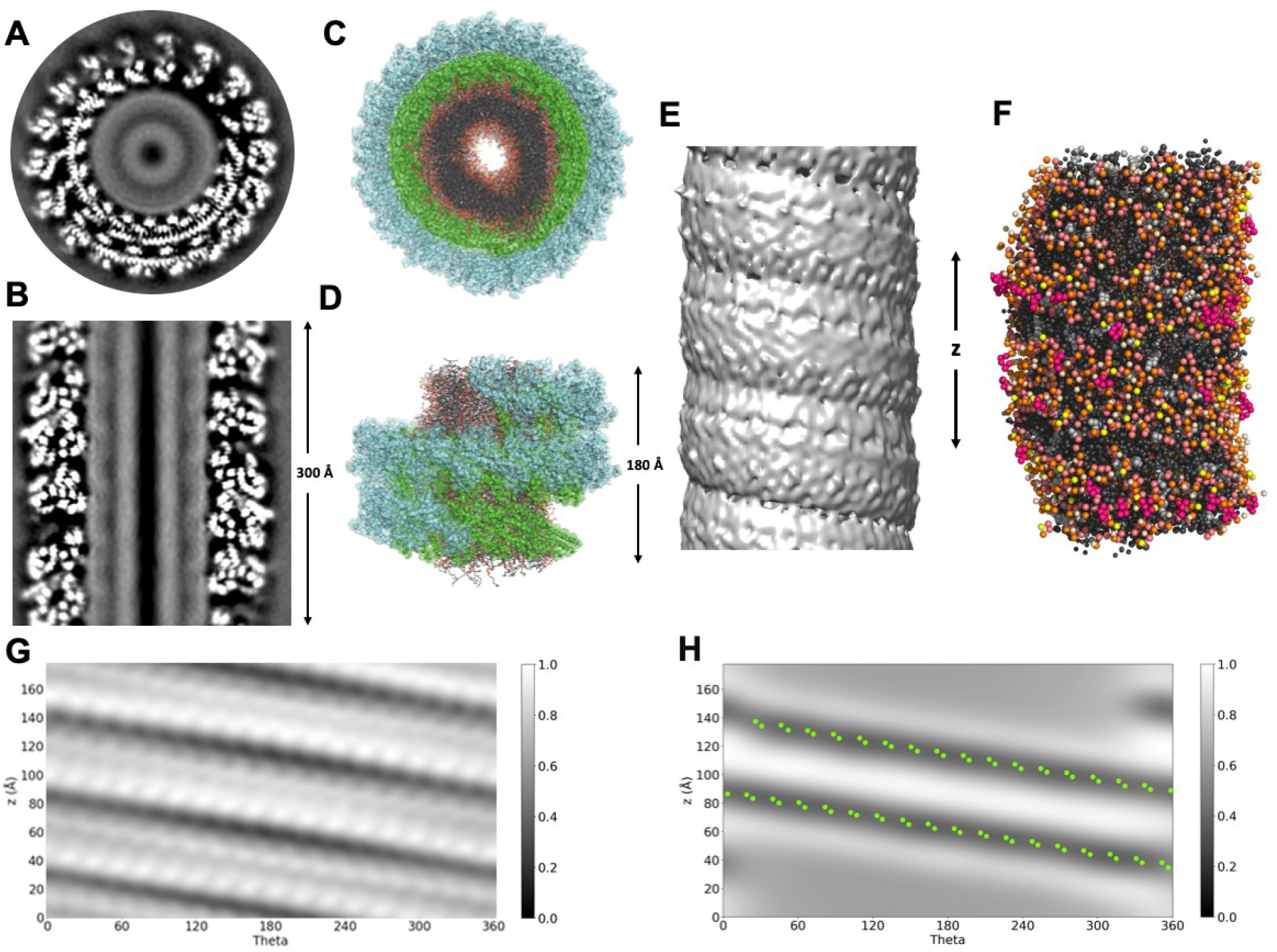
Cryo-EM and CG-MD structures of membrane deformations induced by CHMP1B and IST1. A) Grey-scale top-down and B) side views of central slices of the cryo-EM density for the CHMP1B/IST1 copolymer bound to a cylindrical lipid bilayer comprising the lipids shown in composition #1, Table 1. C) Top-down and D) side views of an equilibrated tubule within the protein coat from CG-MD simulations, in all-atom representation. CHMP1B in green, IST1 in cyan, cholesterol in light grey, lipid tails in dark grey, phospholipid glycerol moieties in salmon, phospholipid phosphates in orange, SDPC headgroups in beige, POPS headgroups in yellow, and PIP2 headgroups in pink. E) Side view of the lipid bilayer nanotube’s cryo-EM density with protein masked. F) Side view of the simulated CG-MD membrane tubule with protein hidden shows two bands where headgroups are displaced, and lipid tails are exposed. Color scheme is in C and D. G) EM density of the outer leaflet of the bilayer visualized in cylindrical coordinates. The periodic footprint of CHMP1B is apparent at this depth. H) Two-dimensional headgroup density of outer leaflet lipids-only averaged over all CG-MD simulations (headgroup exclusion zone, dark bands) with F9 and F13 sidechain locations (green dots).

To aid visualization of the headgroup exclusion zone, we transformed the protein and lipid positions into cylindrical coordinates and plotted the mean density of headgroups in the outer leaflet (Fig. 1H) (*36*). The headgroup-excluding furrow aligned with the bulky hydrophobic residues F9 and F13, which protrude from helix α1 on CHMP1B (green dots in Fig. 1H). Notably, despite the 23 Å distance between these pairs of residues on adjacent CHMP1B subunits, the furrow forms a continuous stripe in the membrane outer leaflet. This aspect of the simulated structures is in agreement with the experimental structure determined by cryo-EM. The EM-derived density map also revealed a furrow of low Coulombic potential aligned with the stripe of F9 and F13 positions along the CHMP1B helical filament (Fig. 1G). Together, the EM maps and CG-MD results suggested that the observed deformation constitutes a stable lipid packing defect due to headgroup displacement by these hydrophobic amino acids of CHMP1B and concomitant exposure of the lipid tails underneath (Fig. 1G-H). Line tension associated with the hydrophobic defect may energetically drive the defects to coalesce into a continuous, helical stripe. Because such an asymmetry in the membrane should alter its mechanical properties, influencing the energetic barriers associated with curvature or fission, we sought to further characterize its structure.

### Specific lipids both enrich at the Phe contact site and change shape with curvature stress

We next examined the predicted molecular structure and dynamics of the lipids throughout the bilayer. This analysis revealed pronounced asymmetries in lipid shape correlated with leaflet localization and proximity to the F9, F13 contact site (Fig. 2A-D). The lipid SDPC, which has a polyunsaturated (22:6) tail at the sn-2 position and a fully saturated (18:0) tail at sn-1, showed the strongest asymmetries. While the saturated sn-1 tail typically remained aligned with the membrane normal throughout both leaflets (Fig. 2A-D), the polyunsaturated sn-2 tail of outer leaflet lipids bent backward and radially outward toward the headgroup region at the F9, F13 contact site (Fig. 2A, C). This shape transformation partially filled the space vacated by the displaced headgroups and enabled the tail to interact directly with the hydrophobic sidechains of F9 and F13. Away from the exclusion zone, the polyunsaturated tail adopted a more typical conformation (Fig. 2B, D).

**Fig. 2.**
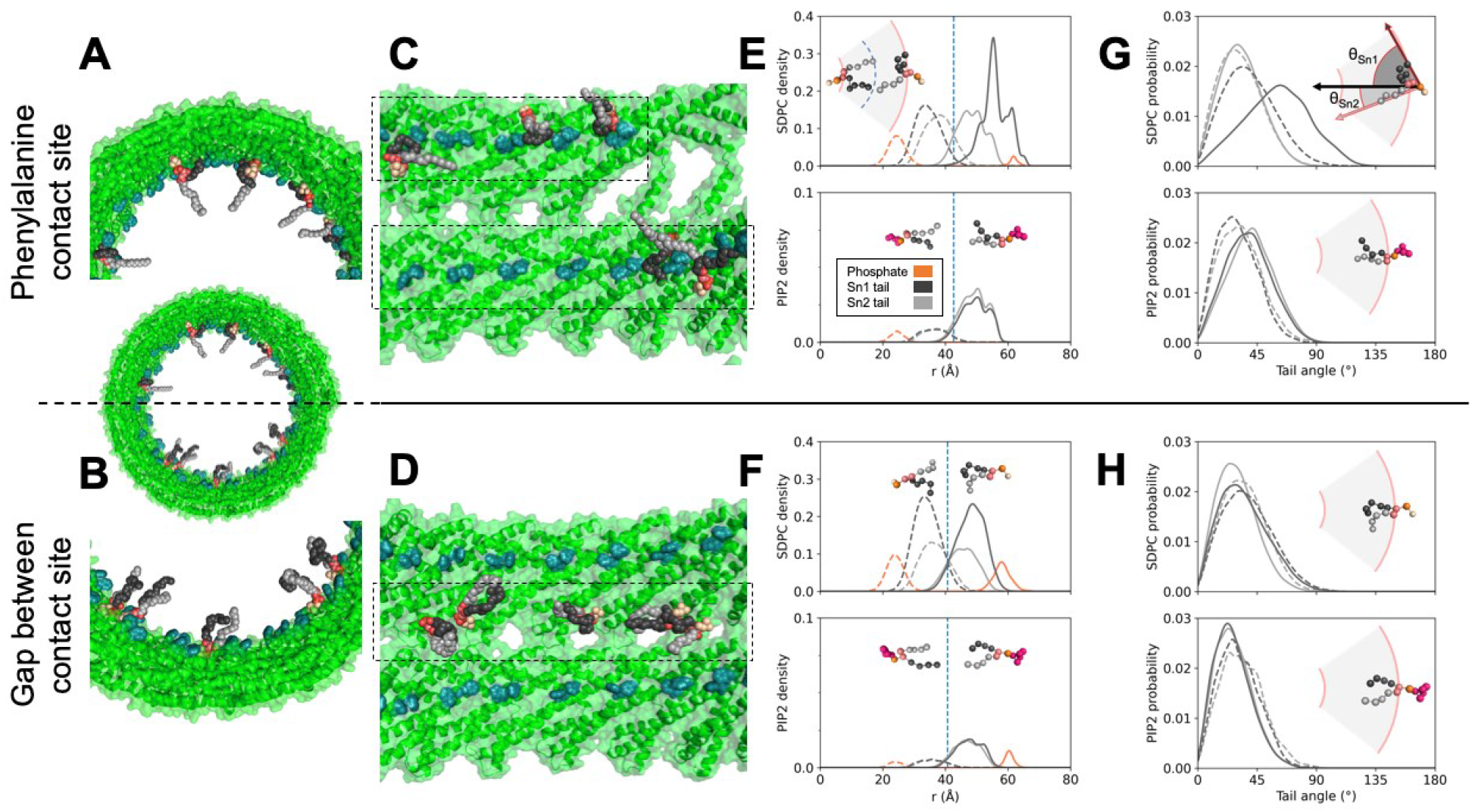
Analysis of lipid conformation and geometry. A) Representative SDPC molecules from a CG-MD snapshot backmapped to all-atom representation with sn-2 tail (black) interacting with F9 and F13 (teal) of CHMP1B (green), while the sn-1 tail (grey) is normally oriented. SDPC headgroup carbons shown in beige, and glycerols in salmon. B) Representative backmapped SDPC molecules from a CG-MD snapshot in the gap between CHMP1B α1 helices. C) Side view of lipids in A, looking from the tubule center toward CHMP1B. Dashed boxes highlight the spiral of F9 and F13. D) Side view of lipids in B. Dashed box highlights the gap zone between stripes of F9 and F13. E) Radial densities of lipid components at bilayer contact sites for SDPC (top) and PIP2 (bottom). Inner leaflet densities in dashed lines, outer leaflet densities in solid lines. Color coding highlights phosphate bead (orange), all sn-1 tail beads (grey), and all sn-2 tail beads (black). Vertical dashed lines denote the local bilayer midplane. F) Radial densities of lipid components at gaps for SDPC (top) and PIP2 (bottom). Note the different y-axis scale for PIP2 in E and F. G) Probability density of lipid tail angles to the bilayer normal for SDPC (top) and PIP2 (bottom) at the Phe contact site. Black lines are sn-2 tails, grey lines are sn-1 tails, solid lines are outer leaflet, and dashed lines are inner leaflet. H) Probability density of lipid tail angles to the bilayer normal at the gaps. In E-H, example CG conformations use the same color scheme as A-D, with PIP2 headgroups in pink to contrast with SDPC headgroups (beige).

To quantify lipid conformation, we calculated the lipid tilt angles and radial densities of the phosphate and tail beads of all phospholipids adjacent to and in between the Phe contact sites for both leaflets (Fig. 2E-H and Fig. S11). We defined the “Phe contact site” as the 30 ° wedge centered through the angular position of F9 and F13, while the “gaps between the contact sites” were defined as the 30 ° wedge centered 180 ° from the contact sites (Fig. S12) (*36*). Consistent with our anecdotal observations of tail “backflipping”, the sn-2 tail of SDPC was highly deformed in the outer leaflet at the Phe contact sites with its mean location shifted radially outward and a substantial density beyond both the bilayer phosphate peak, typically the membrane’s outer surface, and the Phe sidechains (panel e, *r* = 60 Å). On the other hand, in the inner leaflet, both tails are largely unperturbed throughout (Fig. 2E-F). The other lipid tails showed more minor degrees of deformation at the Phe contact sites and normal bilayer behavior in the gaps between furrows.

The tilt angle analysis revealed corresponding asymmetries (Fig. 2G-H). In both leaflets, SDPC tails adopted typical values of ∼30 ° with respect to the bilayer normal—except for the polyunsaturated sn-2 tail at the furrows, which displayed pronounced tilting with a mean value of 60 ° and some extreme conformations greater than 90 ° where the tail bead resided at the membrane surface beyond the radial position of F9 and F13. The other two phospholipid species, POPS and PIP2, which have both saturated and monounsaturated lipid tails, did not exhibit a similarly strong location-dependent perturbation (Fig. S11).

These observations are consistent with the notion that polyunsaturated lipid tails enable membrane shape plasticity under curvature stress, which prior work has suggested may be a general principle of bilayer deformation (*4, 27, 37–42*). Indeed, the polyunsaturated tail of SDPC is known to be highly flexible. This property could facilitate membrane bending if the lipid changes its mean shape in response to curvature stress (*43–46*). Consistent with this hypothesis, replacing SDPC with monounsaturated POPC precluded high-curvature tubule formation (Fig. S13). Specifically, CHMP1B and IST1 were unable to constrict POPC-containing liposomes to high curvature (lipid composition 3), as assessed by negative stain TEM and cryo-EM (Fig. S13) (*36*).

We next sought direct experimental validation of SDPC’s localization in the bilayer and whether it undergoes such dramatic shape transformations. To detect individual lipid species using cryo-EM, we synthesized analogs of each lipid with bromine atoms along the unsaturated lipid tail lengths. Due to their more massive nuclei, bromine atoms generate contrast through enhanced electron scattering. The brominated and unbrominated lipids displayed similar bilayer phase and Langmuir monolayer behaviors, indicating comparable lipid packing and fluidity (Fig. S14). Additionally, in bilayers, SDPC-Br and POPS-Br are fluid at room temperature and macroscopically homogeneous, consistent with previous studies (*47, 48*). Importantly, CHMP1B and IST1 remodeled brominated vesicles into narrow nanotubes yielding structures indistinguishable at ∼3 Å resolution from vesicles prepared with native lipids (Fig. 3 and Fig. S13). Together, these results indicate that halogenation does not meaningfully perturb the properties of these lipids or the bilayers formed from them. We collected high-resolution data and achieved high-resolution reconstructions with vesicles in which each lipid was replaced by its halogenated analog (Table 1, lipid compositions 10-14) or different fractions of SDPC were brominated (Table 1, lipid compositions 10, 15, 16).

**Table 1.** Lipid mixtures used in membrane remodeling assays. All amounts are mole percentages.

| Number | Composition | Components |
| --- | --- | --- |
| 1 | 58.2 : 18 : 17.5 : 6.3 | SDPC : CHOL : POPS : PIP2 |
| 2 | 62.1 : 18.7 : 19.2 : 0 | SDPC : CHOL : POPS : PIP2 |
| 3 | 58.2 : 18 : 17.5 : 6.3 | POPC : CHOL : POPS : PIP2 |
| 4 | 71 : 0 : 21.3 : 7.7 | SDPC : CHOL : POPS : PIP2 |
| 5 | 63.9 : 10 : 19.2 : 6.9 | SDPC : CHOL : POPS : PIP2 |
| 6 | 56.8 : 20 : 17.1 : 6.1 | SDPC : CHOL : POPS : PIP2 |
| 7 | 53.2 : 30 : 16 : 5.8 | SDPC : CHOL : POPS : PIP2 |
| 8 | 42.6 : 40 : 12.8 : 4.6 | SDPC : CHOL : POPS : PIP2 |
| 9 | 29.1 : 50 : 8.8 : 3.1 | SDPC : CHOL : POPS : PIP2 |
| 10 | 58.2 : 18 : 17.5 : 6.3 | SDPC-Br : CHOL : POPS : PIP2 |
| 11 | 58.2 : 18 : 17.5 : 6.3 | SDPC : CHOL-Br : POPS : PIP2 |
| 12 | 58.2 : 18 : 17.5 : 6.3 | SDPC : CHOL : POPS-Br : PIP2 |
| 13 | 58.2 : 18 : 17.5 : 6.3 | SDPC : CHOL : POPS : PIP2-Br |
| 14 | 58.2 : 18 : 17.5 : 6.3 | SDPC : CHOL-I : POPS : PIP2 |
| 15 | 29.1 : 29.1 : 18 : 17.5 : 6.3 | SDPC-Br : SDPC : CHOL : POPS : PIP2 |
| 16 | 14.6 : 43.6 : 18 : 17.5 : 6.3 | SDPC-Br : SDPC : CHOL : POPS : PIP2 |
| 17 | 26 : 22 : 32 : 20 | SDPC : CHOL : POPS : PIP2 |

**Fig. 3.**
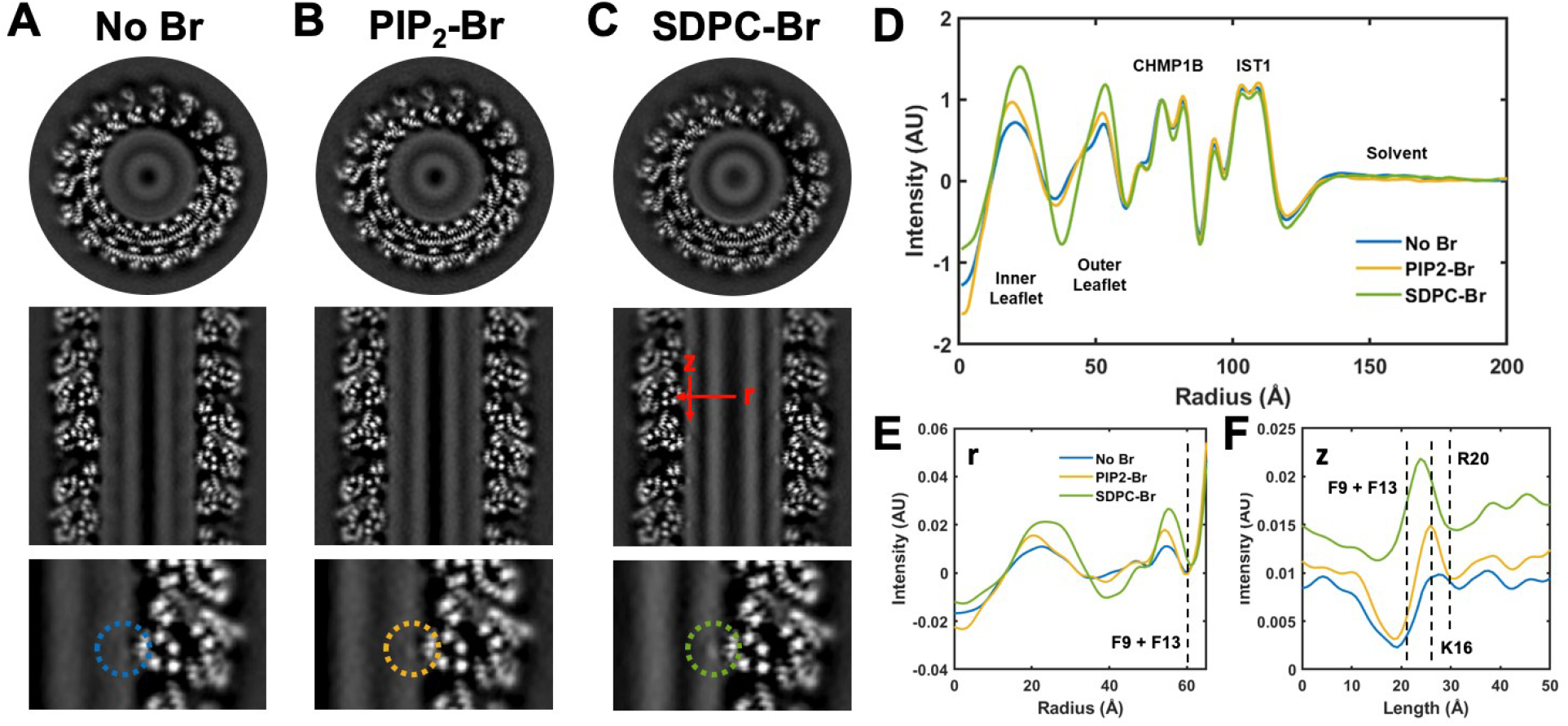
**CryoEM reconstructions of unlabeled and lipid-brominated membrane-bound CHMP1B/IST1 filaments**. (A-C) The top panel shows top-down views of central slices of the reconstructions, the middle panel shows side views of central slices of the reconstructions, and the bottom panel shows zoomed-in views of the region where F9 and F13 of CHMP1B α1 contact the membrane. Dashed circles highlight the protein-membrane contact site. D) Radial averages (around the filament axis) of the reconstructions in A-C show extra intensity from the bromine labels in the lipid bilayer, while the protein structure remains unaffected. E) Line profile along the filament radius at Phe contact site (r, see C middle panel). F) Line profile along the filament axis centered around Phe contact site (z, see C middle panel). The approximate positions of relevant CHMP1B residues along the line scans are indicated by dashed lines.

For this analysis, the pixel value distributions were normalized to the CHMP1B intensity from radial averages to compare each structure semi-quantitatively. We first compared horizontal and vertical slices through each filament and observed differences in the intensities of the bilayer leaflets with brominated lipids. In particular, the SDPC-Br reconstruction showed intense spots of enhanced density adjacent to F9 and F13, consistent with SDPC-Br enrichment at the site (Fig. 3C) and corroborating the CG-MD simulation data showing that lipid tails reached the surface of the bilayer. The other brominated lipids, which are monounsaturated, also accumulated at the membrane-protein contact site, but to a smaller extent. Further, we resolved differences in the positions of different lipid species relative to the protein contact sites, with the brominated SDPC tail in closer proximity to F9 and F13 both radially and axially than brominated PIP2 and POPS tails (Fig. 3E-F). Intensities indicated that these latter two anionic lipids more closely approached K16 and R20, two basic residues previously identified as critical for CHMP1B membrane binding to membranes (*27*).

The altered SDPC tail conformation where the bilayer contacts CHMP1B residues F9 and F13 suggests at least two hypotheses. First, lipid headgroup displacement by F9 and F13 creates lipid packing defects and curvature stress that is relieved when lipid tails fill in the packing defects. Second, hydrophobic interactions drive the lipid tails to interact with the nonpolar Phe side chain. These ideas, the first focused on steric packing and the second focused on chemical interactions between lipids and amino acids, are not mutually exclusive. We generated a series of CHMP1B double mutants, F9X + F13X (where X = A, E, or L), to test the relative contributions of each possibility.

These CHMP1B double mutants still bound to and remodeled native and brominated lipid bilayers, forming membrane-bound copolymers with IST1 indistinguishable from the wild-type CHMP1B/IST1 copolymer in helical symmetry (Fig S15). Interestingly, F9 and F13 substitution by small nonpolar Ala sidechains eliminated SDPC-Br enrichment at CHMP1B contact sites (in lipid composition 10). SDPC also failed to concentrate in the Phe to Glu double mutant. Interestingly, in both the Phe to Ala and the Phe to Glu double mutants, the protein appeared to induce an elastic deformation of the bilayer without opening an appreciable hydrophobic defect. Simulations of the Phe to Ala double mutant also showed reduced SDPC tail enrichment and backflipping, as well as increased headgroup density shifted radially inward, at CHMP1B contact sites, consistent with an elastic deformation (Fig. S16). By contrast, SDPC-Br was observed to accumulate at contact sites with large hydrophobic Leu side chains to a similar but lesser degree than the wild-type protein. Removal of the protein coat in simulations resulted in dramatic reduction of SDPC backflipping and recovered mostly normal bilayer structure and lipid conformation (Fig. S17). Together, these observations are consistent with chemical interactions between greasy side chains and polyunsaturated tails stabilizing the backflipped lipid tail conformations (*36*).

### Constriction leads to anisotropic inner leaflet thinning

Yesylevskyy and coworkers predicted that a highly curved bilayer’s inner, concave leaflet would be thinner than the outer, convex leaflet based on atomistic simulations (*49*). We tested this hypothesis by examining the Coulombic potential (electron scattering) profile from the unbrominated sample and fitting the profile with three Gaussian functions using least-squares (*36, 50*). As shown in Fig. 4A and 4C, the inner concave leaflet is 3.6 Å thinner than the outer convex leaflet, in agreement with Yesylevskyy et al. and our own CG-MD simulations of the same system (Fig. 4B and 4C). Differences in lipid conformations, with the tails of inner leaflet lipids adopting more splayed configurations than those of the outer leaflet, may drive this asymmetry (Fig. 2G, H). This lipid-splay effect is also likely the origin of the overall bilayer thinning as a function of curvature we previously reported (*27*).

**Fig. 4.**
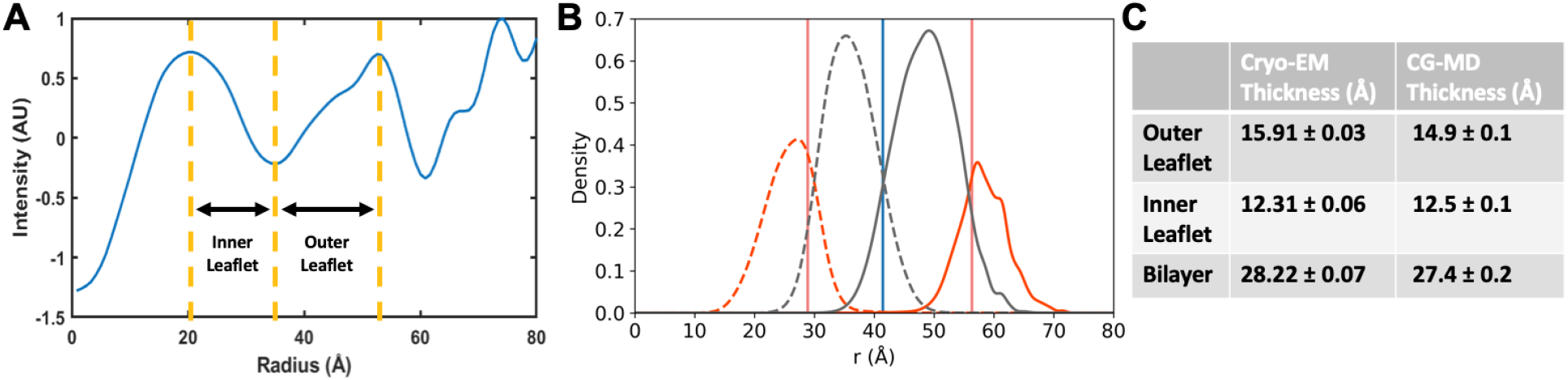
Bilayer and leaflet thicknesses of a membrane nanotube from simulation and experiment. A) Radial profiles from cryo-EM reconstructions provide estimates of inner and outer leaflet thickness values by computing distances from each leaflet peak to the aliphatic trough. B) Simulated hydrophobic (gray), phosphate (orange), and glycerol (pink dashed line) lipid components for the inner leaflet (dashed lines) and outer leaflet (solid lines) are plotted. C) Summary of calculated leaflet and bilayer thicknesses from cryo-EM and CG-MD. Uncertainty is reported as the standard deviation between independent measurements from each half map for cryo-EM, and as the standard deviation between ten simulation replicates for CG-MD.

### The constricted membrane is compositionally asymmetric and heterogeneous

Both MD simulations and cryo-EM reconstructions suggested the bilayer composition is not homogeneous, with difference at the F9, F13 contact site versus the gap between the contact sites. By integrating the areas under the curve for each leaflet from bromine-labeled reconstructions normalized to the unlabeled reconstruction, we estimated the composition of each leaflet in these two regions (Fig. 5A-B) (*36*). As shown in Fig. 5C, the tubule leaflet composition appears to be quite different from the starting bulk values (Table 1, composition 1). Anionic lipids are enriched in the filament at the expense of SDPC, likely due to the highly cationic luminal surface of the protein coat (Fig. S13, S18). We repeated the CG-MD simulations with the composition estimated from cryo-EM (Table 1, lipid composition 17), and this updated simulation displayed analogous asymmetric leaflet thinning and outer leaflet headgroup exclusion displayed by the CG-MD simulations with the initial lipid composition (Fig. S19) (*36*).

**Fig. 5.**
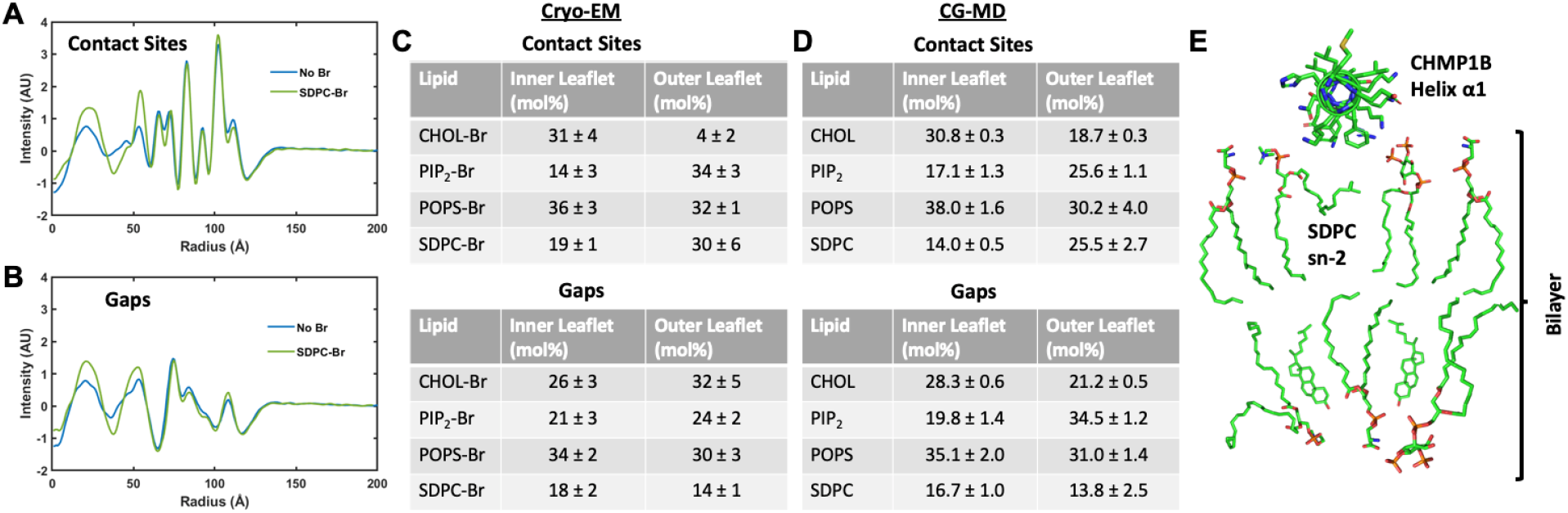
Analysis of lipid bilayer composition at membrane-protein contact sites and between contact sites from simulation and experiment. Radial profile of cryo-EM density A) at CHMP1B contact sites and B) between contact sites for the unbrominated and SDPC-Br samples. See Fig. S27C-D for all profiles. C) The molar composition of each leaflet at membrane-protein contact sites (top) and at the gaps (bottom). Uncertainty was calculated as the standard deviation from independent calculations from each of the two half maps. D) The corresponding compositions for these two sections from CG-MD are shown in for the contact sites (top) and at the gaps (bottom) from simulations using the enriched lipid composition. Uncertainty was calculated as the standard deviation between three replicates of CG-MD trajectories. E) Cartoon model of bilayer structure and composition at CHMP1B contact sites.

Both simulation and experiment indicated that cholesterol was enriched asymmetrically in the inner leaflet versus the outer leaflet at bilayer-protein contact sites. We confirmed the behavior of cholesterol by solving structures with different concentrations of unbrominated cholesterol (Table 1, lipid compositions 4-9). We also took advantage of the fact that cholesterol has a lower Coulombic potential than phospholipids. As the concentration of cholesterol in the lipid mixtures increased, the inner leaflet decreased in intensity relative to the outer leaflet. This observation is also consistent with cholesterol enrichment in the inner leaflet (Fig. S20).

We also estimated that PIP2 was highly enriched not only in the outer leaflet compared with the inner leaflet, but also in the membrane nanotube compared with the initial bulk composition of the liposomes. In light of this, we next investigated the importance of PIP2 for tubule formation by carrying out membrane remodeling assays with vesicles containing no PIP2 (Table 1, lipid composition 2). In this case, CHMP1B is unable to stably bind and remodel vesicles, suggesting that electrostatic interactions are indispensable for its functions (Fig. S13). Similarly, K16E or R20E mutations on CHMP1B helix α1 ablate membrane binding in the presence of PIP2, presumably due to a lack of favorable electrostatic interactions between basic residues on the protein and anion-rich lipid bilayers (Fig. S18).

Finally, due to the pinning of PIP2 by K and R residues and of SDPC by F9 and F13 residues, the diffusion coefficients of lipids in the outer leaflet of the bilayer are dramatically slowed in CG-MD simulations (Fig. S21). This reduction in diffusion coefficients has functional implications for membrane fission as membrane shearing becomes possible as the diffusion rate of lipids decreases (*5*). The necessity of SDPC and PIP2 for membrane binding and constriction highlights the importance of local lipid composition as a regulator of membrane remodeling.

## Discussion

We combined cryo-EM, heavy atom-labeled lipid probes, and CG-MD simulations to investigate membrane structure and mechanics in ESCRT-III constricted tubules. These efforts revealed how phenylalanine residues from CHMP1B α1 displace headgroups to “score” the surface of the bilayer. Flexible, polyunsaturated lipid tails enrich within the furrow created by these Phe residues, where they contort to interact with the hydrophobic side chains. These surface defects coalesce into a helical furrow that may, along with F9 and F13 headgroup displacement, stabilize the bilayer at this high degree of curvature by decreasing lipid density in the convex outer leaflet. We also showed that this high membrane curvature system generates compositionally asymmetric leaflets.

While lipid tail backflipping is possible for any of the lipid tails, polyunsaturated tails are exceptionally flexible and therefore have low energy barriers to “backflipping” (*4, 41–46, 49, 51, 52*). This role for polyunsaturated lipids in stabilizing curvature and filling in packing defects by undergoing curvature-driven shape changes is consistent with previous reports that SDPC facilitates membrane fission and that polyunsaturated tails prefer to reside near the lipid/water interface, especially in curved bilayers (*4, 46, 49*).

Together, our findings on compositional variation within the leaflets of formed tubules, and the membrane remodeling assays with different initial lipid compositions, are consistent with lipid compositions having a prominent effect on the energy required to bend a membrane. Membrane remodeling machines have adapted accordingly to possess organelle-specific properties, and cells likely govern membrane remodeling by regulating local lipid composition through clustering, transport, and lipid metabolism (*53*).

We hope the tools presented here will help fill the persistent gaps in our understanding of biological structures that are not defined by linear sequences. Ongoing developments in structural biology—including AI-inferred protein structures, the prospects for multi-state inference, 3D flexibility analysis, time-resolved studies, and in situ structures derived from cryo-EM and cryo-ET—will enable studies of still higher-order biological assemblies, including membranes, biomolecular condensates, and the interactions between these states of matter (*40, 54–56*). We anticipate that the ability to validate or falsify simulations with heavy-atom cryo-EM will enable new questions to be answered about these critical fluid assemblies, their interactions, and the dynamics governing their functions.

## Acknowledgments

We thank members of the Frost and Grabe labs for helpful discussion, especially P. Thomas for computational assistance, H. Nguyen and H. Aydin for assistance with ESCRT proteins, and D. Argudo and N. Bethel for early discussions concerning membrane tubule mechanics and simulations. We also thank the UCSF Center for Advanced cryoEM, including D. Bulkley, G. Gilbert, M. Harrington, and Z. Yu. Structural biology applications used in this project were compiled and configured by SBGrid. The GPUs used for cryo-EM data processing were donated by the NVIDIA Corporation.

## Author contributions

Conceptualization: AF, MG, FRM, JL Methodology: FM, AM, MT, JL Investigation: FM, AM, JL Visualization: FM, AM, JL

Funding acquisition: AF, MG Project administration: AF, MG Supervision: AF, MG

Writing – original draft: AF, MG, FRM, JL, AM Writing – review & editing: AF, MG, FRM, JL, AM

## Data and materials availability

The cryo-EM maps have been deposited into the Electron Microscopy Data Bank (accession numbers EMD-XXXXX–XXXXX). The cryoEM micrographs have been deposited in the EMPIAR database (https://www.ebi.ac.uk/empiar/) with accession numbers EMPIAR-XXXXX-XXXXX. Any other relevant data and materials are available from the corresponding authors upon reasonable request.

## Supplementary Materials

### Materials and Methods

#### Synthesis and characterization of bromolipids

Bromolipids were synthesized as previously described (*57*). 1-100 mg of each lipid was dissolved in CHCl3 to 1-10 mg/mL. The lipid solutions were stirred on ice, and bromine (stoichiometric with the number of double bonds in the lipid) was added dropwise. The solution was stirred on ice in the dark for 1 hr. Solvent and excess bromine were removed by application of vacuum overnight in the dark. Brominated lipids were then aliquoted and stored at -80 °C until needed. To maintain accurate lipid stock concentrations, no further purification was performed. Consequently, the brominated lipids also contain some unbrominated lipid, whose quantities were determined by mass spectrometry and nuclear magnetic resonance (NMR) spectroscopy.

Proton nuclear magnetic resonance (^1^H NMR) and carbon nuclear magnetic resonance (^13^C NMR) spectra were recorded on a Bruker Avance III HD 400 spectrometer in the UCSF NMR Core Facility. NMR spectra were recorded in CDCl3, except for PIP2-Br, which was recorded in 20:10:1 CDCl_3_:MeOD:D_2_O. Electrospray ionization mass spectra were recorded with a Waters SQD2 mass spectrometer in the Stanford University Mass Spectrometry Facility. Gas chromatography mass spectra were recorded with a HP 7890/5975 GC-MS, equipped with a single quadrupole MS with electron impact ionization source and a 30-meter HP-5MSI (5% phenyl)-methylpolysiloxane column with an inner diameter of 0.25 mm and a film thickness of 25 µm. The carrier gas was helium with a flow rate of 1 mL/min. Chemical structures and NMR spectra of unbrominated and brominated lipids are in Fig. S1-10, and average numbers of bromine atoms per lipid determined by MS and NMR are in Table S1.

We next examined the effects of bromination on the behavior of brominated lipids within a lipid monolayer at the air-water interface. We measured pressure-area isotherms for pure unbrominated and brominated lipids as described in detail previously (*58*). Briefly, we used a KSV NIMA KN 2002 (Biolin Scientific) Langmuir-Blodgett system with a 273 cm^2^ Teflon trough and symmetric Delrin barriers. A piece of Whatman #1 filter paper was used as a Wilhelmy plat to monitor the surface pressure in the trough. We added water to the clean trough and used a vacuum line attached to a Teflon tip to remove any contaminants from the liquid surface. Lipids (1 mM in chloroform) were spread onto the surface of the water with a glass microsyringe and solvent was allowed to evaporate for 10 min. The barriers were then compressed at a rate of 10 mm/min and pressure-area isotherms were collected. All isotherms were recorded at room temperature on a subphase of Milli-Q water and are shown in Fig. S14.

#### Sample preparation

The N-terminal domain of IST1 and full-length CHMP1B were expressed and purified as previously described (*27*). CHMP1B mutants were produced with standard techniques using the QuikChange II kit.

Membrane remodeling assays were carried out as previously described to create CHMP1B-IST1 copolymer-bound membrane nanotubes (*27*). Small unilamellar vesicles (SUVs) were formed by extrusion. To protect the polyunsaturated lipids from oxidation during the formation of vesicles and subsequent membrane remodeling reaction, we add 300 nmol of the desired lipid mixture to a glass scintillation vial under a nitrogen atmosphere (see Table 1 for lipid mixtures). The solvent was evaporated under a stream of nitrogen while rotating the vial. The resulting film was re-dissolved in 100 µL chloroform and evaporated again. The vial with the dried lipid film was placed under house vacuum in the dark for 2 hours. 250 µL of GF buffer (25 mM Tris pH 8.0 and 125 mM NaCl) was added to the scintillation vial and incubated for 10 min in the dark. The lipid film was next resuspended to form multilamellar vesicles by vortexing. SUVs were formed by extrusion 31 times through a polycarbonate membrane with 50 nm pore size using the Avanti Polar Lipids Mini Extruder. SUVs were either used immediately or aliquoted, snap frozen in liquid nitrogen, and stored at -80 °C. To form copolymer-bound nanotubes, SUVs (0.5 mg/mL in GF buffer) were incubated with 5 mg/mL CHMP1B (in GF buffer) for approximately 6 hours at RT in the dark. IST1 (in GF buffer + 5% glycerol) was added to 10 mg/mL, and the solution was incubated overnight at RT in the dark.

Samples for negative stain EM and cryoEM were prepared similarly to previous reports (*27*). Briefly, for negative stain EM, 4 µL of the membrane remodeling mixture were applied to glow-discharged, 200 Cu mesh carbon-coated grids (Electron Microscopy Supplies, USA). Grids were stained with 0.75 % (w/v) uranyl formate (Structure Probe, Inc.). All samples were imaged on an FEI Tecnai T12 120 kV electron microscope equipped with a Gatan UltraScan 895 4k CCD camera. For cryoEM, 4 µL of membrane remodeling mixture were pipetted onto glow-discharged R1.2/1.3 Quantifoil 200 Cu mesh grids (Quantifoil) in a Mark IV Vitrobot (FEI). After a 10 s wait time at 19 °C and 100 % humidity, grids were blotted for 4 s with Whatman #1 filter paper and plunged into liquid ethane. Grids were stored under liquid nitrogen until imaged.

#### Cryo-EM imaging

Cryo-EM data were collected on 300 kV FEI Titan Krios or 200 kV FEI Talos Arctica microscopes. The Krios was equipped with a Gatan BioQuantum energy filter and Gatan K3 direct electron detector. The Krios was operated at a nominal magnification of 105kx with a total dose of 67 e^-^/A^2^. The Arctica microscope was operated at a nominal magnification of 28kx or 36kx with a total dose of 61 e^-^/A^2^. The Arctica was equipped with a Gatan K3 direct electron detector. The cameras on both microscopes were operated in correlated double sampling and super resolution modes. Samples with lipid compositions 2-8 and 18 were imaged with the Arctica, while samples with lipid compositions 1 and 10-17 were imaged with the Krios (all lipid compositions are summarized in Table 1). Images were collected with a 50 µm C2 aperture and 100 µm objective aperture. The defocus was varied between -0.6 and -2 µm. Semi-automated data collection was carried out with SerialEM.

#### Data processing

The software programs used to process and visualize data were compiled and configured by SBGrid (5*9*).

We processed the cryo-EM data in a manner similar to that reported previously, except that filaments were automatically picked from the micrographs with SPHIRE-crYOLO (*27, 60*). The rest of the data processing was performed in Relion 3.08 (61). All of the cryo-EM datasets were processed with the pipeline shown in Fig. S11 unless otherwise noted. Sample motion was corrected with MotionCor2 with dose weighting, and defocus values were estimated with CTFfind4 (62, 63). One round of 2D classification was performed to create a more homogenous data set with similar particle diameters and eliminate particles with poor resolution. A hollow, smooth cylinder was used as an initial model for 3D auto-refinement without a mask, and ‘ignore CTFs until first peak’ option for CTF estimation was used. Previously determined helical parameters were used as the initial values in the refinement. Subsequently, these particles underwent a single round of 3D classification to separate 17 and 18 asymmetrical subunits (asu) per turn structures (*27*). Specifically, the number of asymmetrical units, initial twist, and initial rise parameters were changed to select for either variant of the filament. During 3D classification, we used a mask that included both the protein and lipid bilayer, whereas further auto-refinements used masks that solely included the protein. High resolution 3D classes with the desired symmetry were chosen, and each group of classes with the same symmetry went through another round of 3D auto-refinement. Afterward, the refined particles for both the 17 and 18 asu structures went through two rounds of per-particle CTF refinement, with 3D auto refinement between each round, and the final reconstructions were post processed with automatic B factor estimation. The relevant parameters for each step are listed in Table S2. A summary of the pipeline is shown in Fig. S22. We used the CHMP1B/IST1 atomic model previously reported, PDB 6TZ5, which fit all of the 17 subunit per turn reconstructions reported here. All data collection and processing statistics are summarized in Tables S3-5.

We examined whether data collection and analysis parameters had any effect on the final 3D reconstructions by examining the radial profiles. As shown in Fig. S23 applying low-pass filters, helical symmetry, and normalization to CHMP1B helix α1 had no meaningful effects on the bilayer or protein density. While not used for discussion in the main text, 18 subunit per turn reconstructions were built and refined. Structures are shown in Fig. S24.

We estimated the resolution of the local resolution of the lipid bilayer to be ∼6 Å by gold standard FSC (GS-FSC) analysis. We created a mask that only encompassed the bilayer and performed an auto-refinement with FSC resolution estimate in Relion. Consequently, we used the same mask to apply a custom 6 Å low-pass filter to the bilayer in the reconstructions while maintaining the high-resolution, ∼3 Å, features of the protein coat. All the cryo-EM densities shown in the main text have been processed in this way. RELION auto-refinement divides the cryo-EM particles randomly into two halves and refines each half of the data independently. These independent half maps are used to estimate resolution based on the GS-FSC method. We took advantage of these half maps to estimate uncertainty in our analyses as the standard deviation of the measured quantity between the two half maps, shown in Fig. S25.

#### Effects of varying lipid composition

We systematically varied the lipid composition of the vesicles in order to examine the effects on remodeling by CHMP1B and IST1. Vesicles and filaments formed with bromolipids were indistinguishable from those formed with unbrominated lipids (see Fig. S13 and Fig. S18). However, changing the lipid composition of the vesicles in other ways did produce morphological changes in the membrane-bound CHMP1B/IST1 filaments. Substituting POPC for SDPC (composition 3) caused the resulting filaments to be significantly wider than those formed from composition 1 (see Fig. S13). The mole percent of cholesterol was also varied between 0 and 50, while keeping the ratio of the other three components constant (Table 1). We repeated the previously described membrane remodeling assay with vesicles made from each of these lipid compositions and visualized them with negative stain EM. As shown in Fig. S13A and S20D, CHMP1B and IST1 remodel vesicles from compositions 4 – 8 (0 – 40% cholesterol) into membrane-bound filaments that are indistinguishable from those formed from lipid composition 1 (Table 1, 18% cholesterol). However, at 50% CHOL, CHMP1B and IST1 bound to vesicles but could not remodel them into thin nanotubes. Finally, removing the PIP2 from the lipid mixture (Table 1, lipid composition 2) abolished protein binding to the vesicles (Fig. S13I, M).

SDPC is known to reduce the bending rigidity of lipid bilayers, while cholesterol is known to increase the bending rigidity of fluid phase bilayers. Both replacing SDPC by the less flexible POPC and increasing the cholesterol concentration causes the filaments to become wider. These observations are consistent with a simple mechanical model in which CHMP1B and IST1 apply force to the bilayer, whose final diameter depends on the bending rigidity and spontaneous curvature of the bilayer. As SDPC and cholesterol likely change both the bending rigidity and spontaneous curvature of the bilayer, more work is needed to disentangle these effects. The failure of tubule formation to occur after removing PIP2 from the lipid mixture highlights the importance of electrostatic interactions between CHMP1B and the bilayer. Similarly, mutation of K16 and R20 to Glu abolishes CHMP1B binding to bilayers containing PIP2 (Fig. S18B). Cholesterol is significantly different structurally than the phospholipids in the lipid mixture and has lower electron density. We exploited this difference to confirm that cholesterol is overall enriched in the inner leaflet of the bilayer, consistent with the data from CHOL-Br (Table S6). We tracked into which leaflet cholesterol partitions by titrating from 0 to 40 mol% cholesterol and quantifying how much each leaflet decreased in electron scattering intensity. The inner leaflet decreases in intensity preferentially up to 30 mol% cholesterol, after which it saturates (Fig. S20).

We also conducted the remodeling assay substituting in the 19-iodo derivative of cholesterol (CHOL-I; Table 1, lipid composition 14), which is commercially available from Sigma-Aldrich, as a comparison to the brominated derivative we prepared. Cryo-EM reconstructions and normalized intensity curves, with comparisons to the normal unhalogenated mixture (Table 1, lipid composition 1) and the CHOL-Br mixture (Table 1, lipid composition 11), are shown in Fig. S26.

#### Analysis of leaflet compositions

We analyzed the leaflet compositions by comparing the increase in intensity of each leaflet for brominated samples versus the unbrominated reference according to the procedure shown schematically in Fig. S27. First, we investigated the structure and composition of the membrane-protein contact sites and the region of the bilayer farthest from the protein (gaps between contact sites). To facilitate this comparison, we rotated each Z slice of the cryo-EM reconstructions so that membrane-protein contact sites were aligned axially. We then projected the central 30 % of each reconstruction along the Z axis, improving the signal to noise ratio. We then calculated radial averages and the area of each leaflet curve above 0, which we defined as the solvent background intensity. The areas for each leaflet of the samples without brominated lipids were subtracted from the areas of the respective leaflets of each of the reconstructions with brominated lipids. This yielded the extra scattering in each leaflet due to each brominated lipid. These values were then normalized to the numbers of bromine atoms per molecule (see Table S1). The values for each leaflet were summed, and the fraction of the total for each brominated lipid was calculated, such that the composition of each leaflet adds to 100 mol%. Second, the bulk, radially averaged compositions of each leaflet were estimated by applying the same procedure but omitting the rotation alignment of the membrane-protein contact sites. Both analyses were performed separately on each of the half maps, and the reported uncertainties are the standard deviations between the analyses from the two half maps. The estimated bulk leaflet compositions are shown in Table S6.

#### Sub-stoichiometric SDPC-Br

We probed whether SDPC-Br has the same affinity for the CHMP1B-bilayer contact sites as SDPC. We performed a titration by substituting 25 % and 50 % of the SDPC in lipid composition 1 (Table 1) for SDPC-Br, resulting in lipid compositions 16 and 15 (Table 1), respectively. We then created 3D reconstructions of the resulting filaments as previously described and compared the intensities at the protein-bilayer contact sites. If SDPC-Br behaves exactly the same as SDPC, the intensities at the contact sites will be linear with SDPC-Br mol %.

As shown in Fig. S28, lipid bilayers in which less than 100 % of the SDPC is brominated display intensities at the contact site that are intermediate between samples with 0 % and 100 % SDPC-Br. The change is not completely linear and shows somewhat higher than expected SDPC-Br enrichment when the SDPC-Br mol fraction is between 0 % and 100 %. This result suggests that SDPC-Br has a slightly higher affinity for the CHMP1B-bilayer contact sites than SDPC or that SDPC-Br is more enriched in the lipid bilayer within the protein coat than SDPC. Lipid bilayer vesicles are often seen protruding from the ends of filaments (Fig. S13A, F, J and Fig. S20A). This scenario provides the opportunity for some lipids to be enriched in the highly curved lipid bilayer nanotube inside the protein coat, while others are enriched in the less curved vesicle. We note that the differences between SDPC and SDPC-Br are likely small as the contact sites do not saturate at lower SDPC-Br mole fractions.

#### Summary of all simulations

Five sets of simulations were conducted. The first set of membrane tubule simulations produced most of the data discussed in the main text (all main text simulation data except Fig. 5). This set consisted of 10 replicate simulations using a CG Martini 2.2 model of two turns of the IST1-CHMP1B protein coat (PDB code: 6TZ5) using a lipid molar composition corresponding to starting lipid composition 1 (Table 1). Full results are presented in Fig. S11. The second set was identical to the first except in silico mutagenesis was used to make the F9A + F13A double mutant of CHMP1B and only 3 replicates were run (full results in Fig. S16). The third set were run using the experimentally derived membrane tubule composition, again with WT CHMP1B (Table 1, composition 17). This more highly anionic lipid mixture was substantially less stable than composition 1, and hence the equilibration procedures and run parameters were adjusted as described in the following sections (full results in Fig. S19). The fourth simulation was run to compare features of the tubules formed in the presence of protein, from simulation set 1, with a tubule of equal curvature but formed without protein. A tubule was formed from spontaneous assembly using a lipid mixture corresponding to composition 1 and run through equilibration and production in the absence of protein. Full results are in Fig. S17. The fifth set consisted of one replicate of a simple flat-bilayer simulation corresponding to lipid composition 1.

#### Simulation sets 1 & 2 summary: CHMP1B/IST1 WT (1) and F9A + F13A CHMP1B mutant (2) – lipid composition #1

The structure of two turns of the WT CHMP1B-IST1 protein coat (34 subunits of each protein, PDB code: 6TZ5) was centered in a 30 x 30 x 18 nm simulation box and converted to CG Martini 2.2 representation. For the F9A + F13A simulations, the protein structure was first edited to contain the mutations and then coarse-grained. The protein was fully position restrained throughout simulation to maintain the cryo-EM resolved protein structure in relation to the membrane. Martini model lipids were chosen to correspond to those used in cryo-EM (Table, 1, lipid composition 1): SDPC (Martini name PUPC), POPS (Martini name POPS) and CHOL (Martini name CHOL) were directly available from the Martini website. A di-oleic PIP2 model was constructed from the Martini model 1-palmitic-2-oleic PIP2 (Martini name POP2) by changing parameters for the second bead of the CG palmitic tail from the saturated C type to the unsaturated D type. The total number of lipids was tuned using a set of initial tubule self-assembly simulations as described in SI Section “<u>Simulation sets 1&2 tubule self-assembly”</u> with results in Fig. S29, arriving at a total of 1300 with 754 SDPC, 234 POPS, 234 CHOL, and 78 PIP2. Details on lipid placement are described in the following. The simulation box was then solvated using standard Martini water and ions to neutralize overall system charge and reach 150 mM NaCl. Simulations were run using Gromacs 2018.8, using Martini-recommended run parameters. Reaction-field electrostatics were used with a 1.1 nm Coulomb cutoff. The velocity rescale thermostat was used to maintain a temperature of 320 K, with separate temperature coupling groups for protein, lipids, and solvent. The timestep and pressure control method differed between equilibration and production phases and are described in detail in the following. One round of minimization was performed prior to dynamics, which proceeded in three general steps—spontaneous tubule assembly, leaflet equilibration, and production—described in the following. The results described in the main text are from analysis on the last 2.4 μs of the production phase of each of the ten replicates for the WT simulations. Three replicates were run for each mutant simulation, with analysis also on the last 2.4 μs of the production phase of each replicate.

#### Simulation sets 1&2 tubule self-assembly

The initialization protocol was developed to promote rapid tubule assembly by initially placing randomly oriented lipids within the entire cylindrical space encompassed by the protein coat. An even grid of positions spaced 4 Å in each direction was generated, containing points within a 5.75 nm radius cylinder centered in the box along the z-axis. This radius was chosen to avoid placing lipids so near the protein that they would immediately insert into the protein coat rather than joining the tubule. A shell script was then used to iterate through the list of grid positions, using the Gromacs insert-molecules function to add lipids one at a time at each grid position until the desired composition and packing were reached. The order of the list of grid positions was randomized for every new replicate, such that individual replicates started with different initial placements for the lipids. This was done to improve sampling of lipid localization and overcome the limits in lipid diffusion in the protein-associated outer leaflet. After lipids were placed, the systems would be solvated and minimized as described in the Methods. Standard MD was then run using a 0.03 ps timestep and semi-isotropic Berendsen barostat. Tubulation was readily achieved within the low hundreds of ns. Each tubulation phase was run for 1.5 μs before moving to leaflet equilibration, described in the next section.

The initial placement of lipids introduces a strong biasing effect toward rapid formation of a tubule, which is desired for efficiency but also brings up concerns about proper equilibration of lipids between leaflets, amount of water within the tubule lumen, and total amount of lipid within the box. The first two concerns are handled with leaflet equilibration. To identify the proper number of lipids, we ran initial tubulation simulations using a range of total lipid counts, shown in Fig. S29.

#### Simulation sets 1&2 leaflet equilibration

The dense initial packing of lipids means that, during tubulation, there is insufficient opportunity for phospholipid flip-flop to ensure properly equilibrated lipid densities in the two leaflets. We therefore followed the procedure introduced by Risselada and Marrink, which serves to equilibrate leaflet lipid densities—and, for mixed membranes, leaflet compositions—in simulated vesicles (64). A repulsive flat-bottomed position restraint potential that acts on phospholipid tails and cholesterol was applied, oriented perpendicular to the tubule axis (along the y axis of the simulation box) at the top of the simulation box. The force constant was 1 kJ/mol*nm^2^. This introduces two solvated pores at opposite sides of the tubule, at which phospholipids can exchange between the inner and outer leaflets over time until the preferred equilibrium lipid densities and compositions are reached, after which the pore can be closed for production. These pores also allow water and ions to move between the now-connected compartments of the tubule lumen and the rest of the bulk solvent in the simulation box, allowing the system to reach preferred densities of each component in these spaces.

To smoothly open the pore and avoid possible bubbling from applying too strong a force from the repulsive potential on a lipid, pores were slowly introduced with 3 short, 25 ns phases using a reduced timestep of 0.02 ps and increasing the radius of the repulsive potential in each phase: 0.5, 1.0, then 1.5 nm. Finally, 9 μs of leaflet equilibration with a 2.0 nm pore radius and a 0.03 ps timestep was run. This total simulation length was chosen based on monitoring of leaflet compositions over time during equilibration. Pressure coupling was changed to the Parrinello-Rahman semi-isotropic barostat for all phases of leaflet equilibration, and the same barostat was continued for production. Following the 9 μs of 2.0 nm pore equilibration, the 3 short 25 ns phases with reduced 0.02 ps timestep were repeated to close the pore, moving between radii of 1.5, 1.0, and 0.5 nm. An example of leaflet compositions over time during the 9 μs pore equilibration, taken from one replicate of Simulation set 1 (WT CHMP1B-IST1 with lipid composition 1), is shown in Fig. S30. Overall leaflet compositions for each replicate, calculated across the production stages, are shown in Tables S7 (Simulation set 1) and S8 (Simulation set 2).

#### Simulation sets 1&2 production

Following leaflet equilibration, production was run for 3 μs using the same MD parameters. The last 2.4 μs of each production simulation was used for analysis, with a 12 ns frame rate.

#### Simulation set 3 (WT CHMP1B/IST1 composition 17) summary

Simulation of WT CHMP1B/IST1 with lipid composition 17 required modifications to the protocol due to instabilities in the membrane that resulted in persistent bubbling across a range of run parameters. These simulations were ultimately run entirely in NVT, as only turning off pressure control reliably avoided bubbling. Full details on the run parameters, other tested combinations, and anticipated effects on the results are in the following section. The local composition results described in main text Fig. 5 are from analysis on the last 2.4 μs of the production phase of each of the three replicates.

#### Simulation set 3 assembly, equilibration, and production

Differential scattering analysis of tubules formed with halogen-labeled lipids revealed a difference in protein-formed tubule composition from the bulk lipid composition, motivating a set of simulations that match the experimentally-derived composition of 26:32:22:20 SDPC:POPS:CHOL:PIP2 (Table 1, lipid composition 17). We initially attempted to carry out simulations with the same procedures and parameters used for Simulations sets 1 & 2 but found the new membranes to be substantially less stable, producing persistent vacuum bubbles in the simulation box. We believe this to be due to the much higher negative charge density in the membrane from the enrichments in POPS and PIP2.

In an attempt to produce stable tubule systems, many modifications, alone and in combination, to the parameters were tested. These included increasing the applied pressure by the barostat, changing the barostat algorithm, reducing the timestep to 15 fs (timesteps of 20-40 fs are generally recommended for Martini 2.2), and waiting to turn on pressure coupling only once the production phase was started. All of these modifications failed to avoid bubble formation. It was ultimately necessary to maintain an NVT system throughout equilibration and production in order to complete these simulations without bubbling. We therefore proceeded with the modified procedure described below. Constant use of the NVT ensemble introduces a possible mismatch between the density based on the initial fixed volume and the true preferred density of the system at the run conditions. In the ten replicates of the Simulation set 1 system, the average box dimensions at the end of production were 29.81 x 29.81 x 17.75 nm, a 0.6 % change in the x and y dimensions and 1.4 % change in the z dimension from the initial 30 x 30 x 18 nm. So, while the required use of NVT for these simulations is not desirable, and we will seek to develop alternative approaches to stably run these and other compositions with pressure coupling on in future work, the error here is tolerable for this set.

In summary, four main changes were made to the simulation procedure, the first and most important of which is the use of NVT throughout. Second, the initial placement of lipids during setup was changed such that none were placed within the cylindrical region of the repulsive pore-forming potential used for leaflet equilibration, and the simulation could therefore begin directly with leaflet equilibration and the repulsive potential on. Third, the applied radii and force constant of the repulsive potential were changed during the three opening and closing steps of the pore, as were the lengths of each—though the main organization of these phases remained the same. Finally, the length of the leaflet equilibration stage was reduced as was the timestep, and this shorter timestep was also used during production.

After editing the set of grid points where lipids are inserted to remove points within the cylindrical space where the repulsive pore-forming potential is applied for leaflet equilibration, the simulations were initialized with the repulsive potential turned on. This was done to reduce overall simulation time, by removing the tubulation step (as was done for Simulation sets 1 & 2), and to reduce the chance that a lipid might experience a strong repulsive force from the repulsive restraint potential during pore formation.

The simulations were therefore started with the three short equilibration steps to expand the pore: 100 ns at radius 1.5 nm, 50 ns at radius 1.75 nm, and 50 ns at radius 2.0 nm, all using a 0.02 ps timestep and a force constant of 0.5 kJ/mol*nm^2^ for the repulsive potential. Simulations then continued to 6 μs of leaflet equilibration using a 0.025 ps timestep and the same conditions for the repulsive potential. This simulation time was found to be sufficient for the three replicates to converge to similar leaflet compositions (Table S9). The pores were then closed with three additional short equilibration phases: 50 ns at radius 1.5 nm with a 0.02 ps timestep, 25 ns at radius 1.0 nm with a 0.02 ps timestep, and 7.5 ns at radius 0.5 nm with a 0.015 ps timestep.

For production, the timestep was increased back up to 0.025 ps and the simulation was run for 3 μs, with results from the last 2.4 μs of production used for analysis and comparison with the original set of simulations.

While the need to maintain NVT throughout for these simulations is a concern, the lipid tubules for this set were—compositional differences aside—structurally and behaviorally very similar to those of the initial set run with proper pressure coupling, suggesting that the slight mismatch in likely preferred density was tolerable for the system. Further, the 6 μs leaflet equilibration phase allowed sufficient time to properly distribute solvent between the two eventually separated compartments of the tubule inner lumen and the bulk box, relieving one possible source of stresses for the inside vs. outside of the tubule.

##### Simulation set 4 (Protein-free tubule simulation) summary

One simulation of a lipid composition 1 tubule of the same curvature without the protein was run. All parameters used and overall setup were identical to simulation Set 1, with the exception that isotropic pressure coupling was employed, and the leaflet equilibration phase was extended to 30 μs. Production lasted 3 μs, with the last 2.4 μs used for analysis. With semi-isotropic pressure coupling on, and without the protein coat maintaining the tubule shape, the system quickly relaxed to an expanded tubule radius to minimize the curvature energy of the membrane. Therefore, the simulation was run using isotropic pressure coupling, which allowed for fluctuations in the tubule length while maintaining the desired curvature.

##### Set 5 (Flat bilayer) summary

We prepared a flat bilayer with 2023 lipids in the molar ratios of composition 1 in each leaflet in a 36 nm x 36 nm x 14 nm box. The simulation was run using a 0.04 ps timestep for 1 μs.

#### Simulation analysis

Leaflet assignment was performed with single linkage agglomerative clustering on the Cartesian coordinates of the phospholipid phosphate beads at every frame, unambiguously producing two clusters corresponding to the inner and outer leaflets (Fig. S31). For each frame, cholesterols were assigned to a leaflet by determining the nearest phospholipid phosphate to the cholesterol hydroxyl group and assigning the sterol to the same leaflet.

Cylindrical coordinates (*r, θ*) were determined relative to the long axis of the membrane tubule, which was aligned with the z-axis of the box. For each frame we determined the optimal position of the tubule center (*x*_0_, *y*_0_) in the xy-plane that optimally placed all CHMP1B F9 + F13 (or F9A + F13A) residues along the protein filament equidistant from the cylindrical center line using a least-squares fitting procedure. All atomic coordinates for that snapshot were then calculated in that reference frame. For the protein-free tubule, the mid-line of the cylinder was taken as the z-axis of the box. Example resulting cylindrical coordinates for membrane-facing residues of WT CHMP1B are shown in Fig. S32.

To quantify variation in lipid composition and structural properties depending on the position relative to certain features of the protein coat, primarily the helical stripe formed by CHMP1B F9 and F13, we defined zones in (*θ*, *z*) space based on the protein coordinates. Within these zones we calculated distributions and averages of lipid and bilayer properties, for comparison to other zones (see Fig. S12). The primary zone consisted of a 30 ° wedge centered around the helix formed by the positions of CHMP1B F9 and F13. For each frame, the cylindrical coordinates of these residues were gathered, and we used the midpoint of each pair from the 34 subunits to fit a line in (*θ*, *z*) space. The “phenylalanine contact site” zone was defined to fit the 30 ° wedge centered around this line. The zone was extended at the ends an extra 1.2 nm (1/2 the length between pairs of F9 and F13 on adjacent CHMP1B subunits), due to the observed continuity between subunits of the membrane surface defect. The zone was then capped with lines perpendicular to the central phenylalanine-fit line in (*θ*, *z*) coordinate space at each end. All additional zones were defined as successive 30 ° helical slices following the same slope to cover the entire protein-contacting surface bound in between the helix formed by F9 and F13 (compare Fig. S12 with Fig. S32). The zone 180 ° opposite the F9 and F13 stripe is referred to as the “gap between contact site” zone. Properties such as local composition and lipid shapes were then calculated for the various zones over time and compared in order to identify local perturbations that correspond with specific residues and features along the CHMP1B membrane-facing surface.

Headgroup density calculations were performed using the NC3 bead of SDPC, CNO bead of POPS, and the C1 bead of PIP2. A 2D gaussian kernel density estimate on the set of (*θ*, *z*) coordinates from each trajectory was then calculated to compute the heatmaps. The grid spacing for the kernel density estimate was set of 0.835 Å in *z* and 1 ° in *θ*, to correspond with the EM density analysis of the tubule surface. To avoid edge effects, positions within the upper and lower 15 % of the borders in *z* and *θ* were mirrored to effectively expand the box, and the kernel density for that expanded box was calculated and then trimmed back down to the unmirrored box dimensions for plotting. Densities for individual trajectories were subsequently averaged and normalized to span the range from 0 to 1.

Radial density profiles for each lipid bead type were calculated for each leaflet within different zones of the membrane surface relative to the protein coat (zones are shown in Fig. S12) using a gaussian kernel density estimate on the *r* coordinates from each trajectory. The bandwidth was set to 0.5 Å, and densities were calculated from 0 to 80 Å with a 0.2 Å grid spacing. Densities were then normalized by 2*πrΔr* to account for the increasing effective volume at greater *r*, and then multiplied by the number of beads of the specified type within that zone. Thus, densities reflect enrichment/depletion of different lipid species and components. Replicate profiles were averaged to arrive at the final density profile.

Lipid tilt vectors for each tail were calculated by first computing lipid orientation vectors for each tail defined as the vector from the lipid phosphate bead to the last bead of each tail. The bilayer normal was assumed to be the radial component of the cylindrical coordinate system, employing +*r*^ for the inner leaflet and -*r*^ for the outer leaflet. Hence, tilt values of 0 ° correspond to lipids with configurations normal to the local membrane plane regardless of leaflet assignment. Lipids were assigned to zones based on the location of the phosphate bead. Per zone angle distributions for each lipid type and tail were calculated across each replicate using a Gaussian kernel density estimate spanning 0 to 180 ° with a bandwidth of 0.05 ° and a grid spacing of 0.9 °. Replicate profiles were averaged to arrive at the final density profile.

Local compositions per zone were calculated by gathering all non-headgroup beads within each zone and counting the number of beads present from each lipid species. The total number of beads per species was then normalized by the number of beads that make up a whole lipid of each individual lipid type in the model, excluding the headgroups, to get a fractional molar number of each type of lipid present (i.e., 12 beads for SDPC, 11 for POPS, 8 for CHOL, and 11 for PIP2). Headgroups were excluded since they have an outsized impact for PIP2 lipids (5 beads per headgroup versus 1 for SDPC and POPS). The normalized molar values for each lipid type were then summed and used to calculate local molar compositions per zone and leaflet. Compositions from independent trajectories were averaged to arrive at final reported values.

On a per zone basis, we calculated the locations of the bilayer midplane and planes for the outer and inner leaflet glycerols as follows (Fig. S11C). For the midplane calculations the normalized radial densities of all hydrophobic bead types of the inner and outer leaflets (SI Section 16.1) were gathered. Next, the point of intersection along the radius r where the inner leaflet hydrophobic density matched that of the outer leaflet was used to define the midplane. The mean radial coordinate of the glycerol beads in the inner and outer leaflets across the production run within the zone were calculated and values used as the inner and outer leaflet glycerol planes per zone. The per zone leaflet thicknesses were calculated as the differences between the glycerol plane locations and the midplane and were then averaged across the twelve zones and across all simulations to obtain a mean thickness for the entire membrane surface for comparison to cryo-EM. We chose to use the glycerols to calculate leaflet thicknesses, as opposed to other lipid beads, because thicknesses based on the glycerols have been found to better correspond with experimentally derived leaflet thicknesses in with other work (*21*). Leaflet and bilayer thicknesses for all simulation sets are summarized in Table S10.

We calculated lipid diffusion coefficients to determine the impact of the tubule curvature and the protein coat on lipid dynamics. For each individual lipid, the mean squared displacement (MSD) was calculated using unwrapped cylindrical coordinates. On a cylindrical surface, the minimum distance between two points is a helix with arc length 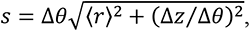 where *Δθ* is the unwrapped change in angular coordinate, Δ*z* is the change in z coordinate, and *r* is the cylindrical radius. The MSD as a function of time was calculated using increments of 12 ns, on the range spanning from 12 ns to 1.2 μs. Windows were sample across the full 2.4 μs of production data. Since the radial position, *r*, of each lipid fluctuates, we calculated *s* using the time-series average value 〈*r*〉 computed over the appropriate time window. Previous work has shown that apparent MSDs computed on curved geometries depends on which lipid bead is used for the calculation (64). We therefore calculated MSDs using the second glycerol beads, as a central element of each phospholipid. Individual MSD time series were averaged over all lipids of a given type for a given leaflet, then a least-squares fit over the sampling timespan from 600 to 1200 ns was used to calculate the diffusion coefficient D (Fig. S21).

#### Backmapping of CG structures to all-atom for figure generation

Standard Martini backmap protocols were followed to generate all-atom configurations from CG simulation frames. All-atom CHARMM36 parameters were used for lipids and protein. A CHARMM36 topology and CG mapping file were constructed for di-oleic PIP2, using pre-existing files for 1-palmitic-2-oleic PIP2. Images presented in the text (Fig. 1C-D, Fig. 2A-D) are taken after the two rounds of minimization during backmapping: without and then with non-bonded forces turned on, to get all-atom representations that most directly correspond to CG configurations sampled during production. A shell script was used in between the first and second rounds of minimization to manually move apart pairs of atoms within 0.5 Å of each other to avoid clashes that would cause the second round of minimization to crash.

---

**Fig. S1.**
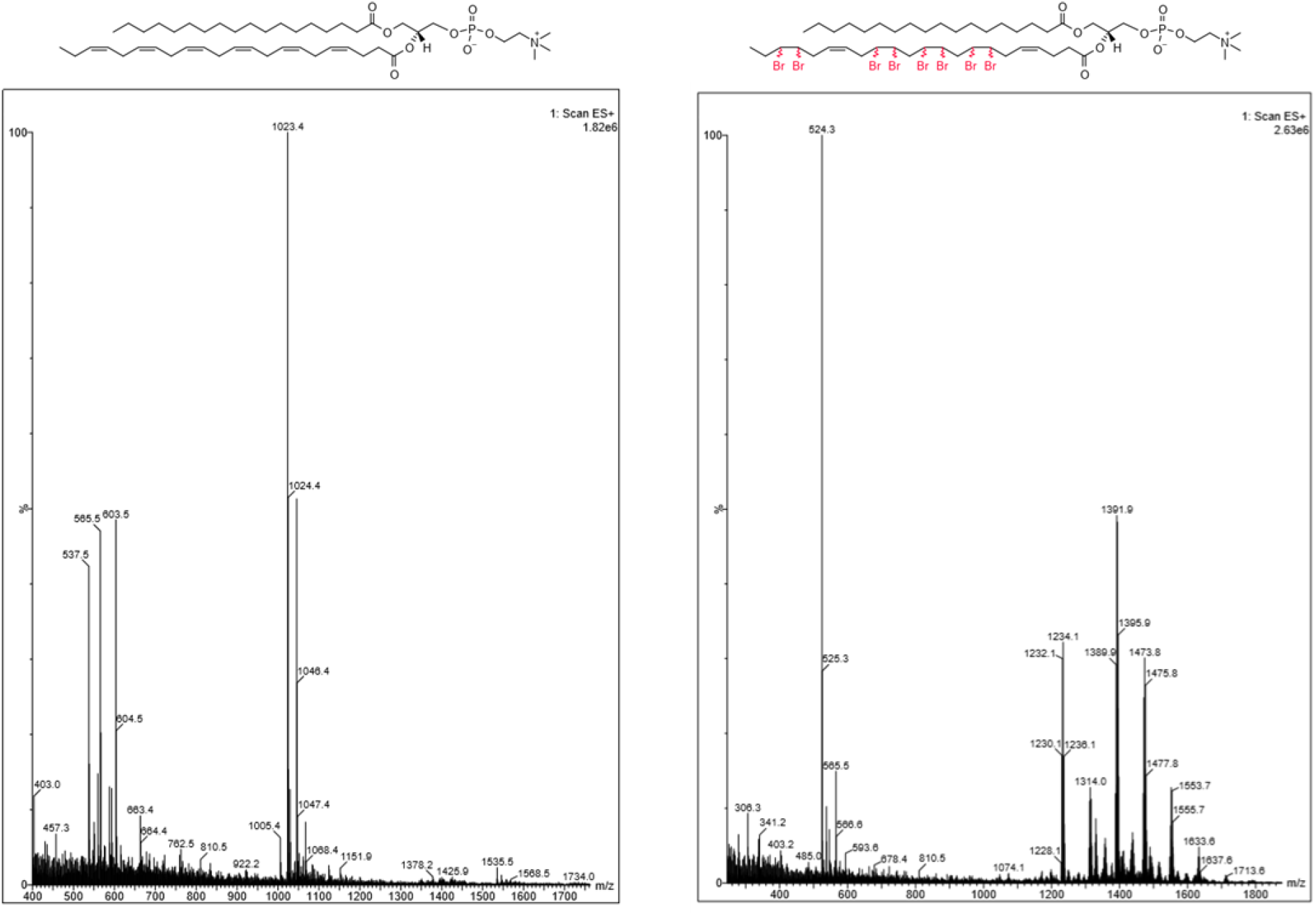
Electrospray ionization mass spectra (ESI-MS) of SDPC (left) and SDPC-Br (right). Partially brominated chemical structure is shown to depict the average level of bromination from the mixture of products.

**Fig. S2.**
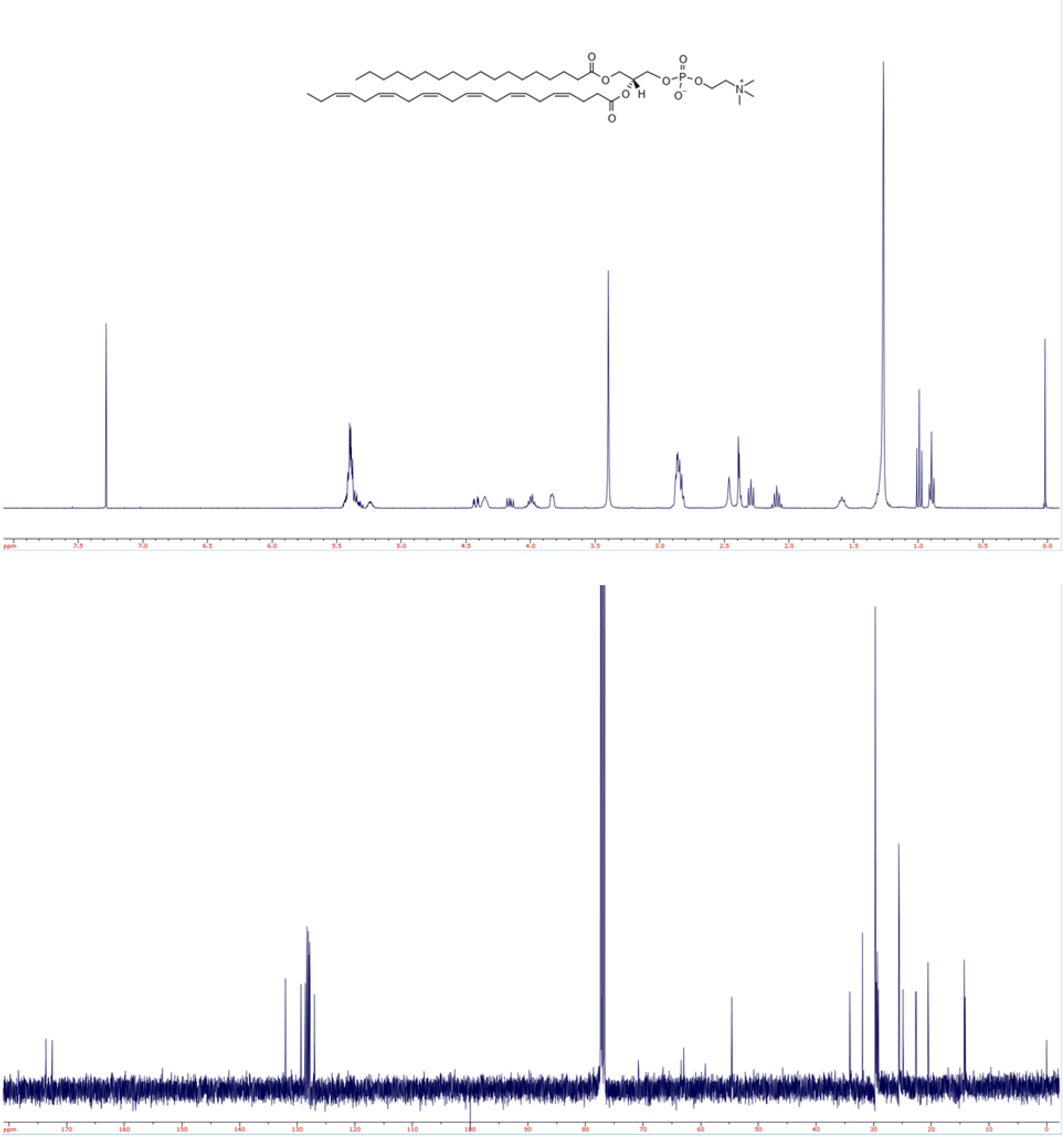
1H (top) and 13C (bottom) NMR spectra of SDPC.

**Fig. S3.**
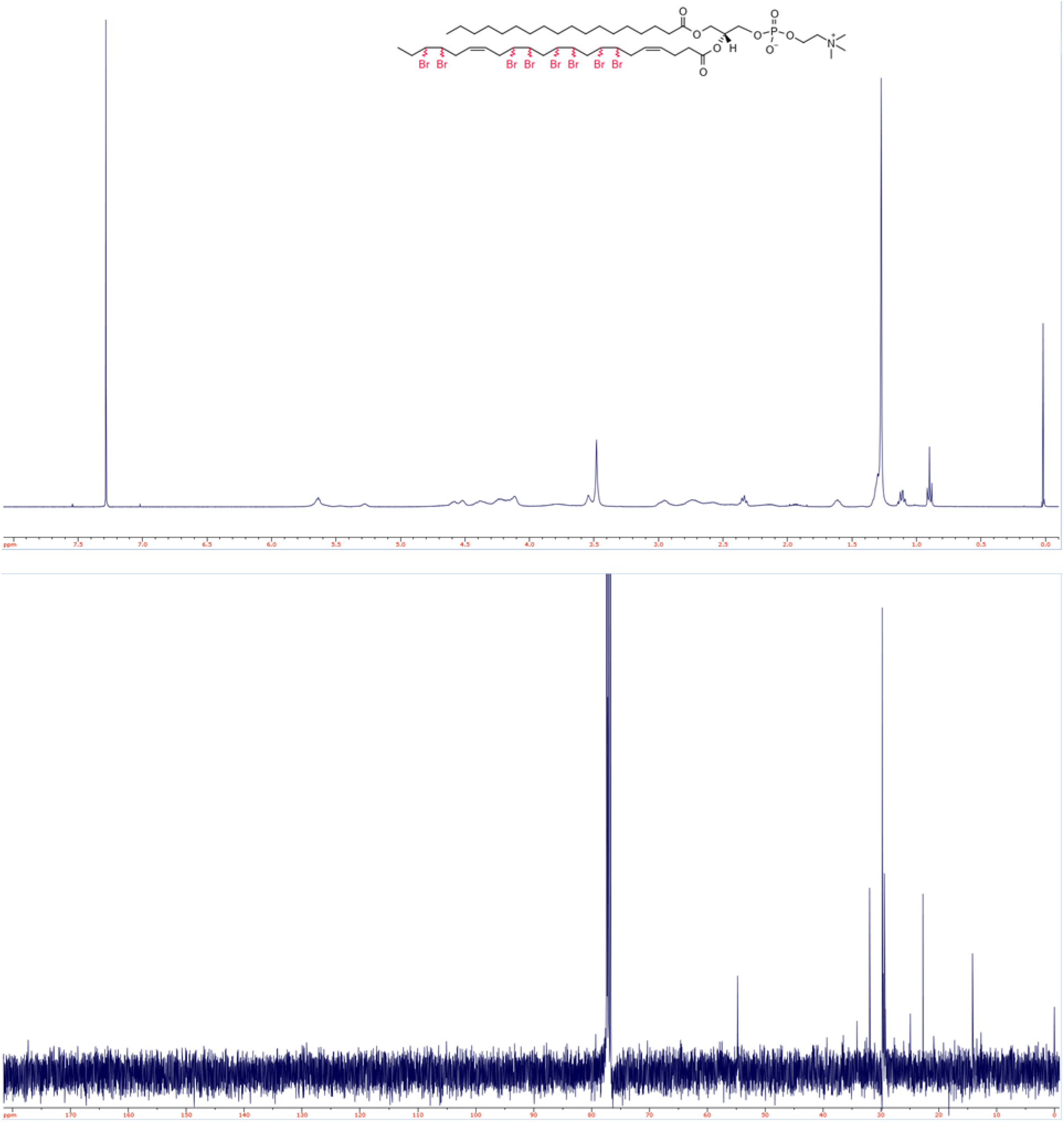
1H (top) and 13C (bottom) NMR spectra of SDPC-Br. Partially brominated chemical structure is shown to depict the average level of bromination from the mixture of products.

**Fig. S4.**
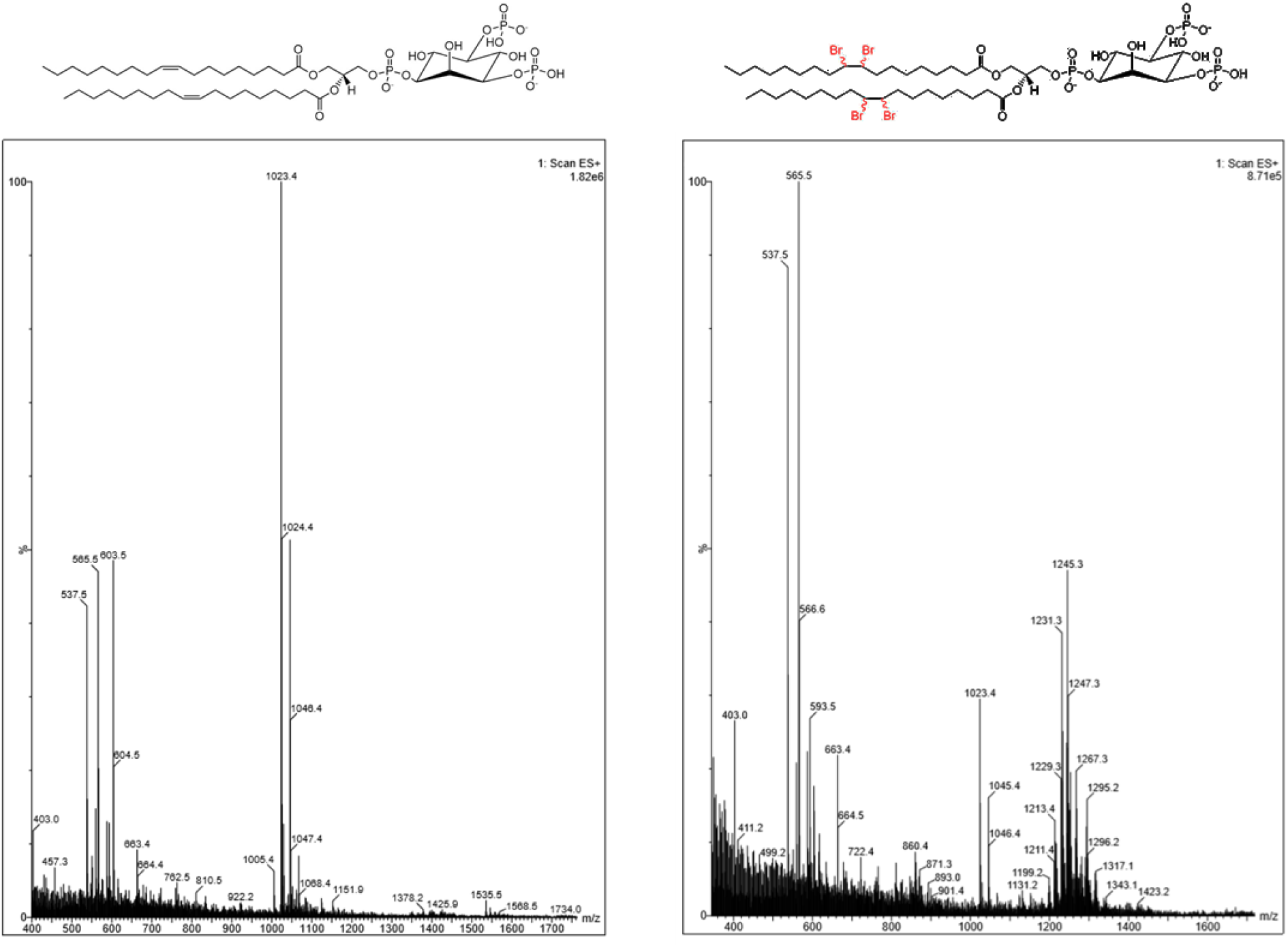
Electrospray ionization mass spectra (ESI-MS) of PIP2 (left) and PIP2-Br (right).

**Fig. S5.**
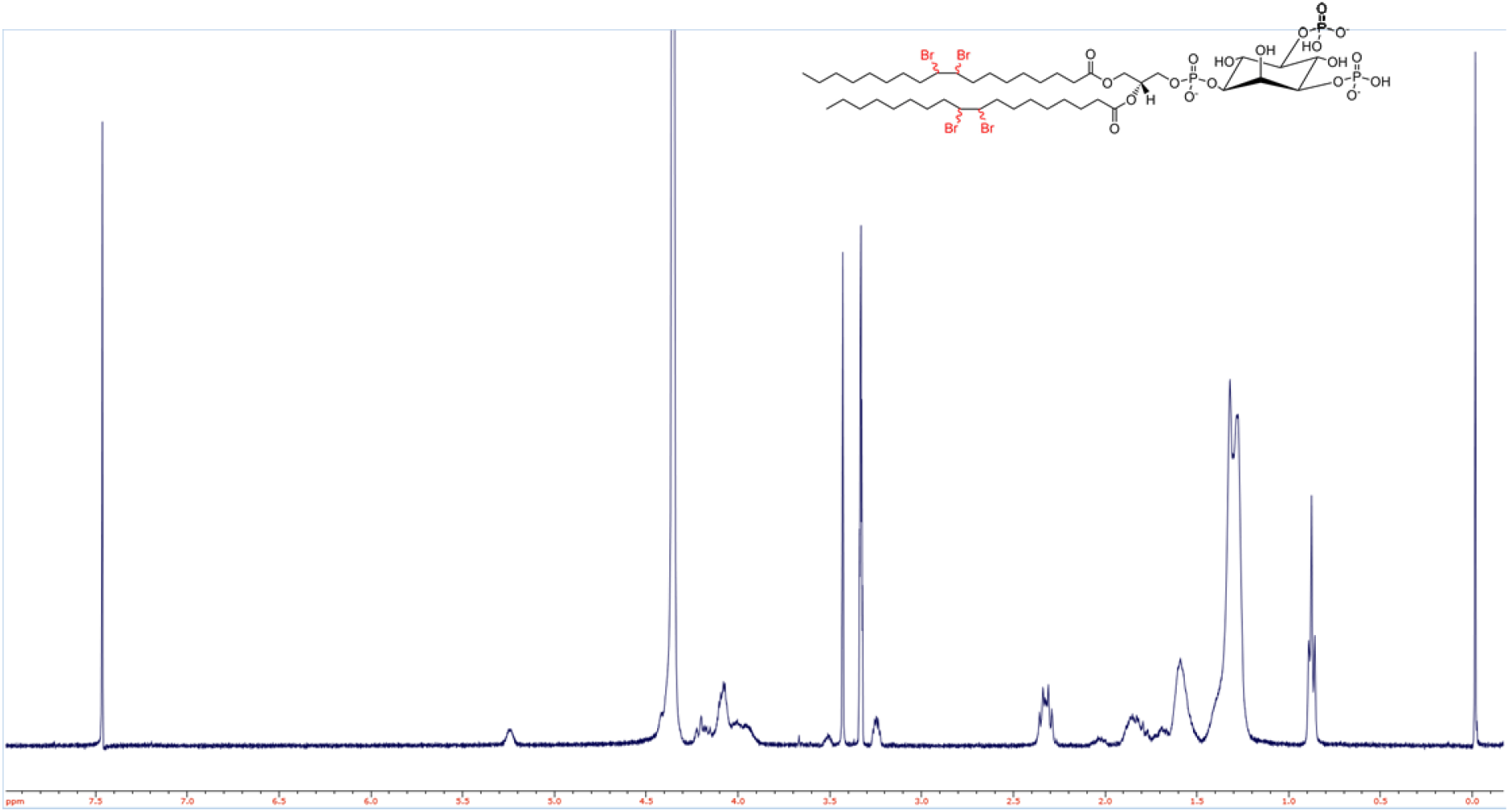
1H NMR spectrum of PIP2-Br.

**Fig. S6.**
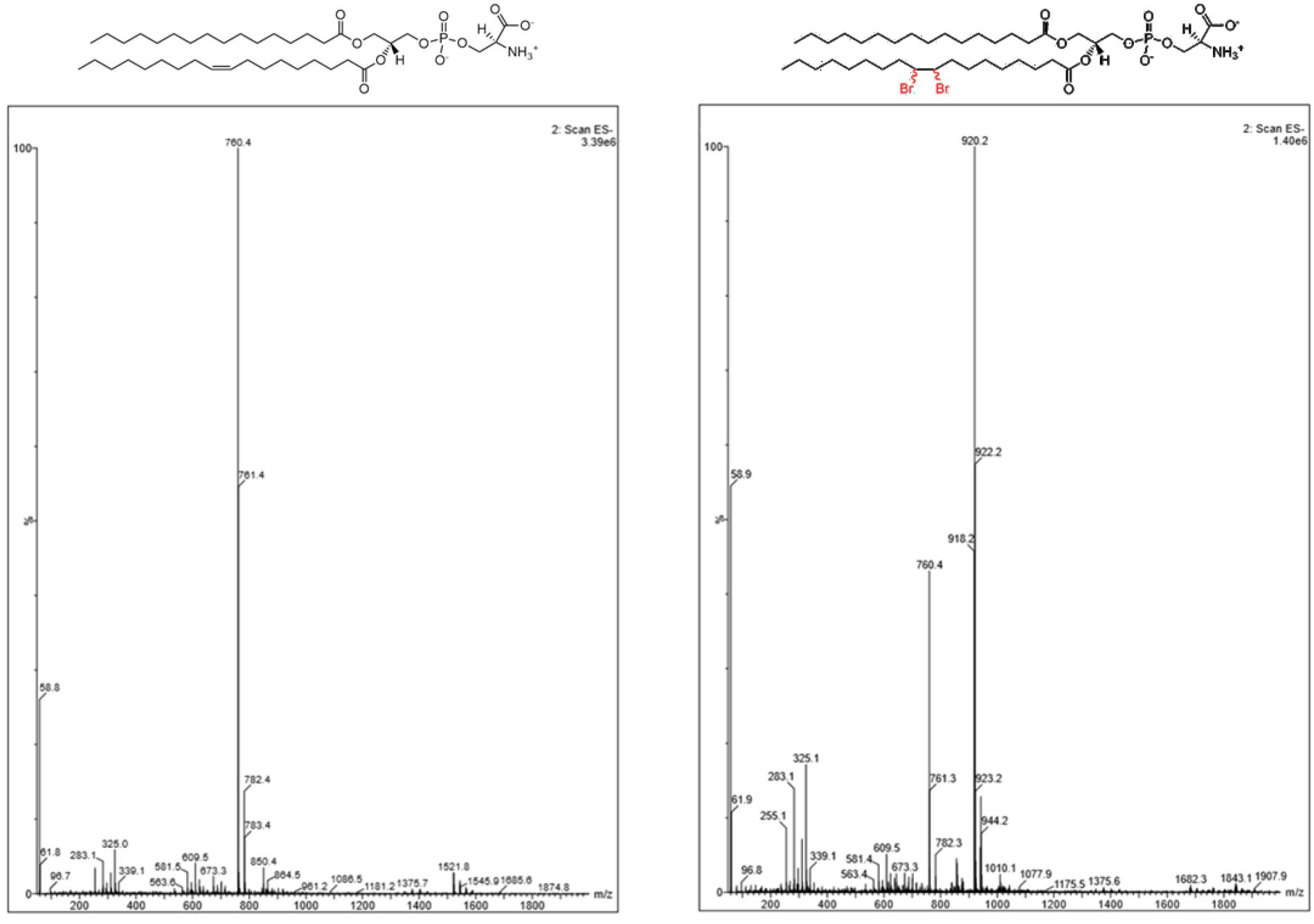
Electrospray ionization mass spectra (ESI-MS) of POPS (left) and POPS-Br (right).

**Fig. S7.**
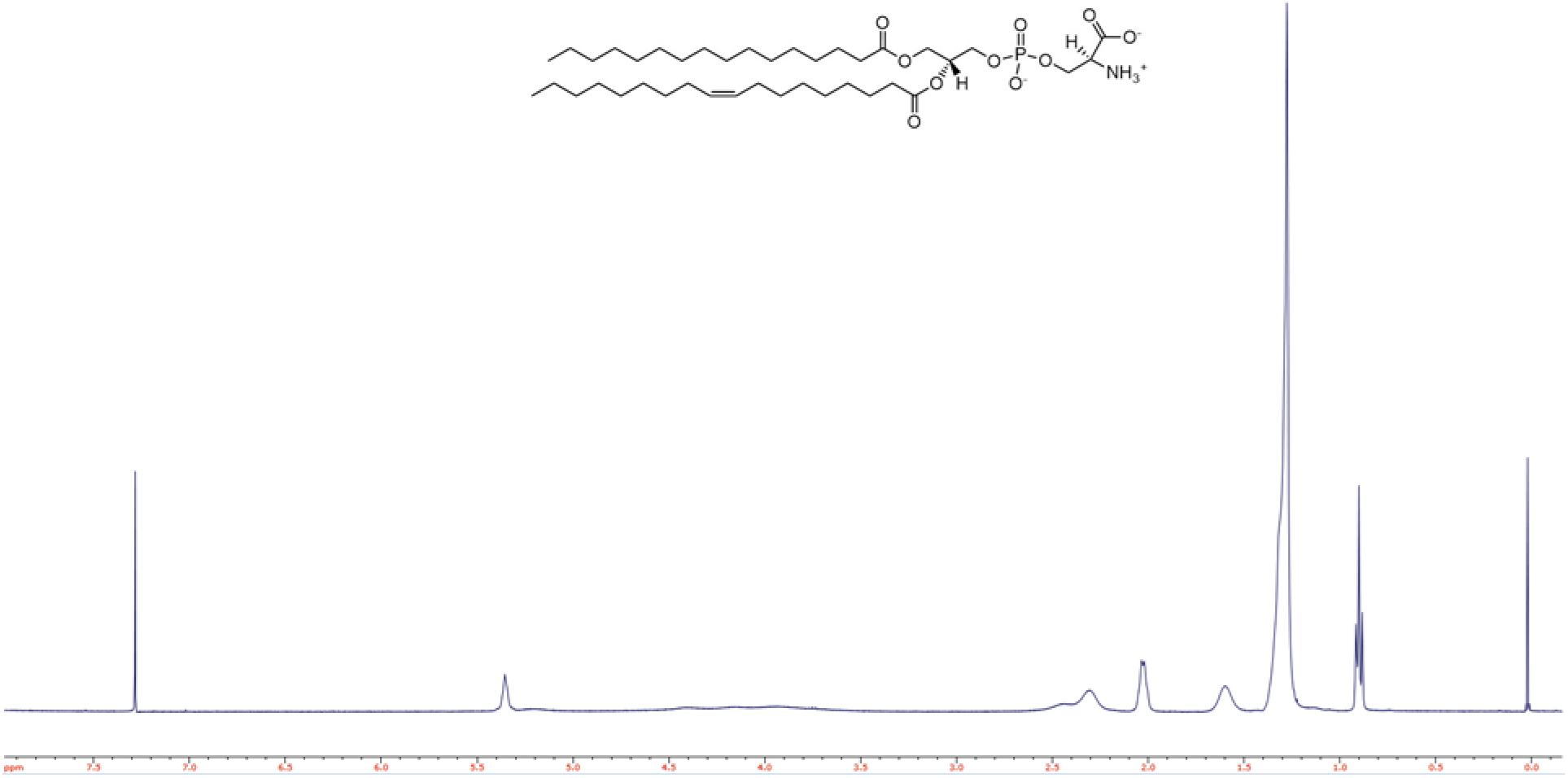
1H NMR spectrum of POPS.

**Fig. S8.**
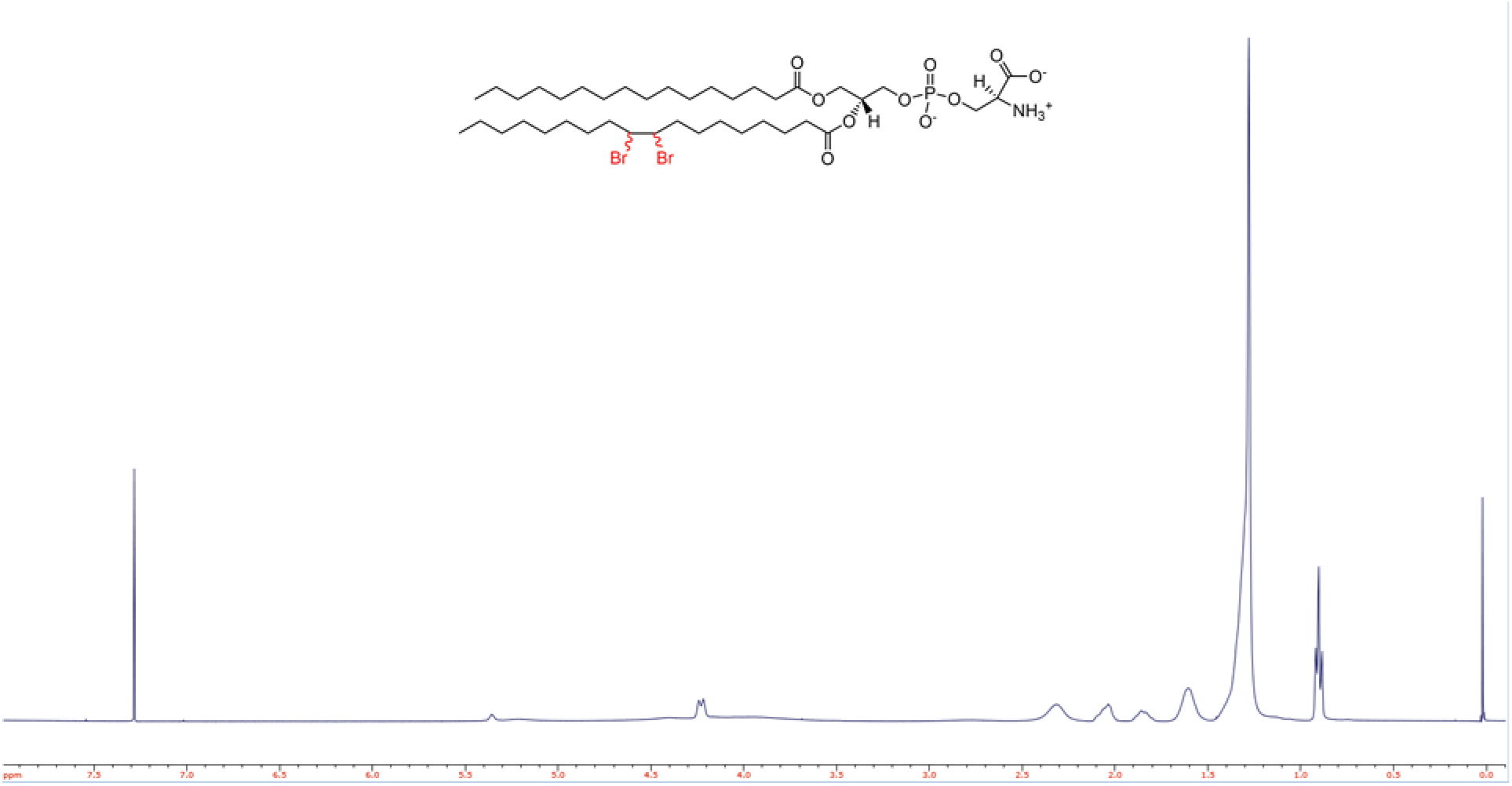
1H NMR spectrum of POPS-Br.

**Fig. S9.**
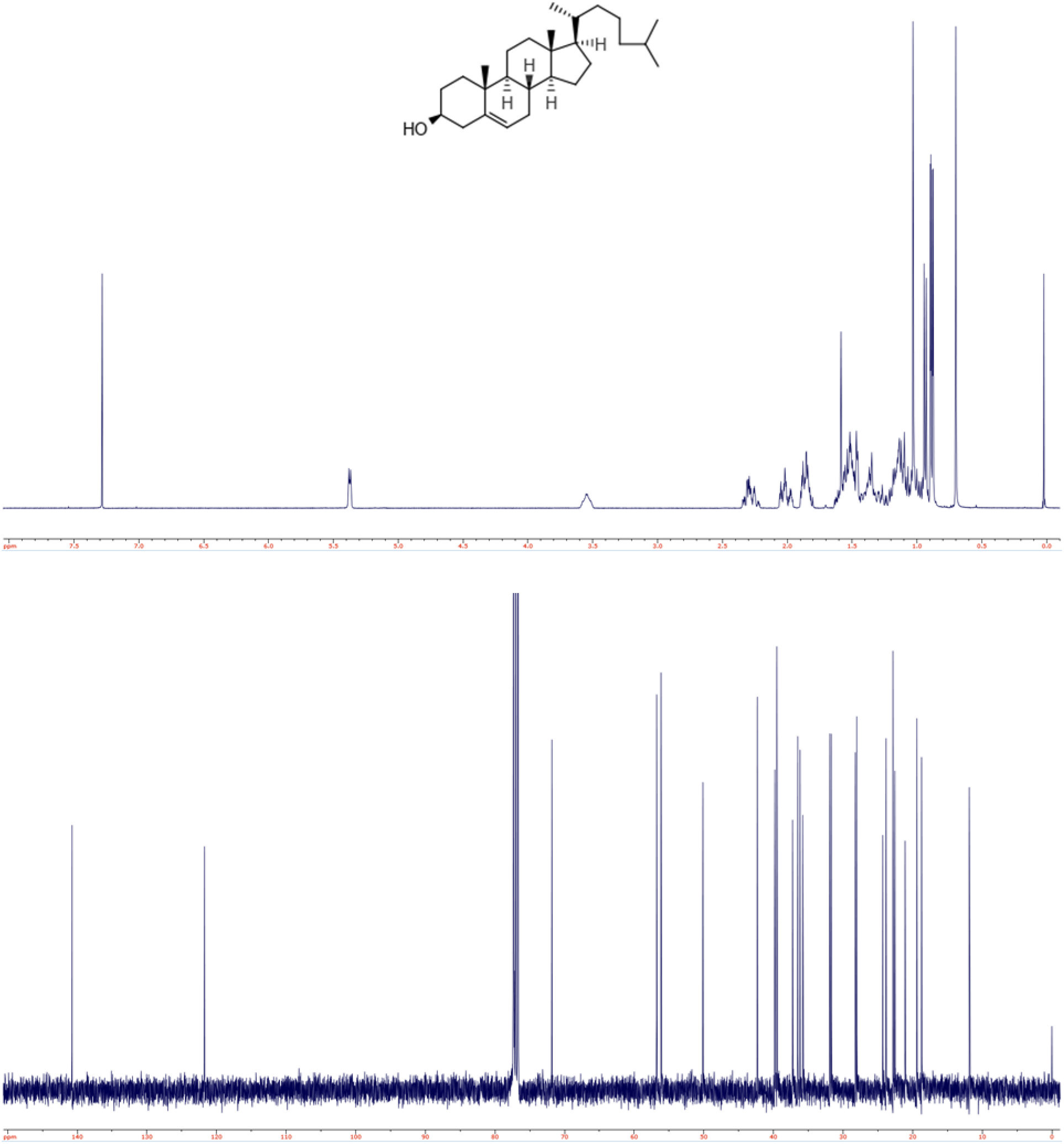
1H (top) and 13C (bottom) NMR spectra of CHOL.

**Fig. S10.**
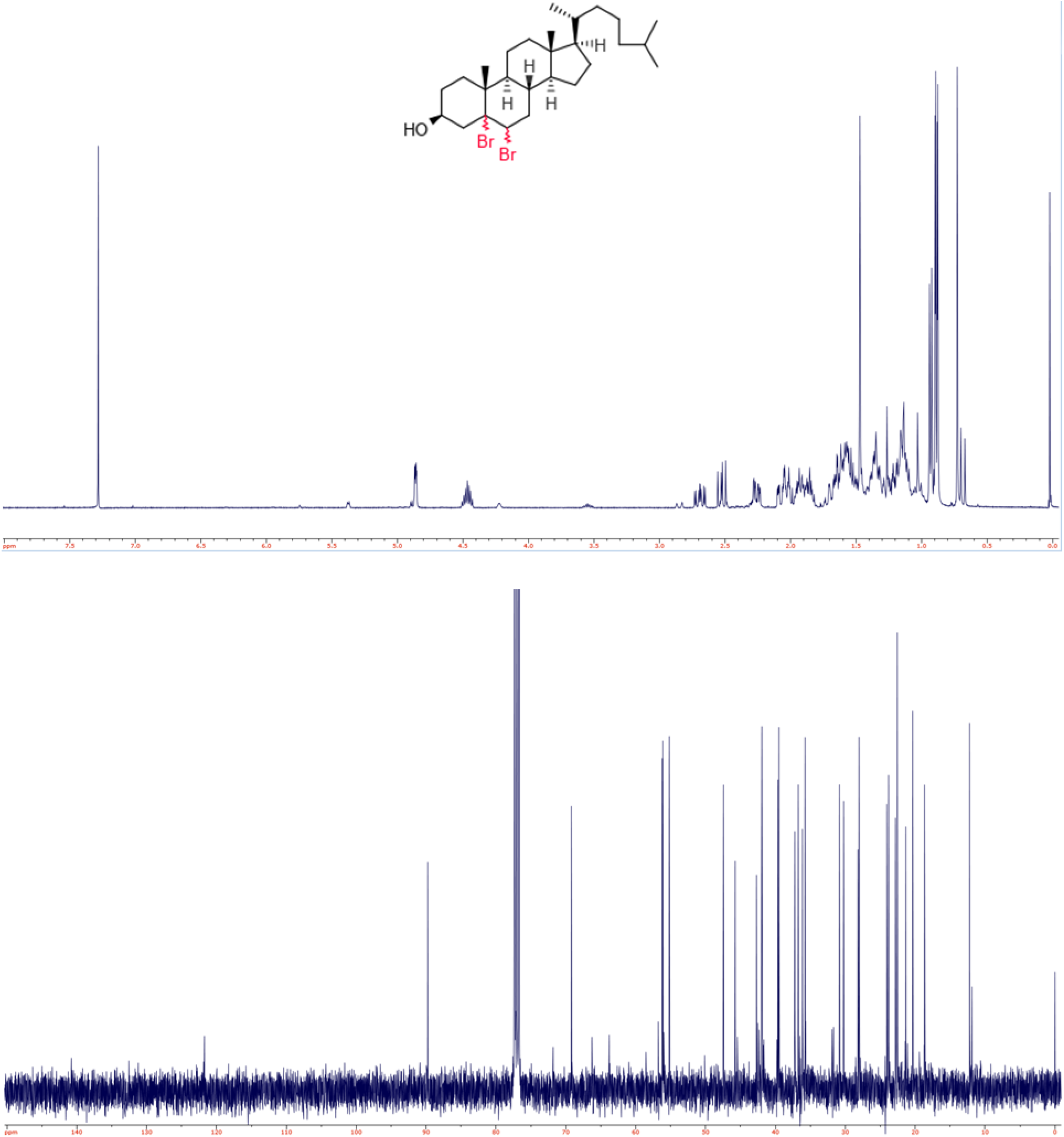
1H (top) and 13C (bottom) NMR spectra of CHOL-Br.

**Fig. S11.**
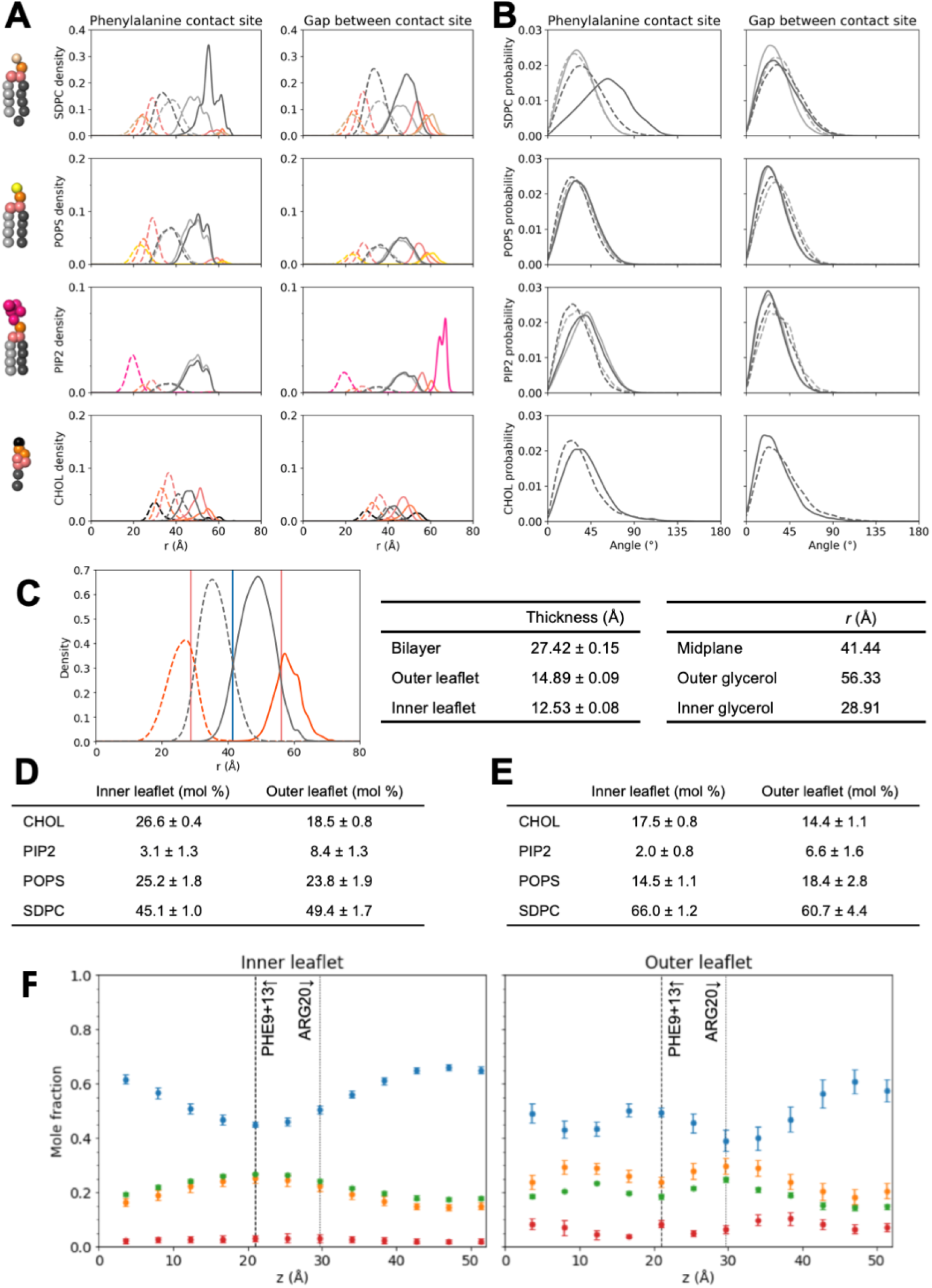
Lipid bilayer structure from CG-MD simulations with wild type CHMP1B-IST1 and lipid composition 1 simulations, averages of 10 independent replicates. A-B) Full radial density (A) and tail angle data (B) at the phenylalanine contact site (left panels) and the gap between the contact site (right panels). Radial density profiles correspond to different segments of the lipid, color coded as shown on the CG representations at the left. Phospholipid color coding: sn-1 tail (light grey), sn-2 tail (dark grey), glycerol (light pink), phosphate (orange), and headgroups (SDPC - beige, POPS - yellow, PIP2 - hot pink). Cholesterol color coding: hydroxyl (black), R1 and R2 ring beads (orange), R3-R5 ring beads (light pink), and aliphatic tail (dark grey). Outer leaflet densities are drawn as solid lines; inner leaflet densities are drawn as dashed lines. Note y-axis scale varies by lipid type due to abundance. At the phenylalanine contact site in the outer leaflet, phospholipids show a characteristic depletion in headgroup density, and the polyunsaturated tail of SDPC (dark grey, solid line) shifts outward radially. Tail angle distributions (B)corroborate this observation. Color scheme and line styles follow the same scheme as (A), and all distributions are normalized to 1. Outer leaflet lipid tails are more splayed at the phenylalanine contact site, sampling higher angles relative to the bilayer normal, especially the polyunsaturated tail of SDPC. C) Bilayer and leaflet thicknesses determined from mean radial lipid positions. Total radial densities of hydrophobic (dark grey curves) and hydrophilic (orange curves) lipid content show the overall bilayer structure, with inner leaflet densities in dashed lines and outer leaflet densities in solid. The bilayer midplane (blue vertical line) is given by the intersection of the hydrophobic distributions from the inner and outer leaflets. The outer and inner glycerol planes (pink vertical lines) are determined by the mean of the corresponding glycerol distributions. Bilayer and leaflet thicknesses are summarized in the center table, and mean locations of midplane and glycerol planes are in the right table. The inner leaflet is thinned relative to the outer leaflet, likely due in part to the overall increase in tail splay (B) of inner leaflet lipids compared to outer leaflet lipids, with the exception of those local to the phenylalanine contact site. D-E) Mean local compositions at the phenylalanine contact site (D) and at the gap between the contact site (E). The ± value is the standard deviation in mole fraction across the 10 independent replicates. F) Local compositions per leaflet per zone, plotted as the axial z position of the zone to correspond with Fig. 3H (main text). Each lipid is color coded: SDPC (blue), POPS (orange), CHOL (green), and PIP2 (red). Vertical dashed and dotted lines denote locations of membrane facing CHMP1B residues for reference. See Fig. S12 for zone definitions. Error bars represent ± 1 standard deviation between replicates. Strong variations in local composition are seen in each leaflet primarily for SDPC, because it is the majority component.

**Fig. S12.**
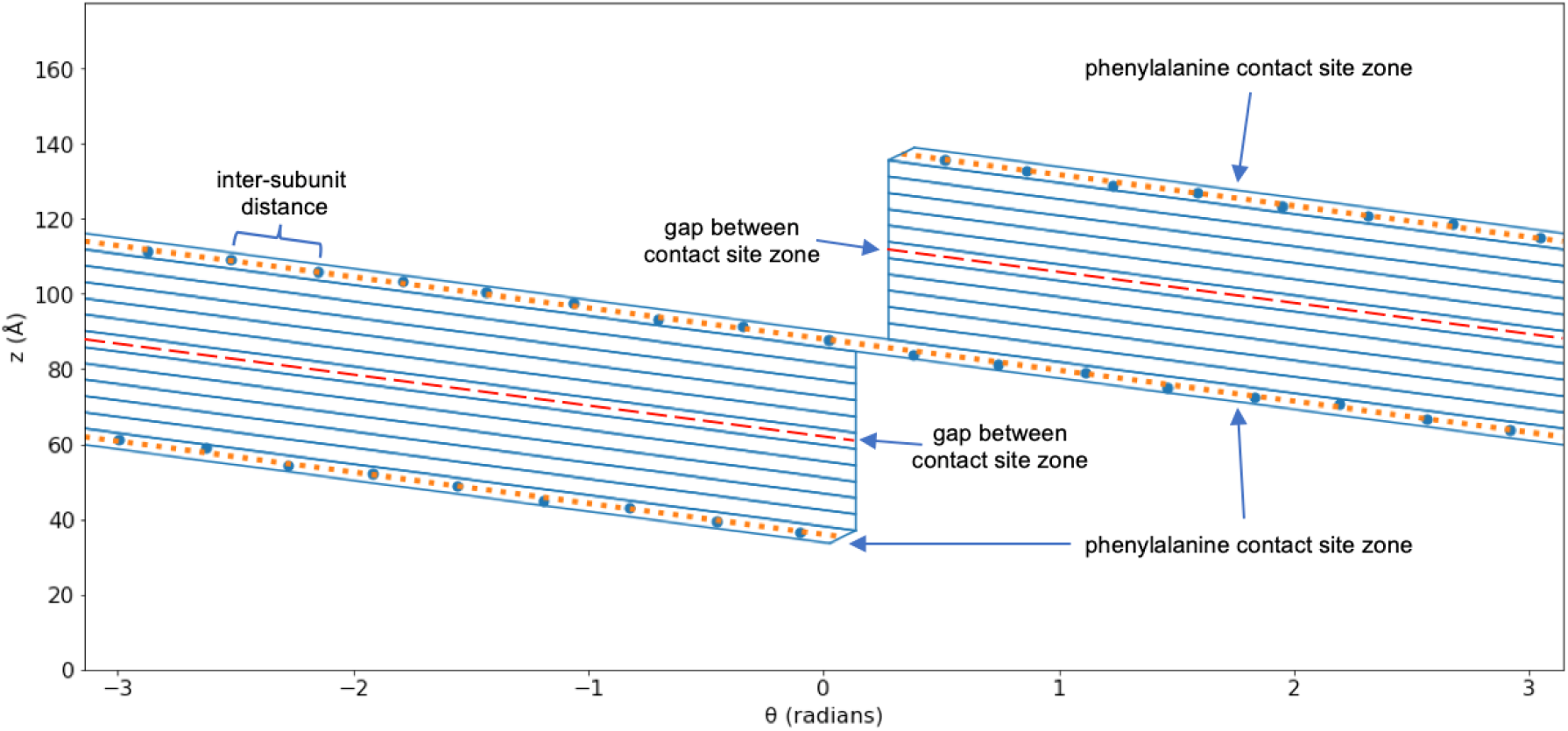
Visualization of (*θ*, *z*) zones used for simulation analysis. The line fit through mean coordinates of each F9 and F13 pair (large dots) defines the principal axis of each zone (shown as continuous spaces encompassed by solid blue lines). The “phenylalanine contact site” zone is the helical 30 ° wedge centered around the F9 and F13 fit line (dotted orange line), capped as described in Methods. All additional zones are successive 30 ° slices layered between the continuous phenylalanine contact zone, capped with vertical lines. The “gap between contact site” zone (wedge around dashed red line) is the zone 180 ° opposite the phenylalanine contact site, e.g. the central of the additional 11 zones (see Fig. 2).

**Fig. S13.**
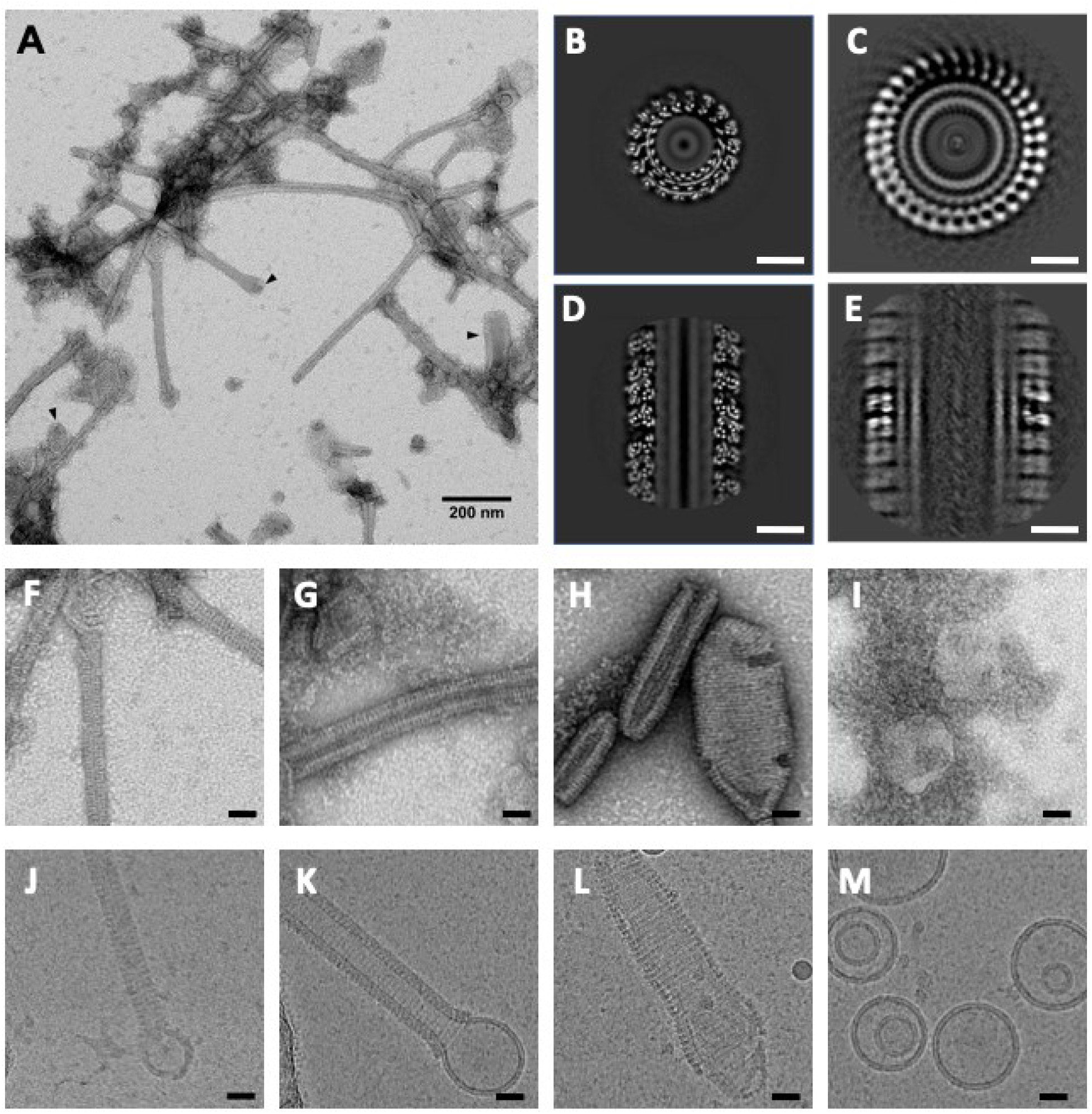
Negative stain TEM and cryo-EM of CHMP1B/IST1 filaments with different lipid compositions. A) Negative stain TEM micrograph of CHMP1B/IST1 filaments. Black arrows identify examples of vesicular remnants protruding from the ends of filaments, vesicles with no protein bound, or unusually wide filaments. B) Horizontal slice of a cryo-EM reconstruction of the 17 subunit per turn structure with lipid mixture 1 C) Vertical slice of a cryo-EM reconstruction of the 17 subunit per turn structure with lipid mixture 1. D) Horizontal slice of a cryo-EM reconstruction of the 34 subunit per turn structure with lipid composition 3. E) Vertical slice of a cryo-EM reconstruction of the 34 subunit per turn structure with lipid composition 3. Negative stain micrographs of F) Lipid composition 1. G) Lipid Mixture 3. H) Lipid composition 9. I) lipid composition 2. Cryo-EM micrographs of J) Lipid composition 1. K) Lipid composition 3. L) Lipid composition 9. M) Lipid composition 2. Scale bars represent 200 nm (A), 10 nm (B-E), and 25 nm (F-M).

**Fig. S14.**
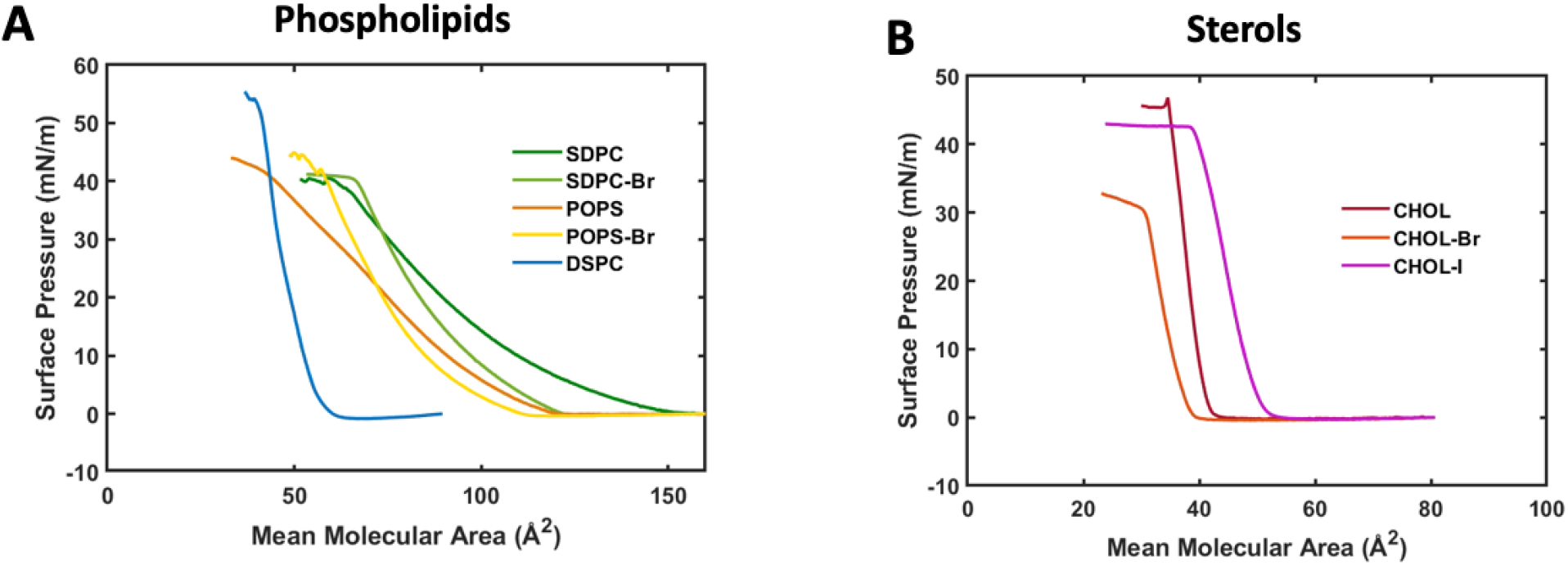
Biophysical characterization of brominated lipids. Langmuir pressure-area isotherms show that brominated phospholipids (A) and sterols (B) behave similarly to their unbrominated analogs in lipid monolayers. Although brominated lipids are less compressible than the unsaturated lipids, they are much more compressible than saturated lipids like DSPC. For both brominated lipids, the mean molecular areas (MMAs) are close to those of the analogous unsaturated lipids at 32 mN/m, the surface pressure that approximates the packing in a lipid bilayer (*65*). Furthermore, both brominated lipids display compressibility characteristic of lipids in the fluid phase at room temperature and similar to those of the analogous unsaturated lipids. The compressibility of the lipid monolayer is equal to the negative slope of the isotherm at a given surface pressure (in this case, 32 mN/m). In both cases, the brominated lipids are slightly less compressible than their unsaturated analogs. As a comparison, the isotherm of DSPC displays behavior typical of a saturated phospholipid with a transition temperature above room temperature. Its MMA and compressibility are much lower than those of the unsaturated and brominated lipids. See Moss, et al for more examples of pressure-area isotherms for a variety of lipid structures (5*8*). The pressure-area isotherms for CHOL-Br and CHOL-I are similar to that of CHOL. CHOL-Br has a slightly lower MMA at 32 mN/m than CHOL, while CHOL-I has a slightly larger MMA. The compressibility for all three sterols are similar. Small differences in MMAs also reflect the uncertainty in the lipid stock concentrations. Overall, these data demonstrate that in monolayers, the brominated lipids behave in a similar manner as the unsaturated lipids from which they are synthesized.

**Fig. S15.**
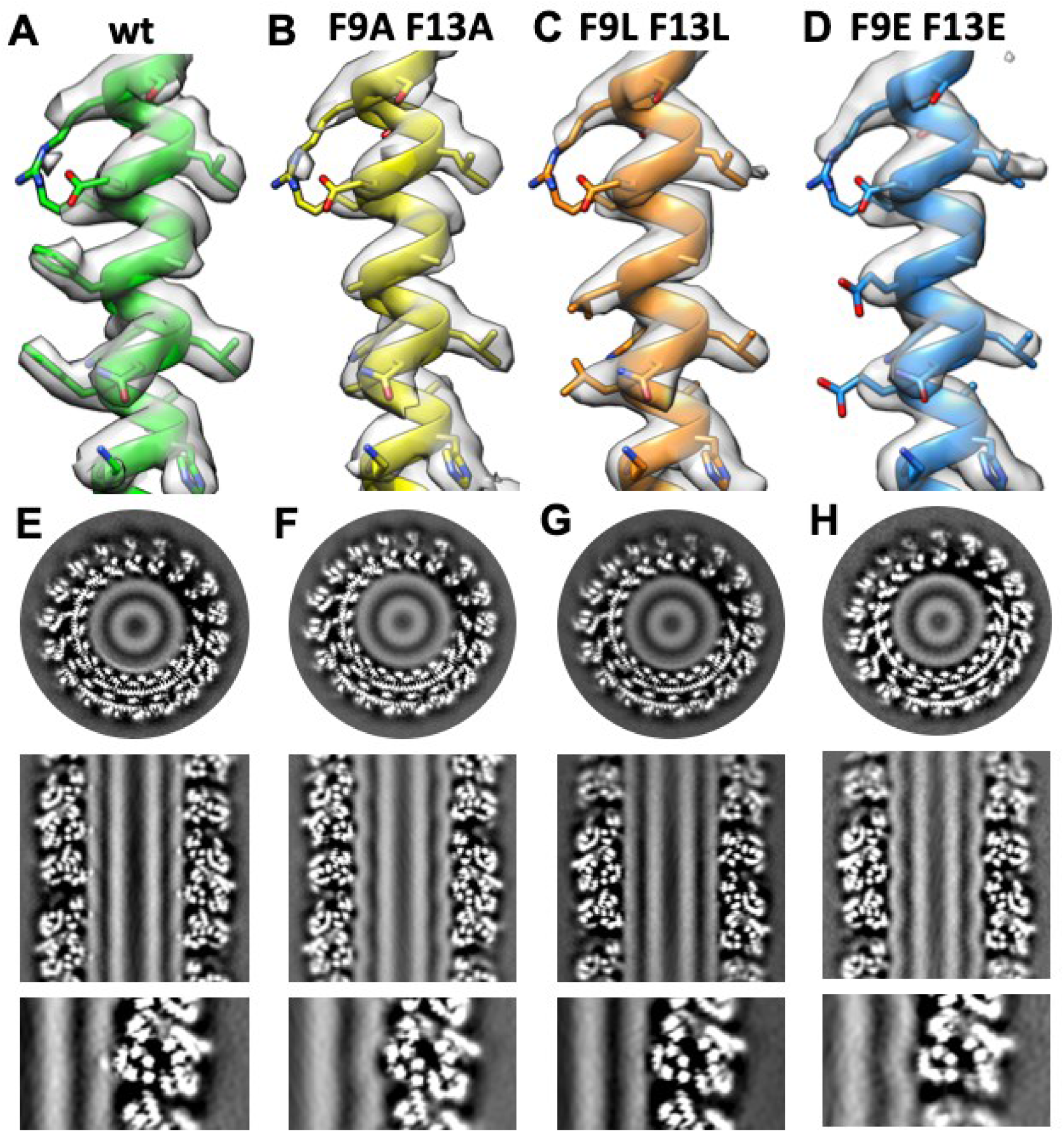
Cryo-EM of CHMP1B F9 and F13 double mutants with SDPC-Br (lipid composition 10). A-D) A section of CHMP1B helix α1 atomic model and cryo-EM density (gray) for wild type and double mutant proteins. E-H) Horizontal (top) and vertical (middle) slices through the cryo-EM densities for the wild type and CHMP1B double mutant proteins. The bottom panels show enlarged images of the membrane-contact sites for each sample. While the protein coat was identical to the wild type CHMP1B/IST1 coat (except for the mutated residues), the underlying lipid bilayer appeared quite different in each case. Counterintuitively, the smallest side chain, Ala, resulted in deep elastic deformations of the bilayer outer leaflet. The larger, charged Glu side chains had a similar effect on the bilayer. Interestingly, in both cases, no accumulation of SDPC-Br at the contact sites was observed. As discussed in the main text, we hypothesize that the SDPC-Br sn-2 tail does not interact with the small, less hydrophobic Ala and charged/polar Glu side chains, so it does not accumulate there. Without the formation of the hydrophobic defect at the bilayer surface characterized by a stripe of back-flipped lipid tails, the outer leaflet instead elastically deforms due to the presence of the charged CHMP1B helix α1. Lipid headgroup density is less depleted than for the wild type CHMP1B. Consistent with this model, substitution of the Phes for Leu, which is intermediate between Ala and Phe in both size and hydrophobicity, eliminates the elastic deformation in the outer leaflet and partially restores the SDPC-Br accumulation at the contact site. We hypothesize that such large changes in the structure of the bilayer will have consequences for the energy required to bend the bilayer and the energy necessary to cross the energy barrier to fission. In the absence of a cell-based assay for membrane remodeling by CHMP1B and IST1, these hypotheses remain difficult to test *in vivo*.

**Fig. S16.**
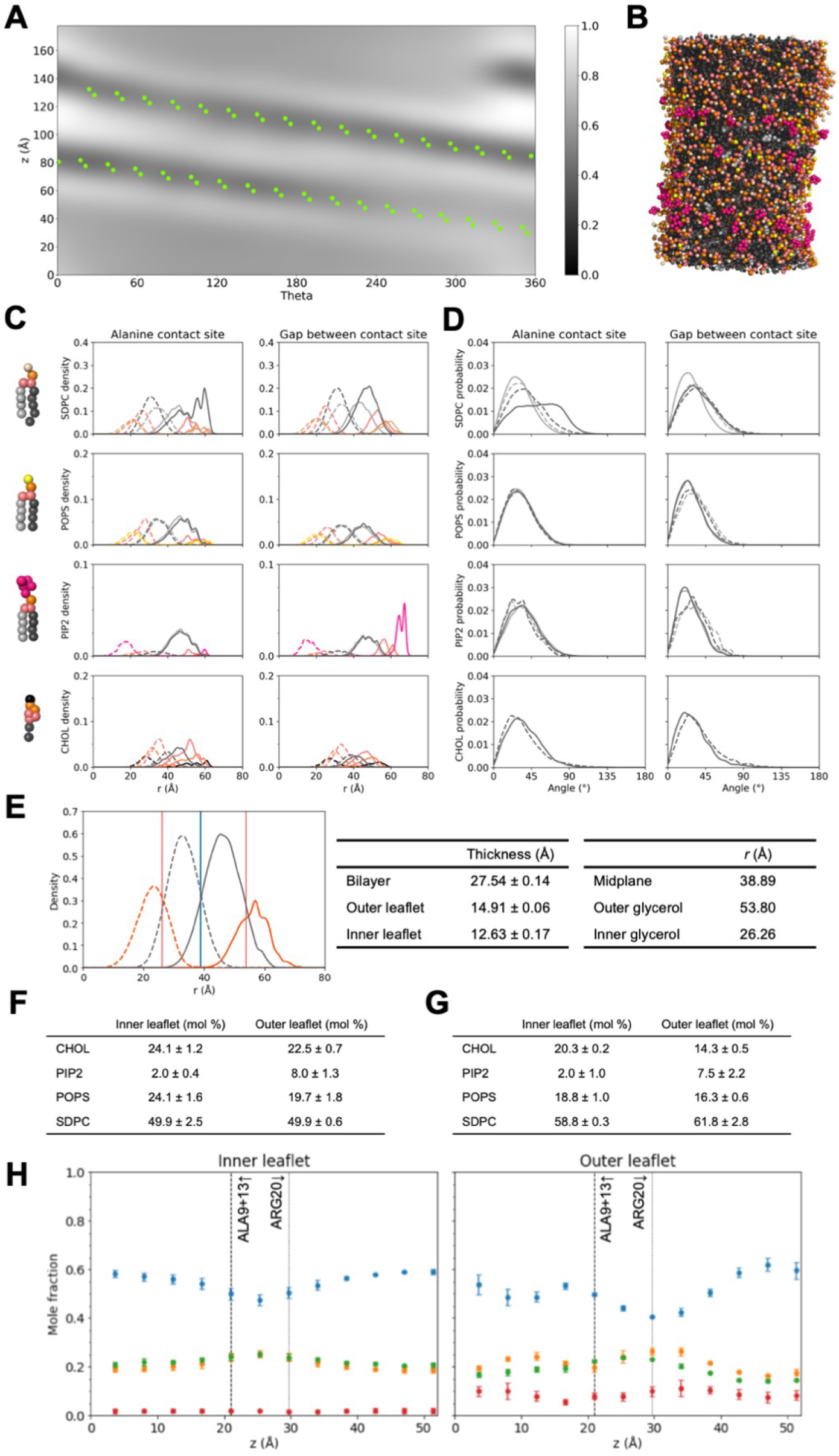
Lipid bilayer structure from CG-MD simulations with double mutant F9A + F13A CHMP1B and lipid composition 1, averages of three independent replicates. A) Average outer leaflet headgroup patterning represented as a two-dimensional (*θ*, *z*) density from three independent replicates. B) Representative snapshot of the outer tubule surface, with protein and solvent hidden (see Fig. S11 for color coding). The smaller alanine sidechains appear less able to maintain a persistent hydrophobic defect across the entire contact site, evidenced by weaker reductions in headgroup density at various segments compared to the WT case (main text Fig. 1h). C-D) Full radial density (C) and tail angle data (D) at the mutated alanine contact site (left panels) and the gap between the contact site (right panels). Radial density profiles correspond to different segments of the lipid, color coded as shown on the CG representations at the left. Phospholipid color coding: sn-1 tail (light grey), sn-2 tail (dark grey), glycerol (light pink), phosphate (orange), and headgroups (SDPC - beige, POPS - yellow, PIP2 - hot pink). Cholesterol color coding: hydroxyl (black), R1 and R2 ring beads (orange), R3-R5 ring beads (light pink), and aliphatic tail (dark grey). Outer leaflet densities are drawn as solid lines; inner leaflet densities are drawn as dashed lines. Note y-axis scale varies by lipid type due to abundance. At the mutated alanine contact site in the outer leaflet, phospholipid headgroups are pushed inward radially, consistent with a putative elastic deformation of the bilayer, while the polyunsaturated tail of SDPC (dark grey, solid line) displays reduced backflipping compared to the WT tubule (Fig. S11). Tail angle distributions (D) corroborate this observation. Color scheme and line styles follow the same scheme as (C), and all distributions are normalized to 1. Outer leaflet lipid tails are more splayed at the phenylalanine contact site, sampling higher angles relative to the bilayer normal, though the peak of the polyunsaturated SDPC tail is shifted to lower angles relative to its behavior in the WT tubule (Fig. S11). E) Bilayer and leaflet thicknesses determined from mean radial lipid positions. Total radial densities of hydrophobic (dark grey curves) and hydrophilic (orange curves) lipid content show the overall bilayer structure, with inner leaflet densities in dashed lines and outer leaflet densities in solid. The bilayer midplane (blue vertical line) is given by the intersection of the hydrophobic distributions from the inner and outer leaflets. The outer and inner glycerol planes (pink vertical lines) are determined by the mean of the corresponding glycerol distributions. Bilayer and leaflet thicknesses are summarized in the center table, and mean locations of midplane and glycerol planes are in the right table. The bilayer has nearly indistinguishable leaflet thickness compared to the WT composition 1 simulation (Fig. S11), though each plane is shifted to slightly lower values of *r*, perhaps reflective of the contact site membrane elastic deformation hypothesis vs. the hydrophobic defect of the WT system. F-G) Mean local compositions at the mutated alanine contact site (F) and at the gap between the contact site (G). The ± value is the standard deviation in mole fraction across the 10 independent replicates. H) Local compositions per leaflet per zone, plotted as the axial z position of the zone to correspond with Fig. 3H (main text). Each lipid is color coded: SDPC (blue), POPS (orange), CHOL (green), and PIP2 (red). Vertical dashed and dotted lines denote locations of membrane facing CHMP1B residues for reference. See Fig. S12 for zone definitions. Error bars represent ± 1 standard deviation between replicates. The variations in local composition appear to follow similar trends as in the WT composition 1 simulations (Fig. S11), though at smaller magnitude.

**Fig. S17.**
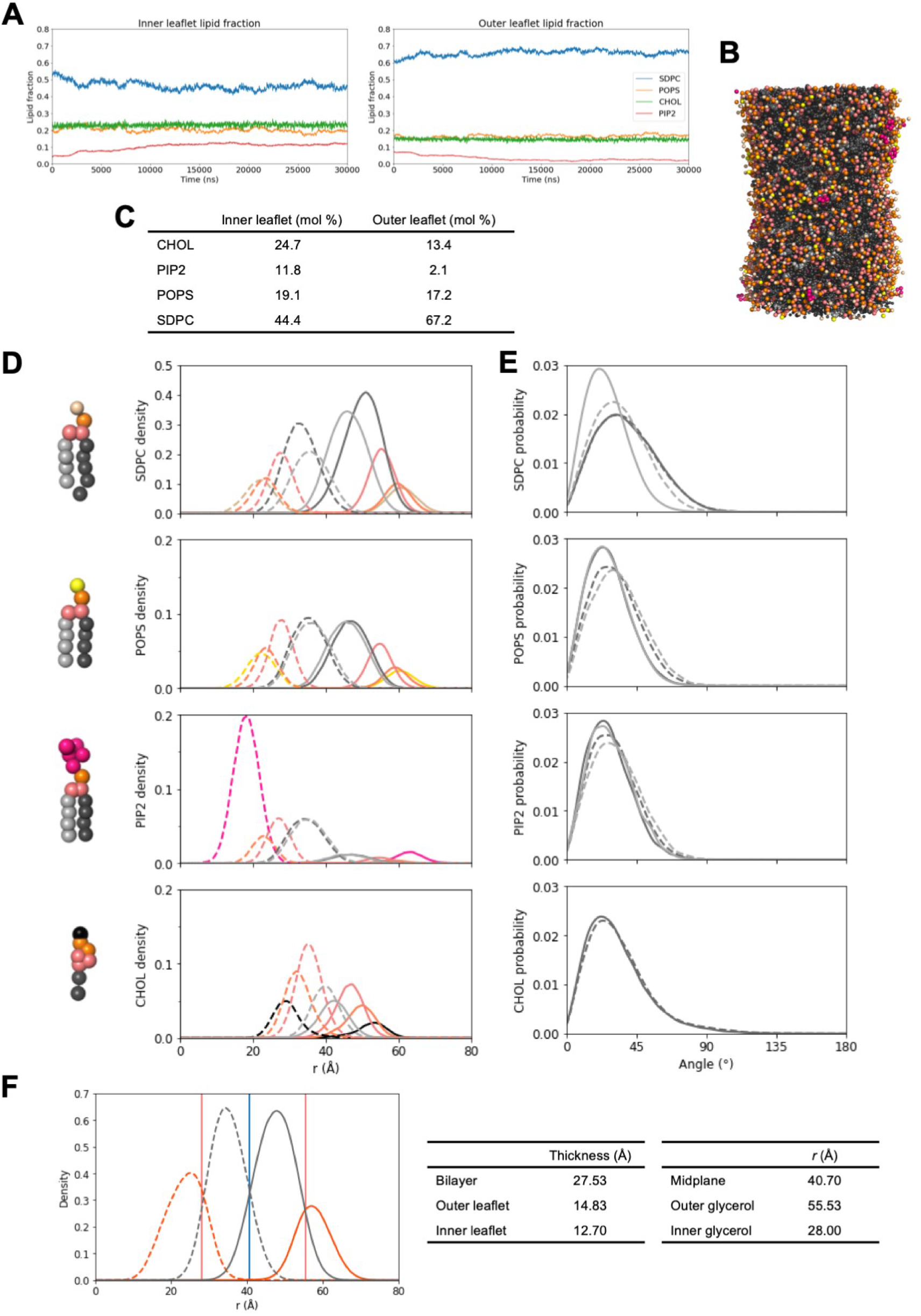
Protein-free lipid bilayer structure from CG-MD simulations of lipid composition 1. A) Compositions of the inner (left) and outer (right) leaflets over time during open-pore leaflet equilibration phase. When formed and equilibrated in the absence of protein even at similar overall dimension (F), there are strong differences in the preferential enrichment of some lipids, most notably PIP2. B) Snapshot of the outer tubule surface from the last frame from the production stage with solvent hidden (see Fig. S11 for color coding). C) Overall leaflet compositions calculated from production data. Relative to the WT CHMP1B-IST1 lipid composition 1 simulations (Table S8), there is stronger SDPC partitioning to the outer leaflet, stronger CHOL partitioning to the inner leaflet, and an inversion of the PIP2 partitioning now to the inner rather than the outer leaflet. D-E) Full radial density (D) and tail angle data (E). Radial density profiles correspond to different segments of the lipid, color coded as shown on the CG representations at the left. Phospholipid color coding: sn-1 tail (light grey), sn-2 tail (dark grey), glycerol (light pink), phosphate (orange), and headgroups (SDPC - beige, POPS - yellow, PIP2 - hot pink). Cholesterol color coding: hydroxyl (black), R1 and R2 ring beads (orange), R3-R5 ring beads (light pink), and aliphatic tail (dark grey). Outer leaflet densities are drawn as solid lines; inner leaflet densities are drawn as dashed lines. Note y-axis scale varies by lipid type due to abundance. In the absence of the protein, lipids are relatively normally oriented and there is no extreme backflipping even by the polyunsaturated tail of SDPC. That tail remains slightly shifted away from the bilayer center due to its flexibility but does not show substantial density past the phosphate peak (compare to Fig. S11). Tail angle distributions (E) reflect this as well. Color scheme and line styles follow the same scheme as (D), and all distributions are normalized to 1. F) Characterization of bilayer and leaflet thicknesses. At left, locations of the bilayer midplane (blue vertical line), outer and inner glycerol planes (pink vertical lines), and outer (solid curves) and inner leaflet (dashed curves) hydrophobic (dark gray) and hydrophilic (orange) radial densities. Bilayer and leaflet thicknesses are shown in the center table, and mean locations of the midplane and glycerol planes are at the right. Even without the protein to shape the formation of the tubule, it has similar dimensions to the other systems.

**Fig. S18.**
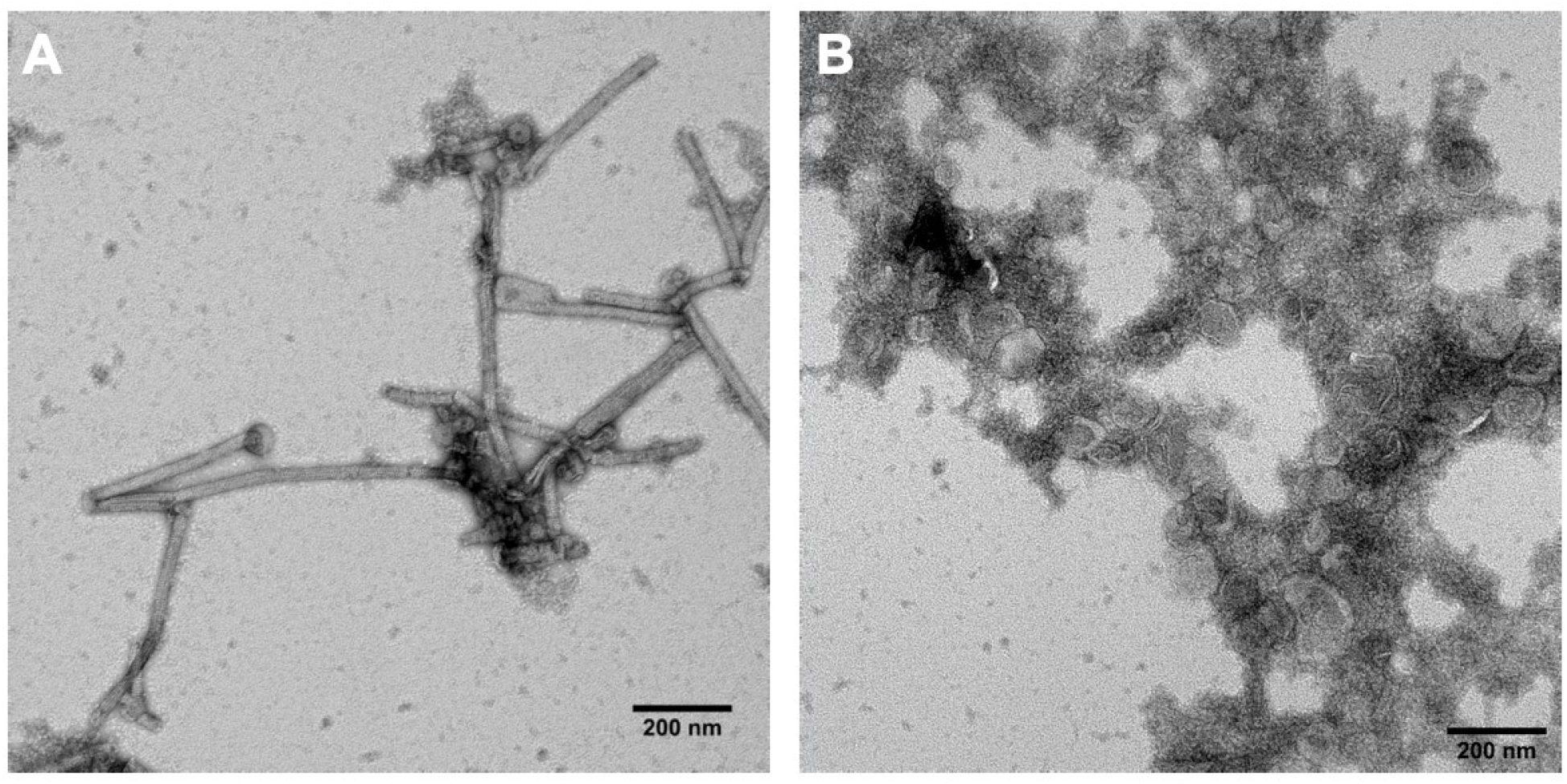
Negative stain TEM micrographs of CHMP1B/IST1 filaments. Negative stain EM was routinely performed to assess the formation and morphology of vesicles and membrane-bound CHMP1B/IST1 filaments. Vesicles and filaments formed with bromolipids were indistinguishable from those formed with unbrominated lipids. Fig. S13 shows a representative micrograph of CHMP1B/IST1 filaments with lipid composition 1 (Table 1), and Fig. S18 shows representative micrographs of CHMP1B/IST1 with lipid compositions 11 (A) and K16E/R20E CHMP1B and IST1 with lipid composition 1 (B). A) Wild type CHMP1B and IST1 with vesicles made from lipid composition 11 (Table 1). B) K16E/R20E CHMP1B and IST1 mixed with vesicles made from lipid composition 1 (Table 1). The double mutant protein does not bind to or remodel vesicles.

**Fig. S19.**
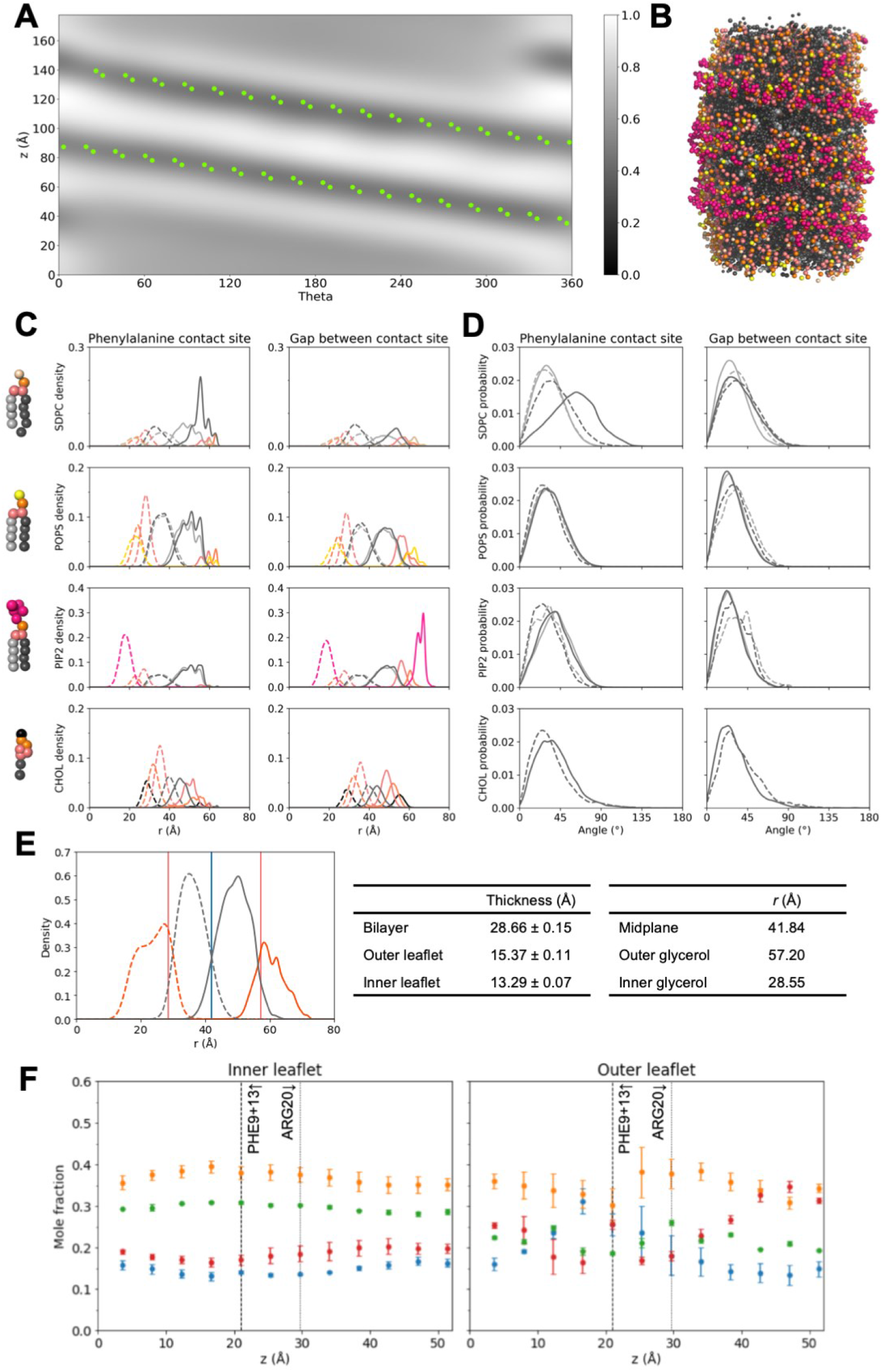
Lipid bilayer structure from CG-MD simulations with wild type CHMP1B-IST1 and lipid composition 17 simulations, averages of 3 independent replicates. A) Average outer leaflet headgroup patterning represented as a two-dimensional (*θ*,*z*) density from three independent 5 replicates. B) Representative snapshot of the outer tubule surface, with protein and solvent hidden (see Fig. S11 for color coding). The scale and persistence of the hydrophobic defect appear similar to the other WT simulations using Composition 1 (main text Fig. 1H). C-D) Full radial density (C) and tail angle data (D) at the phenylalanine contact site (left panels) and the gap between the contact site (right panels). Radial density profiles correspond to different segments of the lipid, color coded as shown on the CG representations at the left. Phospholipid color coding: sn-1 tail (light grey), sn-2 tail (dark grey), glycerol (light pink), phosphate (orange), and headgroups (SDPC - beige, POPS - yellow, PIP2 - hot pink). Cholesterol color coding: hydroxyl (black), R1 and R2 ring beads (orange), R3-R5 ring beads (light pink), and aliphatic tail (dark grey). Outer leaflet densities are drawn as solid lines; inner leaflet densities are drawn as dashed lines. Note y-axis scale varies by lipid type due to abundance. Main structural features seen in the WT protein + lipid composition 1 simulations (Fig. S11) are preserved despite the varied composition. Tail angle distributions (D) behave similarly as well. Color scheme and line styles follow the same scheme as (C), and all distributions are normalized to 1. E) Bilayer and leaflet thicknesses determined from mean radial lipid positions. Total radial densities of hydrophobic (dark grey curves) and hydrophilic (orange curves) lipid content show the overall bilayer structure, with inner leaflet densities in dashed lines and outer leaflet densities in solid. The bilayer midplane (blue vertical line) is given by the intersection of the hydrophobic distributions from the inner and outer leaflets. The outer and inner glycerol planes (pink vertical lines) are determined by the mean of the corresponding glycerol distributions. Bilayer and leaflet thicknesses are summarized in the center table, and mean locations of midplane and glycerol planes are in the right table. The leaflets similarly show differential thinning, and the bilayer is only marginally thicker than that of the WT composition 1 simulations (Fig. S11), perhaps indicating that the reduction in overall SDPC content for composition 17 produces a less flexible, thicker membrane. F) Local compositions per leaflet per zone, plotted as the axial z position of the zone to correspond with Fig. 3H (main text). Each lipid is color coded: SDPC (blue), POPS (orange), CHOL (green), and PIP2 (red). Vertical dashed and dotted lines denote locations of membrane facing CHMP1B residues for reference. See Fig. S12 for zone definitions. Error bars represent ± 1 standard deviation between replicates. Several trends in local composition are inverted relative to the WT composition 1 simulations (Fig. S11).

**Fig. S20.**
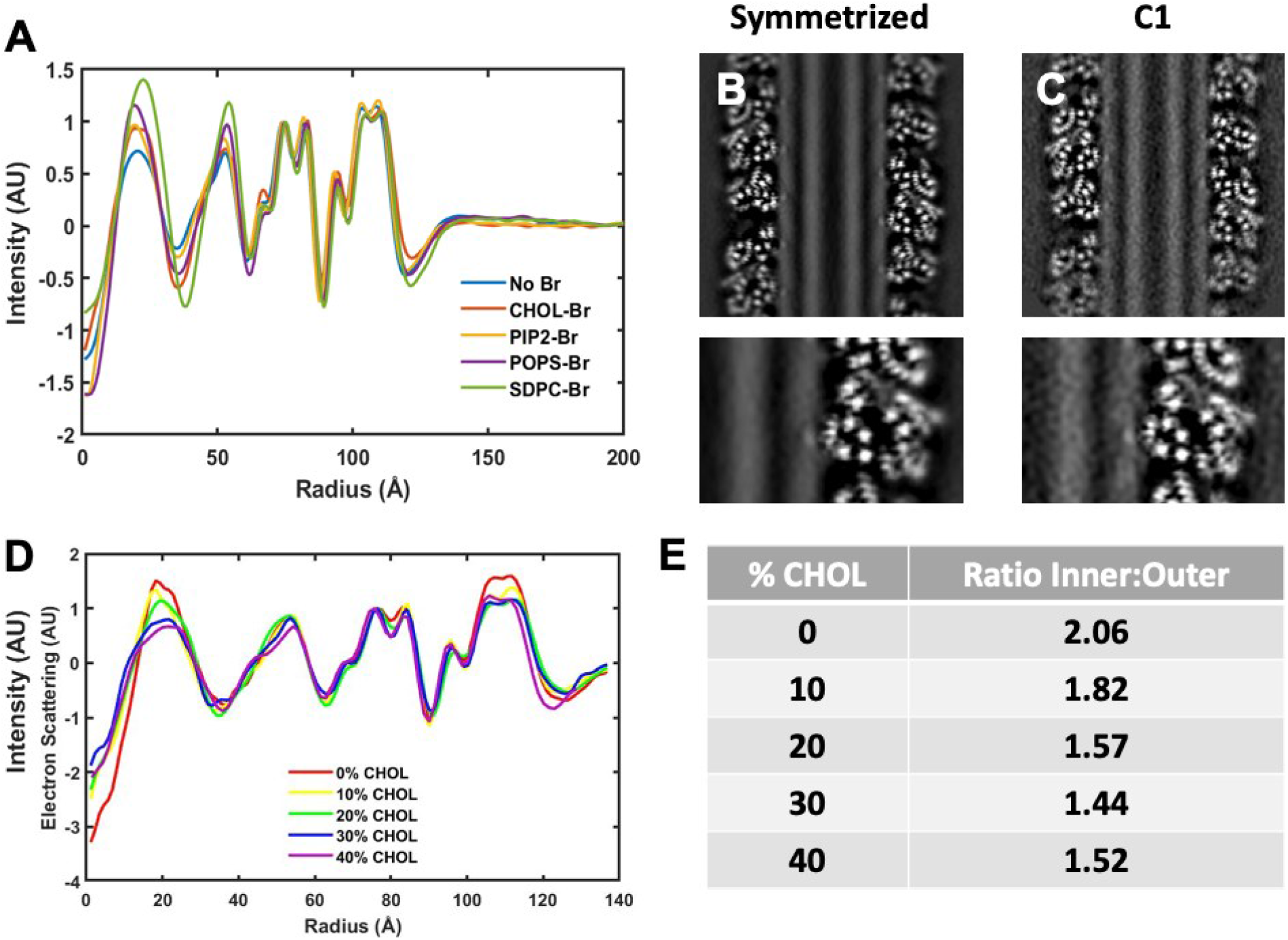
Comparison of cryo-EM reconstructions of CHMP1B/IST1 filaments with different lipid compositions. A) Normalized radial profiles of cryo-EM reconstructions with and without brominated lipids. The central 30 % of each cryo-EM reconstruction was projected along the helical axis, and the radial average was computed. B) Vertical slices through the cryo-EM reconstruction with SDPC-Br with helical symmetry applied and C) without helical symmetry applied. D) Radial profiles of cryo-EM reconstructions with different cholesterol concentrations show overall enrichment of cholesterol in the inner leaflet. E) Ratio of areas under each leaflet peak from (D) show a decrease in the intensity of the inner leaflet peak as the cholesterol concentration increases, consistent with cholesterol’s lack of electron dense phosphates.

**Fig. S21.**
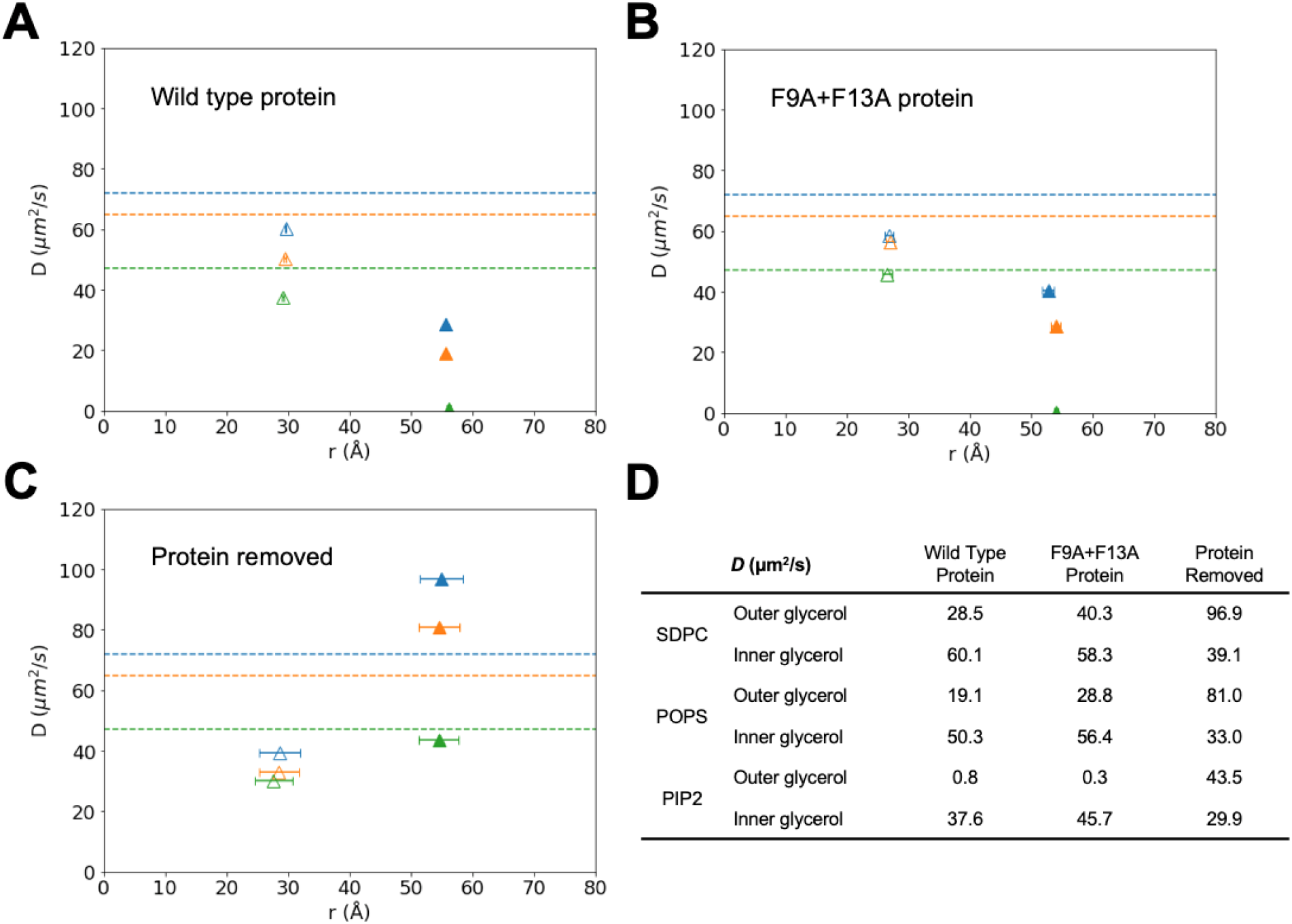
Lipid diffusion rates from CG-MD simulations of WT and F9A + F13A CHMP1B with lipid composition 1. A-C) Phospholipid diffusion rates in the WT (A), double mutant F9A + F13A (B), and protein free (C) simulations plotted against the mean radial coordinate of the lipid’s second glycerol bead (GL2) used for the calculation. For reference, diffusion rates calculated in a flat bilayer are represented as horizontal dashed lines. Lipid color coding: SDPC (blue), POPS in orange, and PIP2 in green. Inner leaflet values are shown with hollow markers and outer leaflet markers are filled. Horizontal error bars at left are ± 1 standard deviation of the mean time-averaged radial coordinate across ten trajectories, and at right are ± 1 standard deviation of radial coordinates across time in the protein-free trajectory. D) Summary of graph values. Diffusion rates in the presence of protein are similar between the WT and mutant CHMP1B simulations, with outer leaflet lipids—especially PIP2—showing dramatically slowed diffusion rates relative to inner leaflet lipids. This reduction in diffusion is likely due to a combination of electrostatic interactions with positively charged CHMP1B residues, which transiently pin charged lipids like PIP2, and the headgroup exclusion region at the F9 + F13 contact site (Fig. 1H), which largely prevents lipids from crossing the helical furrow formed by these residues. The slight increase in diffusion for outer leaflet SDPC and POPS in the mutant F9A + F13A simulations may reflect the lessened headgroup exclusion (Fig. S16) in this system. Contrastingly, with the protein removed each lipid shows faster diffusion in the outer leaflet, as is expected from the tubule geometry (64). We expect that the reduction in inner leaflet diffusion rates for the no-protein case relative to the with-protein simulations is attributable to the higher amounts of saturated lipid tails in the inner leaflet of the no-protein simulations vs. the with-protein simulations (Fig. S17C vs. SI Tables 8, 9), which would lower diffusion rates. The much faster outer leaflet diffusion seen in the absence of protein suggests an overall friction-like effect from the protein acting on the tubule. Finally, the increased diffusion rates in the outer leaflet of the protein-free tubule relative to the flat bilayer suggests that the increase in area per lipid concomitantly increases diffusion rates.

**Fig. S22.**
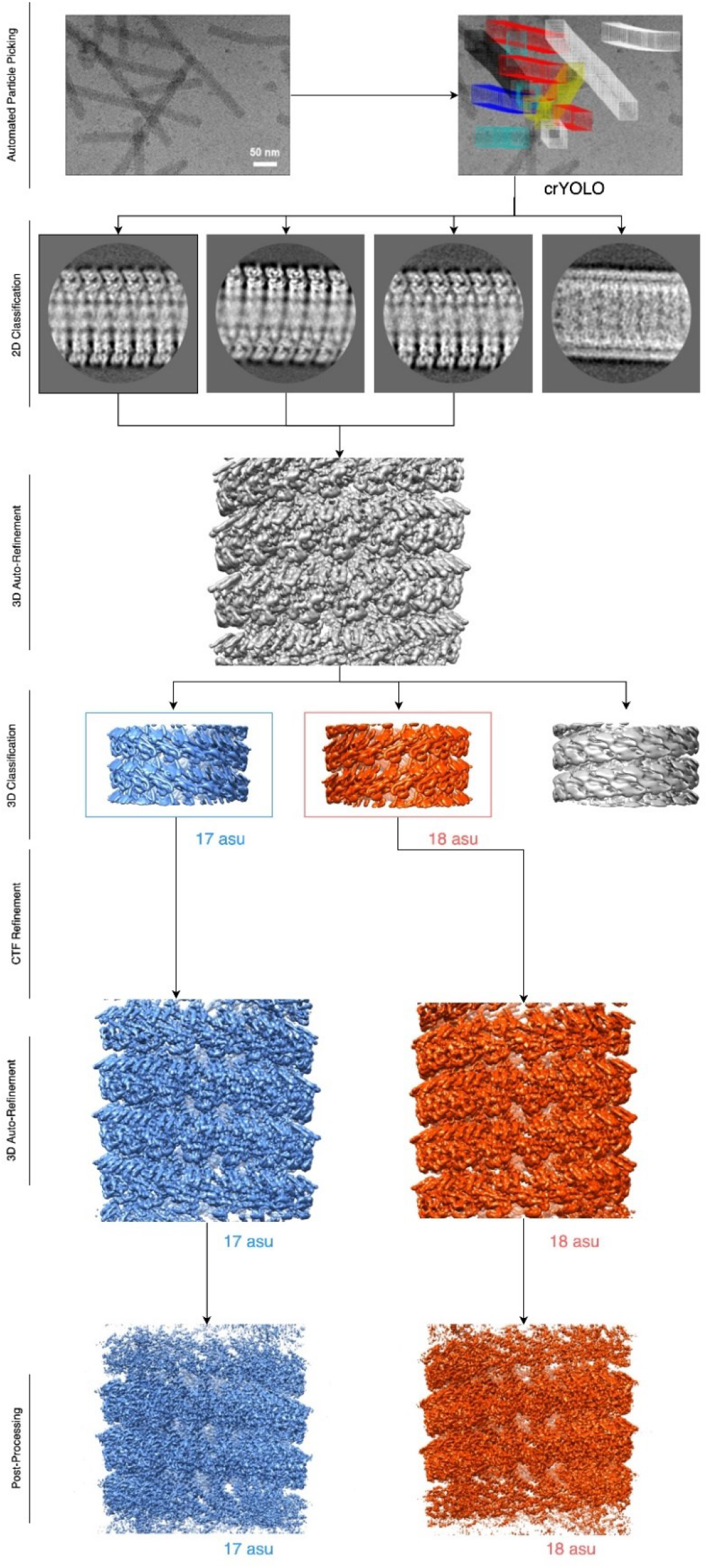
Cryo-EM data processing pipeline. All steps were performed in Relion 3.08 except particle picking, which was performed with SPHIRE-crYOLO.

**Fig. S23.**
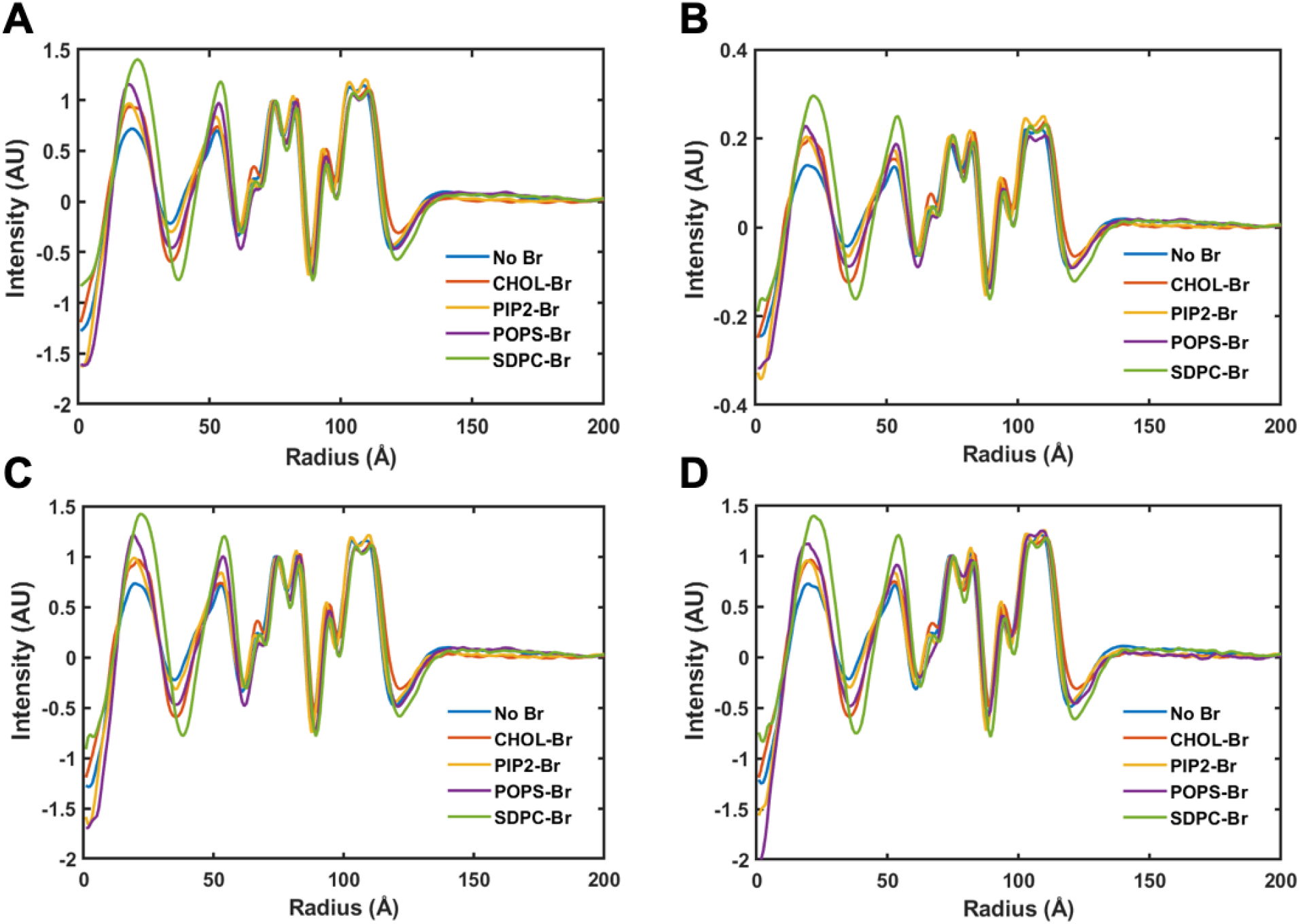
Effects of data processing on electron scattering profiles. A) Normalized electron scattering profiles with helical symmetry and 6 Å low pass filter applied. B) Unnormalized electron scattering profiles with helical symmetry and 6 Å low pass filter applied. C) Normalized electron scattering profiles with helical symmetry applied and no low pass filter applied. D) Normalized electron scattering profiles without helical symmetry or low pass filter applied. All electron scattering profiles are from radial averages of Z-projected density maps.

**Figure S24.**
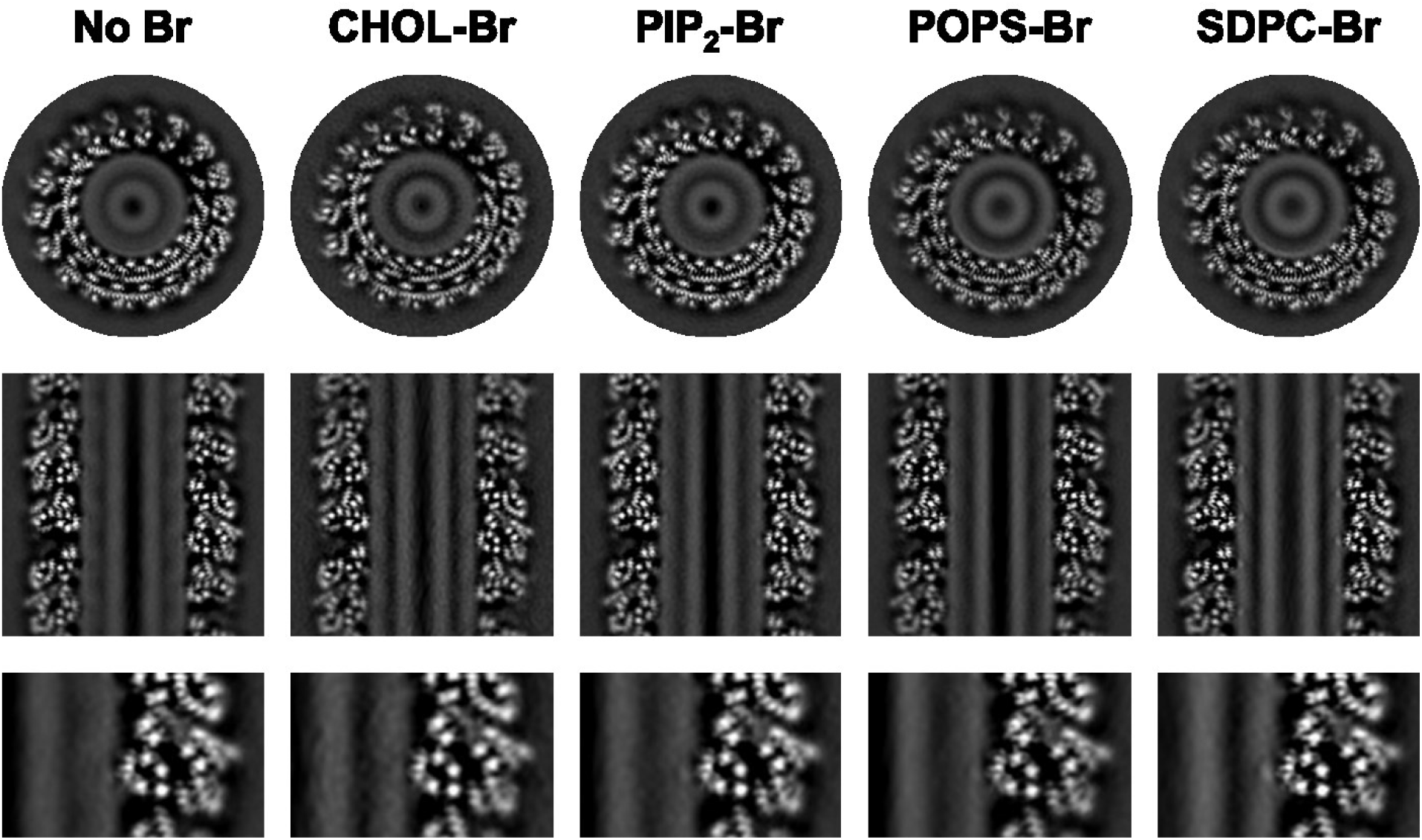
18 subunit per turn reconstructions. Horizontal (top) and vertical (middle) slices through the cryo-EM reconstructions of 18 subunit per turn CHMP1B/IST1 filaments with and without brominated lipids. The bottom panels show an enlarged view of the membrane-protein contact sites. Each of the brominated lipids resulted in cryo-EM reconstructions with different patterns of intensity in the lipid bilayer. For example, CHOL-Br appears to uniformly increase the intensity of the bilayer, while the phospholipids tend to segregate asymmetrically. The phospholipids increase the intensity of each leaflet of the bilayer by different amounts and accumulate to variable extents where CHMP1B contacts the bilayer. These qualitative observations are quantified in Fig. 3 and Fig. 5.

**Fig. S25.**
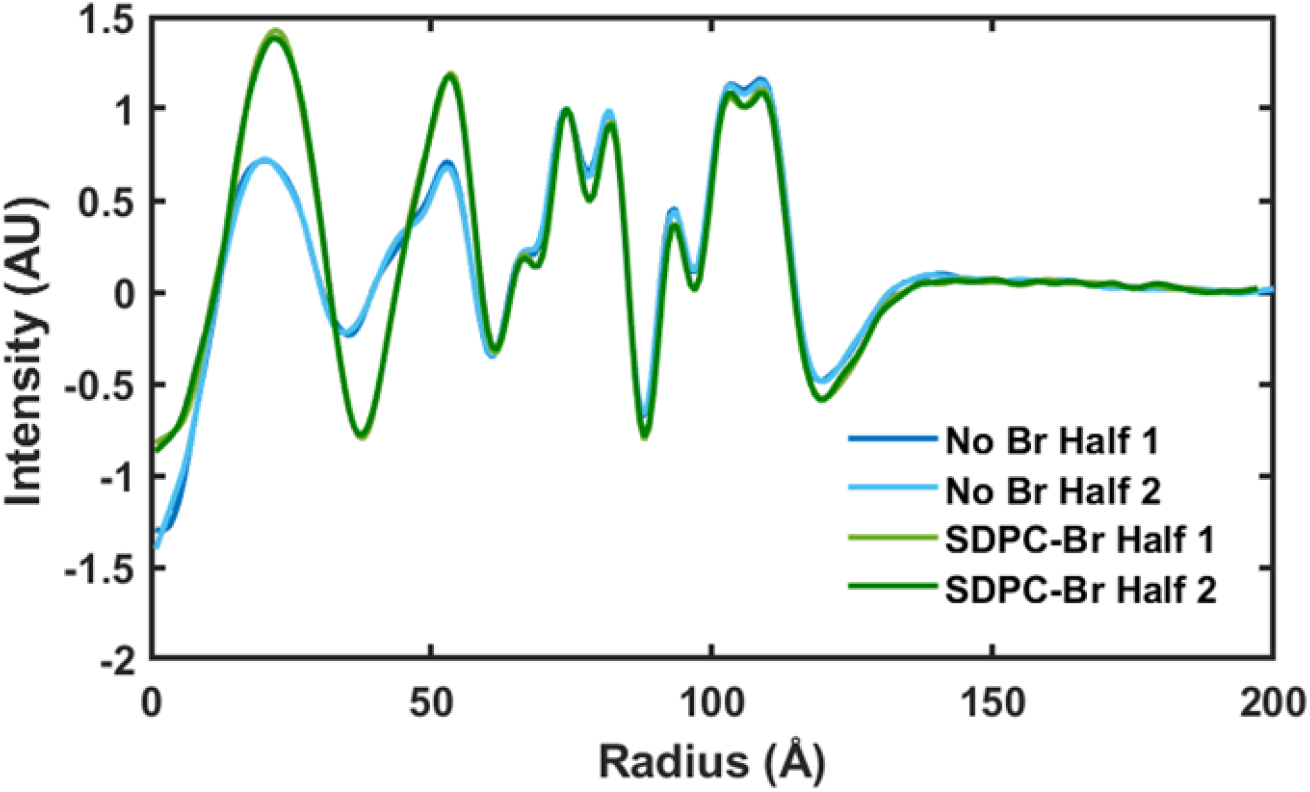
Comparison of half maps for the reconstructions with unbrominated lipids and SDPC-Br (Table 1, lipid compositions 1 and 10).

**Fig. S26.**
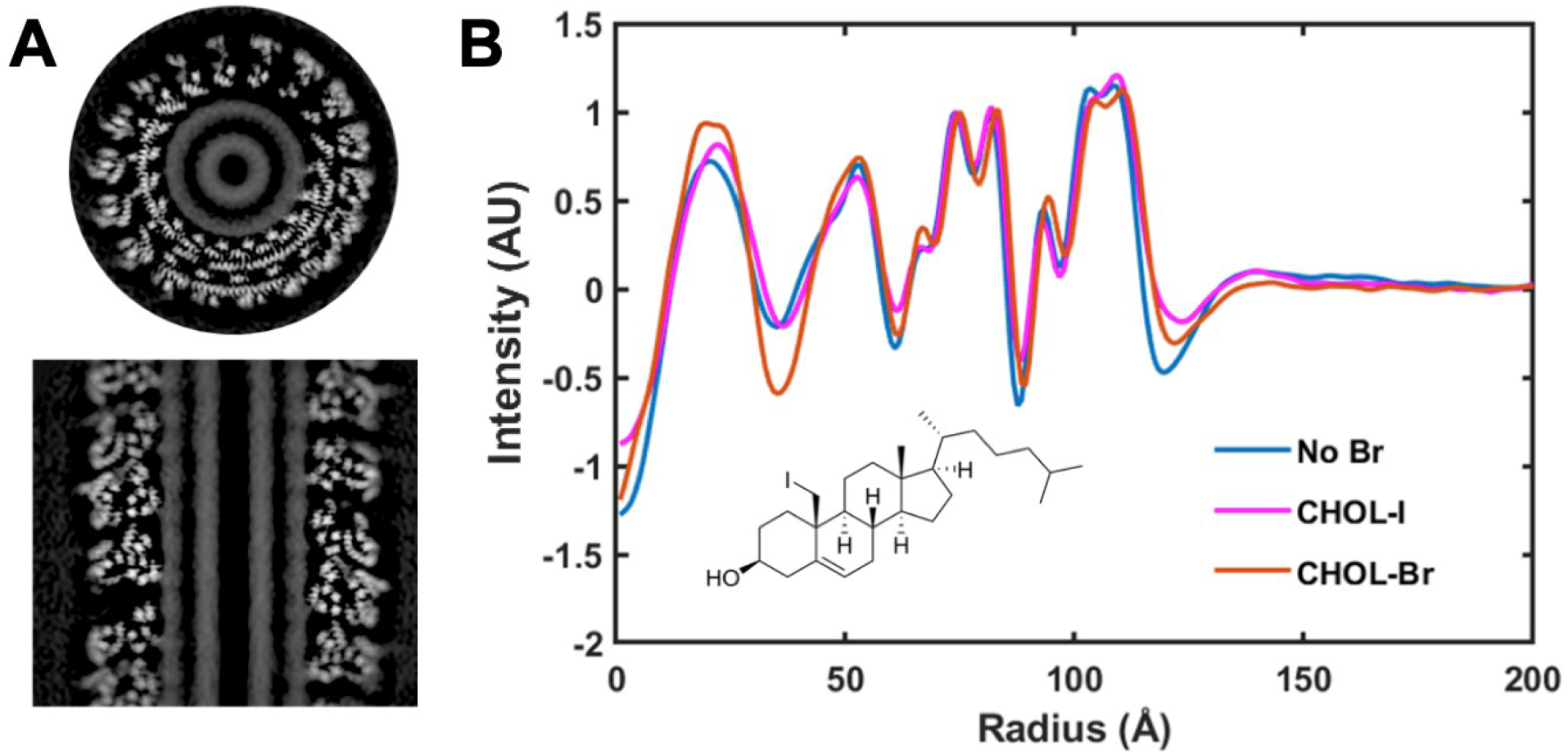
Reconstructions and radial profiles with CHOL-I and comparison to CHOL-Br. A) Horizontal (top) and vertical (bottom) slice through cryo-EM reconstruction with CHOL-I. B) Comparison of radial profiles of cryo-EM reconstructions from samples with no brominated lipids, CHOL-I, and CHOL-Br. Samples with CHOL-I appeared qualitatively similar to those containing CHOL-Br, albeit with less added intensity, as expected based on the electron atomic scattering factors for two bromine atoms versus one iodine atom. As the iodine atom is in a different location than the bromine atoms and should be less perturbative than two bromine atoms, the similar behavior compared to both CHOL and CHOL-Br supports the idea that halogenation of cholesterol is minimally perturbative to its behavior in lipid bilayers.

**Fig. S27.**
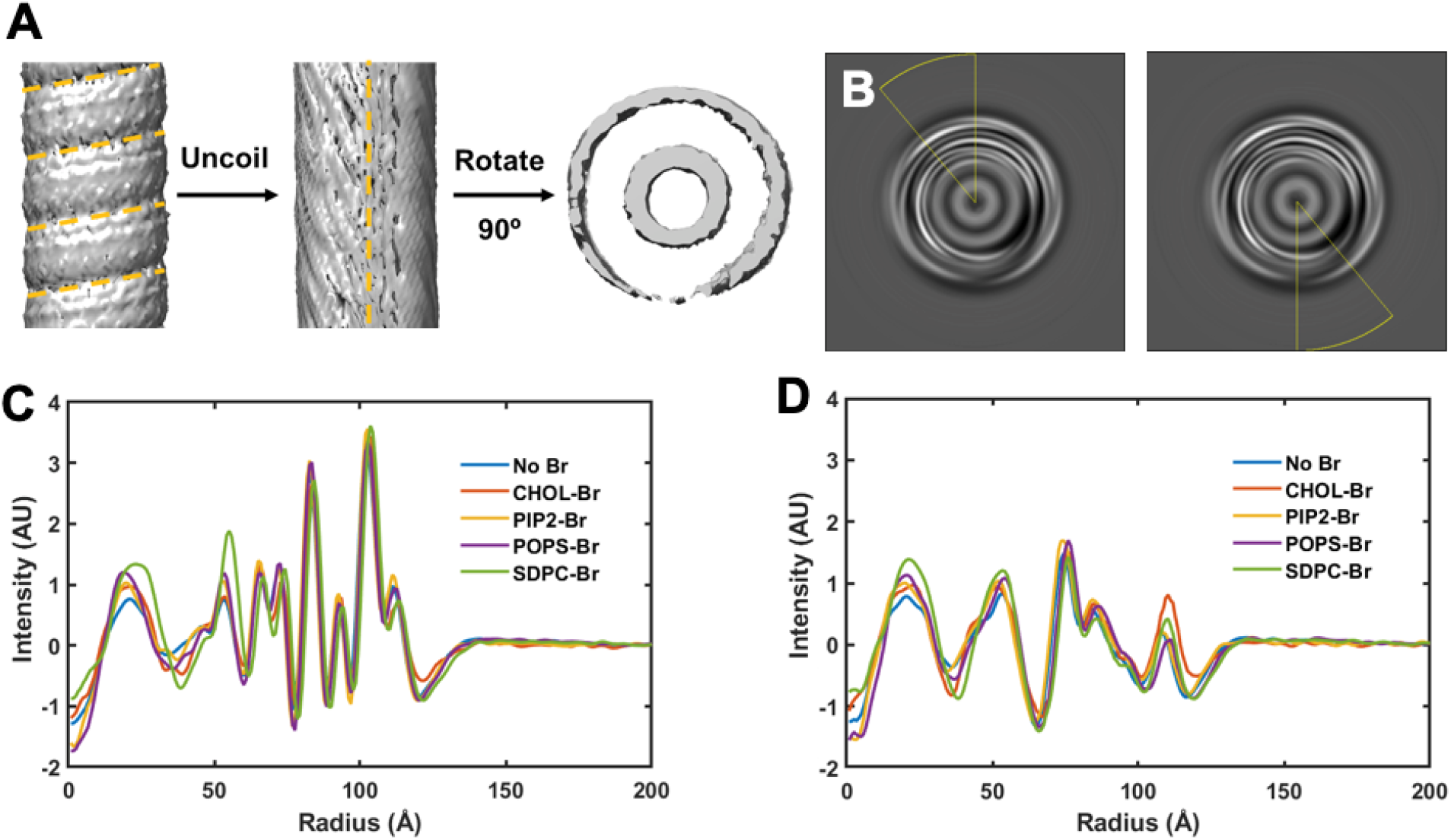
Compositional analysis of lipid bilayers in cryo-EM reconstructions. A) Analysis of the structure and composition of the membrane-protein contact sites and the gaps between contact sites. To improve the signal to noise ratio, each Z slice of the cryo-EM reconstruction was rotated to align it with the previous frame, resulting in an “uncoiled” nanotube that was projected in Z. B) Radial averages for 30 ° at the contact sites (left) and between the contact sites (right), where the protein is farthest from the bilayer. Radial averages for cryo-EM reconstructions at the contact sites (C) and at the gaps between contact sites (D).

**Fig. S28.**
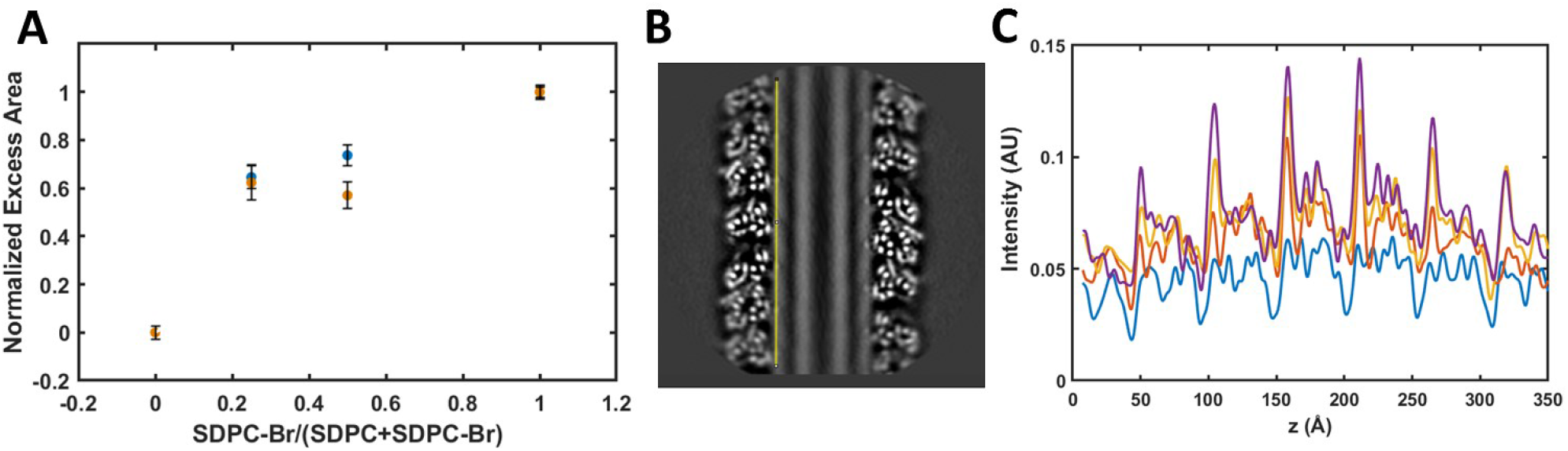
Sub-stoichiometric titration of lipid tubules by SDPC-Br vs. SDPC. A) Quantification of the excess intensity in each leaflet, with normalization per leaflet. Reducing the SDPC-Br concentration results in attenuated Br scattering in each leaflet. The reduction is not linear with SDPC-Br concentration, suggesting that SDPC-Br is somewhat enriched in the lipid bilayer nanotubes relative to SDPC. B) Visualization of the SDPC-Br titration along a Z line scan of the outer leaflet from vertical slices of the cryo-EM reconstructions (C).

**Fig. S29.**
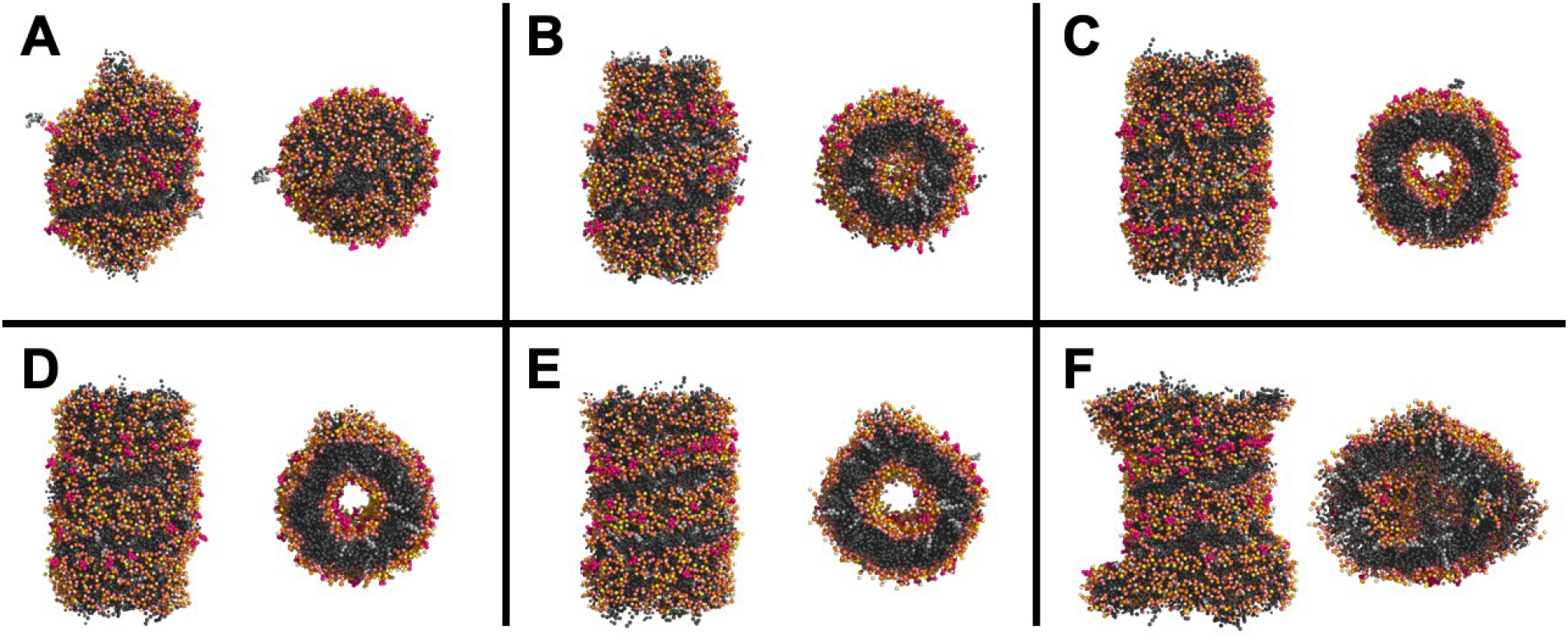
Optimization of lipid density in CG-MD simulations. A-F) Final frame of tubule from side (left) and top-down (right) from simulations tuning total lipid count. Solvent and protein removed for clarity. Lipid composition was composition 1, with total lipid amounts A) 1017, B) 1187, C) 1271, D) 1314, E) 1356, and F) 1695. Cholesterol in light gray, lipid tails in dark gray, glycerol in light pink, phosphates in orange, SDPC headgroups in beige, POPS headgroups in yellow, and PIP2 headgroups in bright pink. Insufficient lipids result in vesicle formation (A) or tubule with varying degrees of neck narrowing away from the protein (B-C); whereas an over-packed tubule bulges outward away from the protein (E) or fills the tubule lumen with lipids (F). A lipid count of 1300 (D) was used for production simulations since this resulted in a consistent tubule radius along the z axis with a clearly defined and solvated tubule lumen. These simulations employed a protein coat (PDB ID 3JC1) with opposite handedness to the coat used for all simulations reported in the main text (PDB ID 6TZ5), but nearly identical dimensions. These simulations used an isotropic Berendsen barostat for pressure coupling and were run with a 0.03 ps timestep for 6 μs, while all other parameters were identical to those described elsewhere in Methods for Simulation sets 1 & 2.

**Fig. S30.**
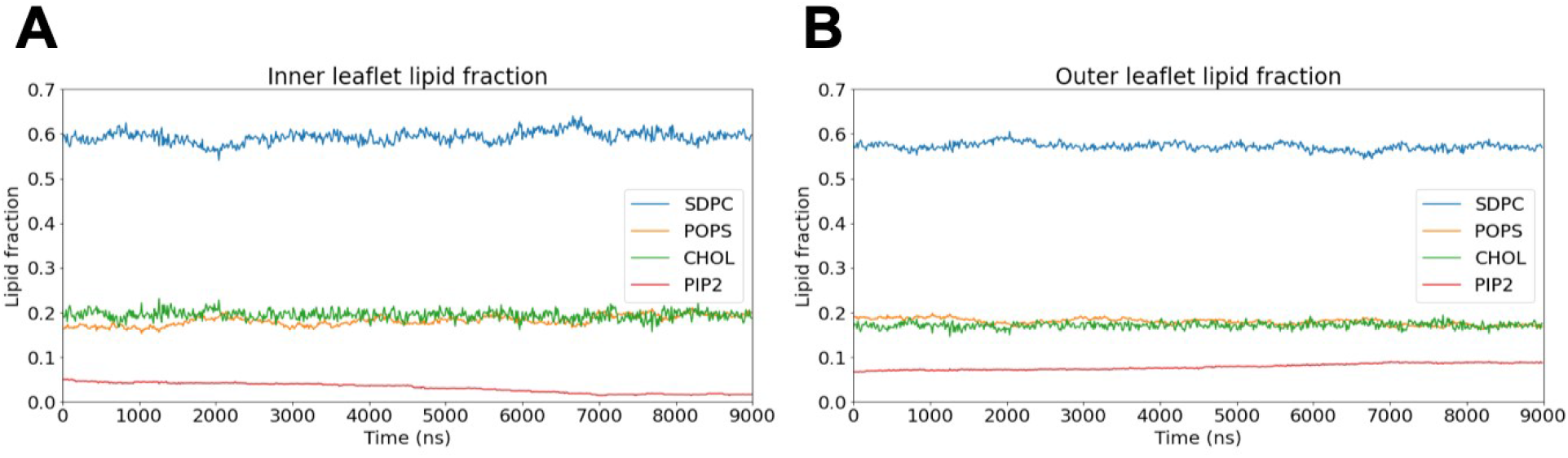
Leaflet compositions over time from one replicate of WT CHMP1B-IST1 composition 1 simulation during leaflet equilibration (procedure described in Methods, Simulation sets 1 & 2 leaflet equilibration). Leaflet assignment was determined using a simple *r* cutoff (r < 42 Å inner, else outer) applied to the lipid center of mass, since the clustering algorithm performed poorly in the presence of the equilibration pore due to the connection between the two leaflets. PIP2 was found to typically require the longest time to equilibrate, likely due to its low total number in the lipid mix and strong electrostatic attraction to the protein coat. Equilibration was run for 9 μs for composition 1 simulations (Simulation sets 1 & 2), and 6 μs for composition 17 simulations (Simulation set 3), as composition 17 replicates more quickly converged to similar leaflet compositions (Table S9).

**Fig. S31.**
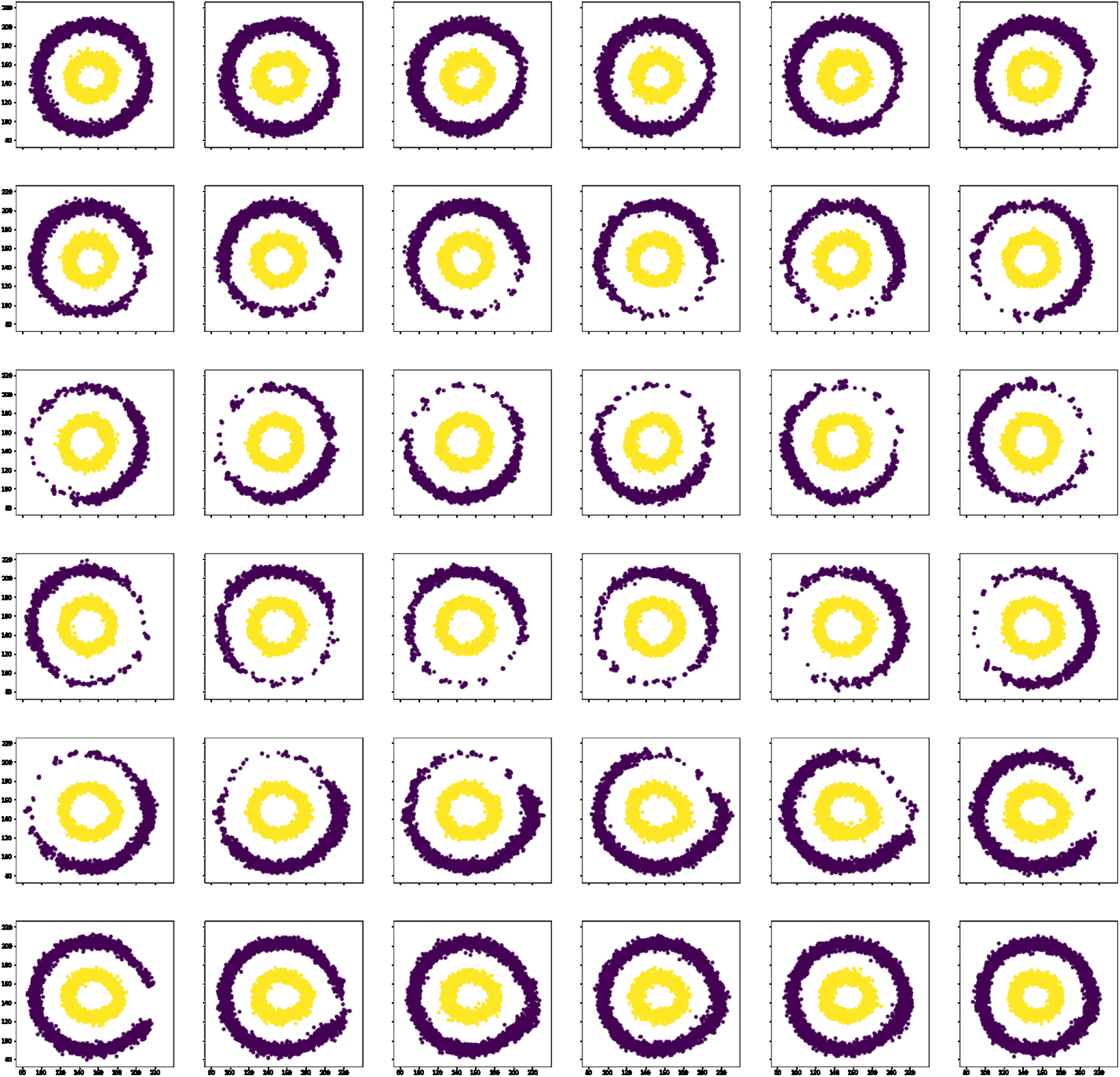
Assignment of lipids to leaflets in simulation. Cartesian (*x*,*y*) coordinates of SDPC phosphate beads from all analyzed frames of one replicate of WT CHMP1B-IST1 Composition 1 production simulation (Simulation set 1), sorted into 5 Å bins in *z* scanning from the bottom of the box (upper left) to the top of the box (lower right), going through subplots left to right in columns and then down in rows. Each SDPC is colored according to the assigned leaflet, dark spots are outer leaflet and yellow spots are inner leaflet. The applied clustering (see Methods section Simulation Analysis) produces cleanly separated leaflets. Scanning through z shows partial exclusion of lipid phosphates spiraling along the helix corresponding to repeats of CHMP1B F9 and F13, and an unperturbed outer leaflet away from the protein at the bottom (upper left few panels) and top (bottom right few panels) of the simulation box.

**Fig. S32.**
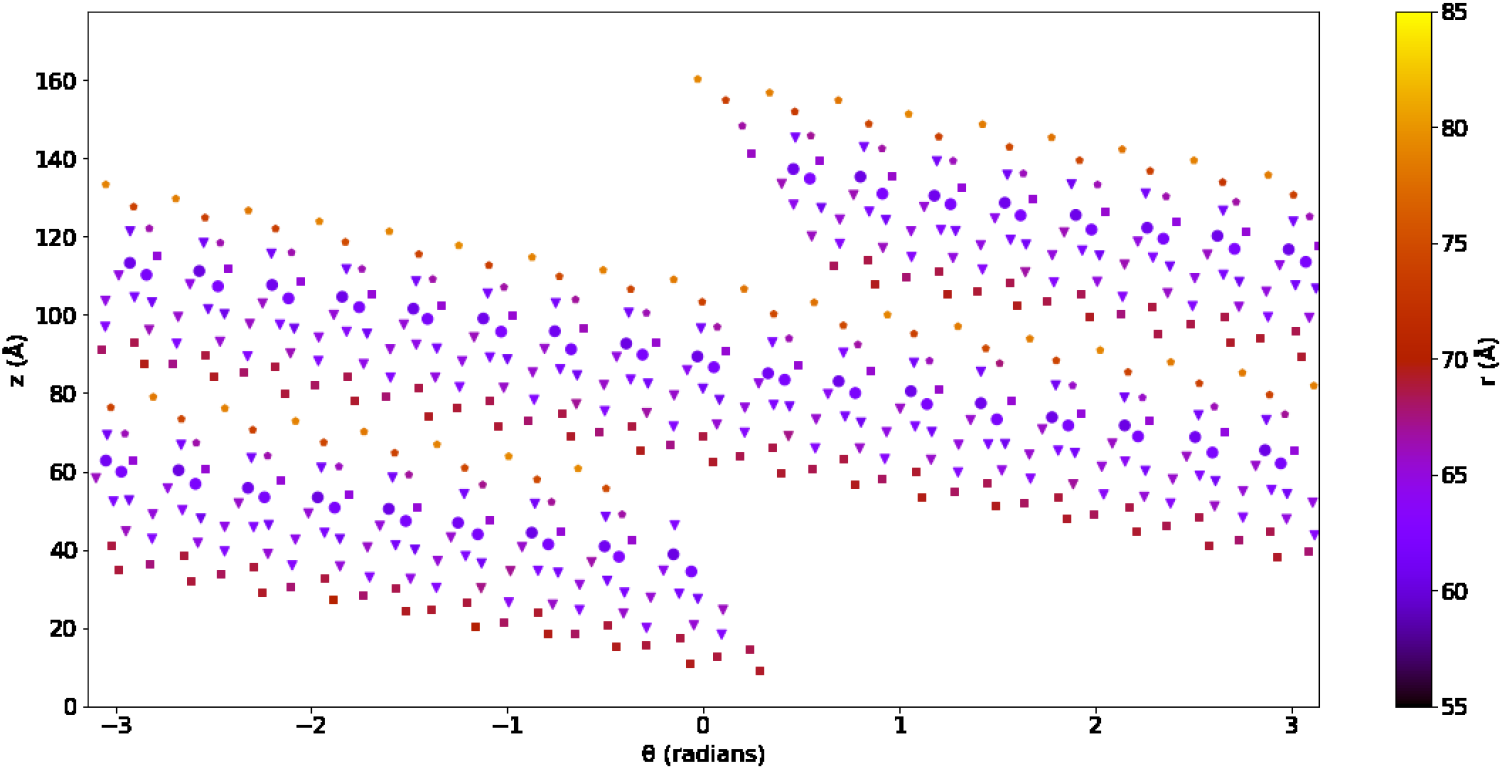
Mean positions of membrane facing CHMP1B residues represented in cylindrical coordinates. The position of the tubule axis in the xy-plane, from which cylindrical coordinates of the protein and lipid are calculated, was updated at each time step to account for subtle motions in the protein coat. The (*x*_0_, *y*_0_) position of axis of tubule was determined by performing least-squares fit on all CHMP1B F9 and F13 positions along the protein coat to place them equidistant from the tubule axis at each time step. Consistent values of *r* (indicated by color bar) for identical residues from different CHMP1B subunits indicate that the procedure results in well-fit consistent cylindrical coordinates applicable to the lipid tubule. Color variation shows that the F9+F13 contact site sits the most inward toward the tubule central axis, and elsewhere the membrane-facing protein surface shifts further out in *r*. F9 and F13 in circles; K6, K12, K16, R20, K23, K24, and K27 in triangles; K30, K35, K38, and K87 in squares; and K90, K97, and K104 in pentagons.

**Table S1.** Average number of bromine atoms per molecules, calculated from mass spectrometry (SDPC-Br, POPS-Br, and PIP2-Br) and NMR (Chol-Br) data.

| Lipid | Bromine atoms per molecule |
| --- | --- |
| SDPC-Br | 7.0 |
| CHOL-Br | 1.7 |
| POPS-Br | 1.6 |
| PIP2-Br | 2.0 |

**Table S2.**
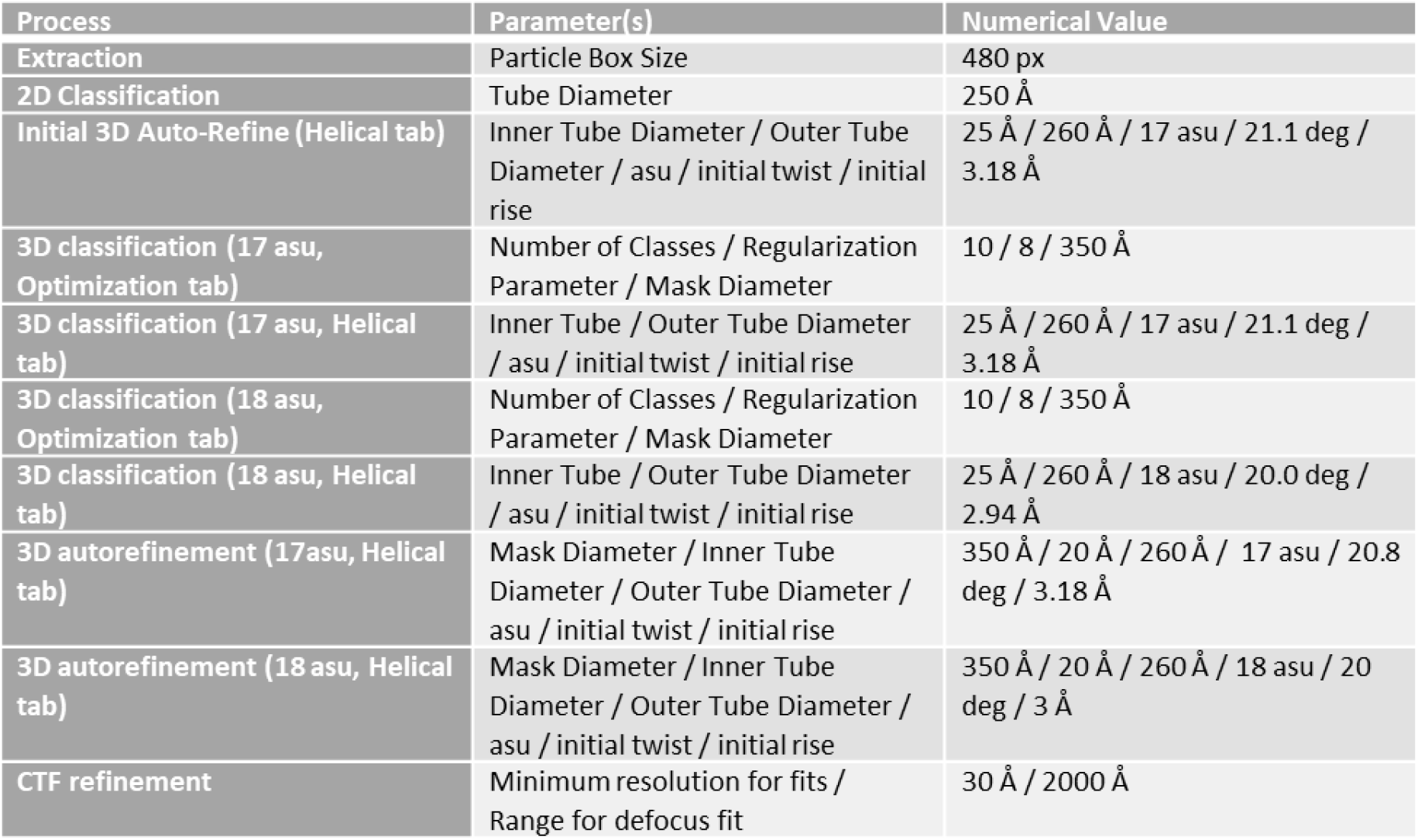
RELION data processing parameters.

**Table S3.** Krios data collection and processing statistics.

| Comp. | No Br<br>17asu | No Br<br>18asu | CHOL-Br<br>17asu | CHOL-Br<br>18asu | CHOL-I<br>17asu | CHOL-I<br>18asu | PIP2-Br<br>17asu | PIP2-Br<br>18asu | POPS-Br<br>17asu | POPS-Br<br>18asu | SDPC-Br<br>17asu | SDPC-Br<br>18asu |
| --- | --- | --- | --- | --- | --- | --- | --- | --- | --- | --- | --- | --- |
| # Mic. | 5,684 | 5,684 | 1,855 | 1,855 | 3,186 | 3,186 | 2,116 | 2,116 | 5,060 | 5,060 | 2,857 | 2,857 |
| # Initial<br>part. | 279,180 | 279,180 | 53,787 | 53,787 | 283,543 | 283,543 | 170,300 | 170,300 | 202,716 | 202,716 | 188,013 | 188,013 |
| # Final<br>part. | 26,764 | 4,103 | 12,904 | 12,868 | 63,616 | 57,773 | 17,971 | 6,698 | 45,351 | 16,746 | 30,752 | 49,713 |
| Mean<br>defocus | -1.2 | -1.3 | -1.1 | -1.1 | -1.4 | -1.4 | -1.2 | -1.2 | -1.2 | -1.2 | -1.5 | -1.5 |
| Pixel Size | 0.835 | 0.835 | 0.822 | 0.822 | 0.822 | 0.822 | 0.835 | 0.835 | 0.822 | 0.822 | 0.822 | 0.822 |
| Rise | 3.07 | 2.99 | 3.08 | 3.00 | 3.05 | 2.93 | 3.06 | 3.00 | 3.09 | 3.00 | 3.09 | 3.02 |
| Twist | -20.81 | 19.99 | -20.82 | 20.00 | -20.82 | 19.97 | -20.81 | 20.00 | -20.81 | 19.99 | -20.81 | 20.00 |
| Res. | 3.1 | 3.7 | 3.8 | 4.3 | 3.2 | 3.1 | 3.2 | 3.5 | 2.8 | 3.1 | 2.9 | 3.0 |

**Table S4.** Krios data collection and processing statistics.

| Comp. | 50%<br>SDPC-Br<br>17asu | 50%<br>SDPC-Br<br>18asu | 25%<br>SDPC-Br<br>17asu | 25%<br>SDPC-Br<br>18asu | SDPC-Br<br>F9A F13A<br>17asu | SDPC-Br<br>F9A F13A<br>18asu | SDPC-Br<br>F9E F13E<br>17asu | SDPC-Br<br>F9E F13E<br>18asu | SDPC-Br<br>F9L F13L<br>17asu | SDPC-Br<br>F9L F13L<br>18asu |
| --- | --- | --- | --- | --- | --- | --- | --- | --- | --- | --- |
| # Mic. | 3,891 | 3,891 | 2,931 | 2,931 | 1,715 | 1,717 | 1,420 | 1,420 | 2,188 | 2,188 |
| # Initial<br>part. | 256,608 | 256,608 | 285,067 | 285,067 | 251,839 | 251,839 | 65,690 | 65,690 | 186,495 | 186,495 |
| # Final<br>part. | 41,516 | 49,975 | 46,287 | 66,567 | 66,916 | 51,093 | 21,747 | 6,994 | 44,673 | 16,539 |
| Mean<br>defocus | -1.4 | -1.4 | -1.3 | -1.3 | -1.3 | -1.4 | -1.2 | -1.2 | -1.2 | -1.2 |
| Pixel<br>Size | 0.822 | 0.822 | 0.822 | 0.822 | 0.822 | 0.822 | 0.822 | 0.822 | 0.822 | 0.822 |
| Rise | 3.05 | 2.97 | 3.07 | 2.99 | 3.05 | 2.95 | 3.10 | 2.98 | 3.10 | 3.00 |
| Twist | -20.81 | 19.99 | -20.83 | 20.00 | -20.81 | 19.99 | -20.79 | 20.01 | -20.80 | 20.01 |
| Res. | 2.9 | 3.0 | 3.1 | 2.9 | 2.9 | 3.0 | 3.4 | 3.5 | 3.2 | 2.9 |

**Table S5.** Arctica data collection and processing statistics.

| Comp. | 0%<br>CHOL<br>17asu | 10%<br>CHOL<br>17asu | 20%<br>CHOL<br>17asu | 30%<br>CHOL<br>17asu | 40%<br>CHOL<br>17asu | POPC<br>Mixture<br>34asu |
| --- | --- | --- | --- | --- | --- | --- |
| # Mic. | 615 | 653 | 541 | 492 | 481 | 502 |
| # Initial<br>part. | 47,415 | 54,720 | 27,984 | 22,959 | 13,974 | 20,545 |
| # Final<br>part. | 2,772 | 3,373 | 1,685 | 1,193 | 3,677 | 2,864 |
| Mean<br>defocus | -1.6 | -1.0 | -1.3 | -1.0 | -1.0 | -1.9 |
| Pixel Size | 1.14 | 1.14 | 1.14 | 1.14 | 1.44 | 1.44 |
| Rise | -20.82 | -20.91 | -20.83 | -20.92 | -20.81 | 1.56 |
| Twist | 3.14 | 3.15 | 3.14 | 3.15 | 3.13 | 10.52 |
| Res. | 4.1 | 4.0 | 6.1 | 4.4 | 4.7 | 8.1 |

**Table S6.** Radially averaged leaflet lipid compositions. The uncertainties are the standard deviations between independent half maps.

| Lipid | Inner Leaflet<br>(mol%) | Outer Leaflet<br>(mol%) |
| --- | --- | --- |
| CHOL-Br | 28 ± 4 | 20 ± 6 |
| PIP<sub>2</sub>-Br | 17 ± 2 | 25 ± 5 |
| POPS-Br | 36 ± 6 | 33 ± 9 |
| SDPC-Br | 19 ± 2 | 22 ± 4 |

**Table S7.**
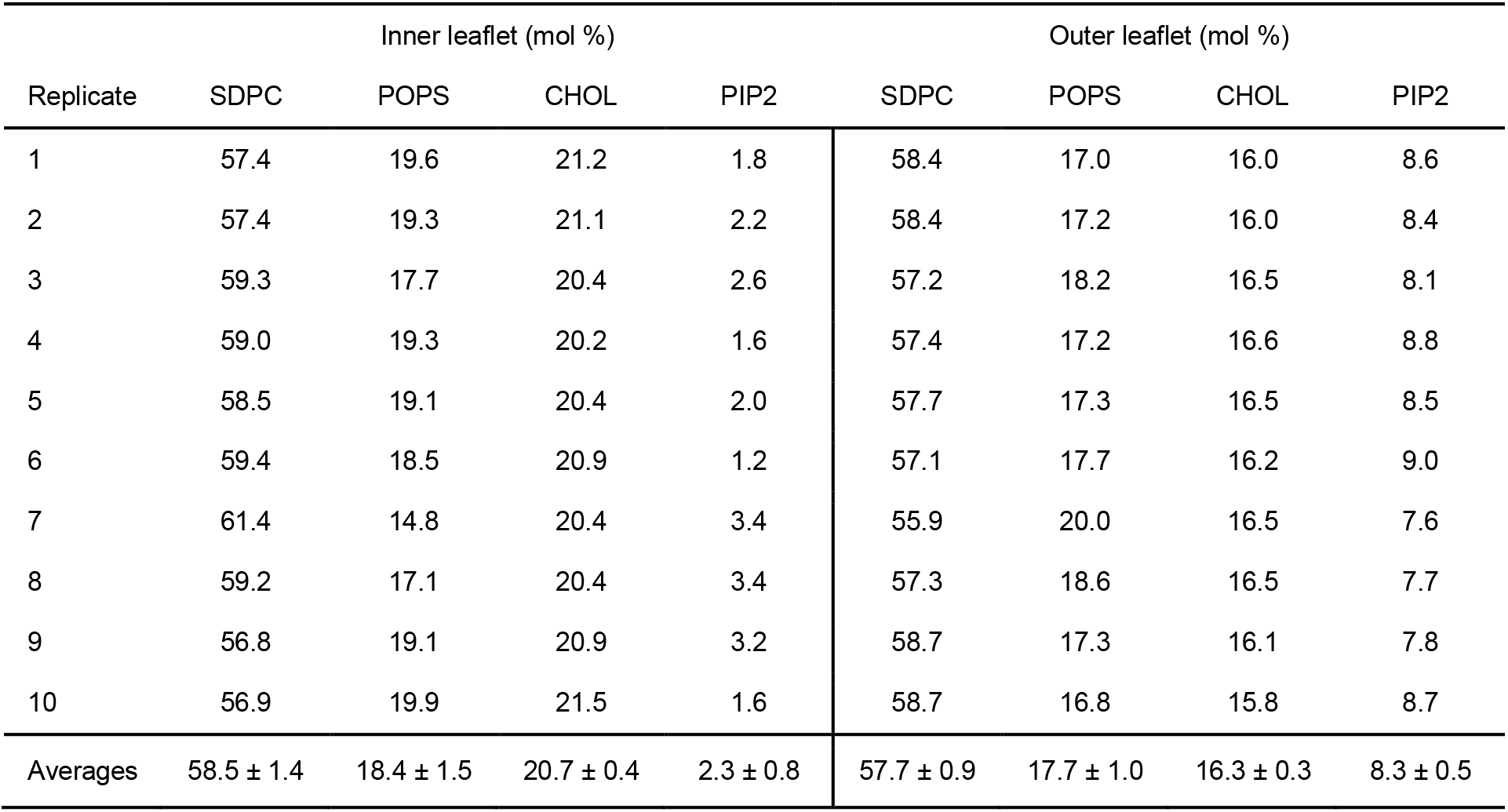
Inner and outer leaflet compositions for all 10 replicates of WT CHMP1B-IST1 production simulations corresponding to lipid composition 1 (Simulation set 1). Mean compositions and ± standard deviations of lipid mol % across 10 replicates in bottom row. The similar leaflet compositions across the ten replicates indicates that the leaflet equilibration time is sufficient.

**Table S8.** Inner and outer leaflet compositions for all 3 replicates of F9A + F13A CHMP1B-WT IST1 production simulations corresponding to lipid composition 1 (Simulation set 2). Mean compositions and ± standard deviations of lipid mol % across 3 replicates in bottom row.

| Replicate | Inner leaflet (mol %) |  |  |  | Outer leaflet (mol %) |  |  |  |
| --- | --- | --- | --- | --- | --- | --- | --- | --- |
|  | SDPC | POPS | CHOL | PIP2 | SDPC | POPS | CHOL | PIP2 |
| 1 | 56.7 | 21.1 | 21.0 | 1.2 | 58.8 | 16.1 | 16.1 | 9.0 |
| 2 | 57.4 | 19.6 | 21.4 | 1.6 | 58.4 | 17.0 | 15.9 | 8.7 |
| 3 | 58.4 | 18.6 | 20.8 | 2.2 | 57.7 | 17.6 | 16.2 | 8.4 |
| Averages | 57.5 ± 0.9 | 19.8 ± 1.3 | 21.1 ± 0.3 | 1.7 ± 0.5 | 58.3 ± 0.6 | 16.9 ± 0.8 | 16.1 ± 0.2 | 8.7 ± 0.3 |

**Table S9.** Inner and outer leaflet compositions for all 3 replicates of WT CHMP1B-IST1 production simulations with EM-resolved lipid composition 17 (Simulation set 3). Mean compositions and ± standard deviations of lipid mol % across 3 replicates in bottom row.

| Replicate | Inner leaflet (mol %) |  |  |  | Outer leaflet (mol %) |  |  |  |
| --- | --- | --- | --- | --- | --- | --- | --- | --- |
|  | SDPC | POPS | CHOL | PIP2 | SDPC | POPS | CHOL | PIP2 |
| 1 | 21.0 | 37.8 | 27.4 | 13.8 | 29.2 | 28.3 | 18.4 | 24.0 |
| 2 | 20.6 | 37.2 | 27.1 | 15.1 | 29.5 | 28.7 | 18.6 | 23.2 |
| 3 | 22.8 | 34.4 | 27.2 | 15.6 | 28.1 | 30.5 | 18.7 | 22.8 |
| Averages | 21.5 ± 1.2 | 36.5 ± 1.8 | 27.2 ± 0.2 | 14.8 ± 0.9 | 28.9 ± 0.7 | 29.2 ± 1.2 | 18.6 ± 0.2 | 23.3 ± 0.6 |

**Table S10.**
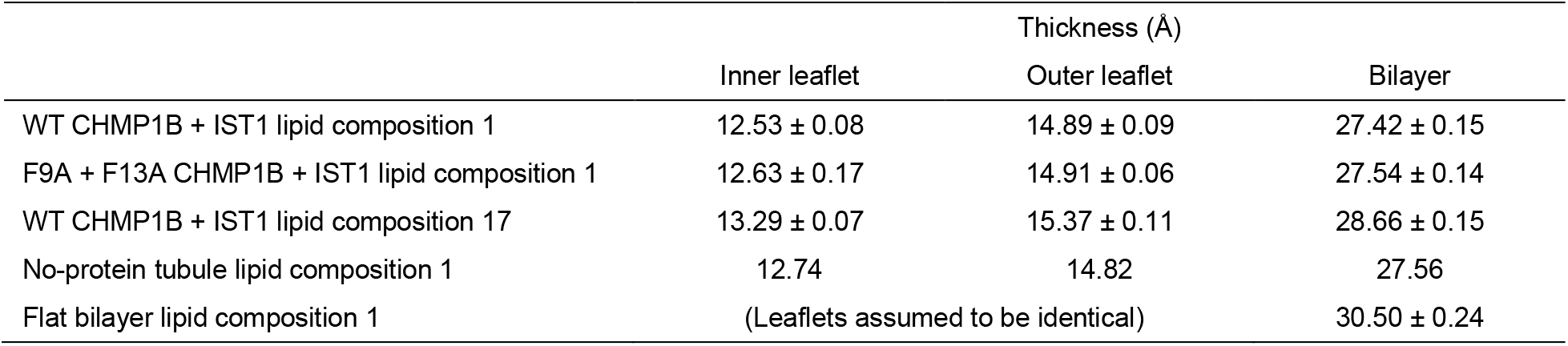
Summary of leaflet and bilayer thicknesses from all simulations. Thicknesses are based on mean positions of the second glycerol bead for all phospholipids, and for tubules the location of the bilayer midplane. Tubule calculation procedures are described in SI Methods, and tubule uncertainties are one standard deviation between calculated thicknesses of independent replicates. Full tubule structural data are in Figs. S11 (Simulation set 1, WT CHMP1B-IST1 with lipid composition 1), S16 (Simulation set 2, F9A+F13A CHMP1B-WT IST1 with lipid composition 1), S17 (Simulation set 4, no-protein tubule with lipid composition 1), and S19 (Simulation set 3, WT CHMP1B-IST1 with lipid composition 17). For the flat bilayer, a 250 x 250 Å patch was divided into a 250 x 250 grid, and for each frame a 2d cubic interpolation was applied over the grid to the positions of upper and lower leaflet second glycerol beads. A mean thickness was then calculated over the grid for each frame, and the mean over 200 frames from the last 400 ns of the simulation was calculated, with the uncertainty being the standard deviation across the 200 frames. Each of the lipid composition 1 tubules are around 10 % thinner than the composition 1 flat bilayer, with most of the thinning coming from the inner leaflet likely due to overall increased splaying of lipid tails in that leaflet. The lipid composition 17 tubules are slightly thicker, likely as a consequence of the composition being enriched in more rigid, saturated phospholipid tails relative to lipid composition 1.

|  | Thickness (Å) |  |  |
| --- | --- | --- | --- |
|  | Inner leaflet | Outer leaflet | Bilayer |
| WT CHMP1B + IST1 lipid composition 1 | 12.53 ± 0.08 | 14.89 ± 0.09 | 27.42 ± 0.15 |
| F9A + F13A CHMP1B + IST1 lipid composition 1 | 12.63 ± 0.17 | 14.91 ± 0.06 | 27.54 ± 0.14 |
| WT CHMP1B + IST1 lipid composition 17 | 13.29 ± 0.07 | 15.37 ± 0.11 | 28.66 ± 0.15 |
| No-protein tubule lipid composition 1 | 12.74 | 14.82 | 27.56 |
| Flat bilayer lipid composition 1 | (Leaflets assumed to be identical) |  | 30.50 ± 0.24 |

